# Defensive symbionts provide high protection against natural enemies at low cost to hosts: a meta-analysis

**DOI:** 10.1101/2024.11.05.622061

**Authors:** Cássia Siqueira Cesar, Eduardo S. A. Santos, Rodrigo Cogni

## Abstract

Defensive endosymbionts (i.e., endosymbionts that protect hosts against natural enemies) are widespread in terrestrial arthropods, especially insects. Protection against natural enemies may contribute to their successful spread in nature. However, the outcome of host–symbiont interactions (mutualism or parasitism) depend on the balance between the costs and benefits of the interaction and on the ecological conditions experienced by the host, such as the presence or absence of natural enemies. Here, we conducted a meta-analysis to quantitatively assess the costs and benefits of symbiont-mediated protection in terrestrial arthropods. We gathered approximately 3,425 effect sizes from 246 studies. Our results show that in the presence of natural enemies (i.e., hosts infected with pathogens or parasites), symbionts provide protection, positively affecting the fitness of hosts. In the absence of natural enemies, symbionts caused little reduction in host fitness. Overall, symbiont-mediated protection was 7.1 times stronger than the cost observed in the absence of natural enemies, indicating strong protection and little evidence of substantial fitness costs to hosts. Natural enemies attacking symbiont-infected hosts had substantial reduction in their fitness. Moreover, the level of protection and cost for both hosts and natural enemies varies between the fitness components, host families, symbiont species, and natural enemy group. Our results reveal a broad symbiont-mediated protection with little evidence of substantial costs associated, which may help explain the successful spread of symbionts in wild arthropod populations.

## 1. Introduction

Symbiotic interactions (i.e., the living together of different organisms) are ubiquitous across the life stages of most organisms and profoundly influence their ecology and evolution (Thompson, 2015; Werren et al., 2008). Symbiosis comprises a continuum of interactions ranging from harmful (i.e., parasitism) to beneficial (i.e., mutualism) (Oliver et al., 2003; Werren et al., 2008; Zug and Hammerstein, 2015). Terrestrial arthropods, particularly insects, are commonly infected with maternally transmitted facultative endosymbionts that use different strategies to infect hosts and spread within host populations (Oliver et al., 2014; Zug & Hammerstein, 2015). Endosymbionts such as *Wolbachia* and *Spiroplasma* are known for being reproductive parasites, manipulating host reproduction to increase the proportion of infected individuals within populations (Ebbert, 1991; Gotoh et al., 2007; Hunter et al., 2003; Turelli, 1994; Werren et al., 2008). For example, *Wolbachia* can induce cytoplasmic incompatibility (CI), in which crosses between infected males and uninfected females produce nonviable offspring, increasing the proportion of infected individuals in populations (Gotoh et al., 2007; Werren et al., 2008). However, reproductive parasitism alone may not be sufficient to guarantee successful invasion of a new host population. For instance, certain *Wolbachia* strains, such as *w*Mel, have incomplete maternal transmission and mild levels of CI (Fenton et al., 2011; Kriesner et al., 2013; Zug & Hammerstein, 2015). Furthermore, not all *Wolbachia* strains induce CI (Gotoh et al., 2007; Kriesner et al., 2013; Zhang et al., 2016), and many endosymbionts found in wild host populations are not reproductive parasites, as observed in aphid-associated endosymbionts such as *Hamiltonella defensa*, *Regiella insecticola*, and *Serratia symbiotica* (Guo et al., 2017). Thus, many endosymbionts rely on alternative strategies beyond reproductive parasitism to spread among and within host populations.

In recent years, studies have shown that many endosymbionts confer protection against natural enemies to their hosts (i.e., pathogens, parasites, or predators that negatively affect host fitness). Endosymbionts that provide such protection are referred to as defensive endosymbionts (hereafter, symbionts) and can protect their hosts from a wide range of natural enemies. For example, symbionts such as *Hamiltonella* and *Serratia* protect aphids against parasitoids (Oliver et al., 2003, 2014), *Regiella* protects aphids and whiteflies against parasitoids and fungal pathogens (Łukasik et al., 2013; Vorburger et al., 2010), *Spiroplasma* protects flies against parasitoids and nematodes (Jaenike et al., 2010; Xie et al., 2010), and *Wolbachia* confers protection against pathogenic bacteria in isopods, viruses in flies, and multiple pathogens in mosquitoes, including viruses, protozoans, nematodes, and pathogenic bacteria (Hedges et al., 2008; Teixeira et al., 2008) (Kambris et al., 2009, 2010; Moreira et al., 2009; van den Hurk et al., 2012; Ye et al., 2013). However, the occurrence and strength of symbiont-mediated protection vary between study systems. Most studies report symbiont-mediated protection, but a few studies report no protection against pathogens or even increased host susceptibility to natural enemies in symbiont-infected hosts (Fytrou et al., 2006; Graham et al., 2012; Longdon et al., 2012; Martinez et al., 2012; Rottschaefer & Lazzaro, 2012; Baton et al., 2013; Van Nouhuys et al., 2016). Hence, protective effects can vary among host families and may be specific to symbiont species and strains, as well as to natural enemy species and strains (Cayetano & Vorburger, 2015; Cogni et al., 2021; Martinez et al., 2014, Martinez et al., 2015; Parker et al., 2013). Consequently, the outcome of host-symbiont interactions - parasitism or mutualism - depends on the symbiont species and strains, on the balance between costs and benefits of symbiont infection, and on the ecological conditions experienced by the host.

Symbiont infection can be costly for hosts, but these costs can be overcome in the presence of natural enemies, shifting the interaction toward facultative mutualism (Zug & Hammerstein, 2015). In this context, symbionts provide fitness benefits to hosts, such as increased survival or fecundity, which may facilitate the infection and spread of symbionts among and within host populations (Werren et al., 2008; Zug & Hammerstein, 2015; Zytynska et al., 2021). In contrast, in the absence of natural enemies, the interaction can be costly for hosts, shifting towards parasitism, where only the symbiont benefits (Martinez et al., 2014; Oliver et al., 2014; Werren et al., 2008; Zug & Hammerstein, 2015). For example, under parasitism, *Hamiltonella*-infected aphids gain direct fitness benefits, such as increased survival, whereas in the absence of parasitism, *Hamiltonella* infection reduces host survival and fecundity, depending on the symbiont strain (Vorburger et al., 2013; Martinez et al., 2018). Similarly, some *Wolbachia* strains provide strong antiviral protection by reducing viral titers in *Drosophila*, but in the absence of viruses, these strains reduce host fecundity and survival (Martinez et al., 2014; Martinez et al., 2015; Pimentel et al., 2025). However, in some cases, symbionts impose only mild or no fitness costs on their hosts (Martinez et al., 2015; Vorburger et al., 2009). Therefore, in most cases, symbionts are expected to reduce host fitness in the absence of natural enemies but increase host fitness and reduce enemy fitness when natural enemies are present.

Variation in the costs and benefits of symbiont infection is often associated with differences between symbiont strains (Martinez et al., 2014, 2015; Martinez et al., 2018). For example, the level of protection against parasitoids and is associated costs varies between *Hamiltonella* strains. This variation can be explained by the association of *Hamiltonella* with bacteriophages called APSEs *(Acyrthosiphon pisum* secondary endosymbiont), which encode putative toxins that are responsible for defense against parasitoids. Different APSEs confer different levels of protection. For instance, *Hd*-APSE3 confers higher toxicity and faster parasitoid mortality in comparison to other strains (Doremus et al., 2018; Martinez et al., 2018; Vorburger & Gouskov, 2011). To test this variation, studies have transinfected (i.e., introduced) *Hamiltonella* strains carrying different APSEs into different aphid clones (Martinez et al., 2014; Martinez et al., 2018). Similarly, in *Drosophila*, *Wolbachia* strains such as *w*Mel and *w*MelCS provide strong antiviral protection by substantially reducing viral titers in comparison to other strains (Martinez et al., 2014, Martinez et al., 2015; Pimentel et al., 2025). Because of their potential to reduce viral infection, these strains are often transinfected into other hosts, such as mosquitoes from genera *Aedes* and *Culex,* to control insect-borne human diseases such as dengue, chikungunya and malaria (Bian et al., 2010; Hughes et al., 2011; Moreira et al., 2009; van den Hurk et al., 2012; Werren et al., 2008; Xue et al., 2018). In these cases, variation in the level of protection is associated with *Wolbachia* density within host cells – higher *Wolbachia* densities provide stronger protection but impose higher fitness costs in the absence of viruses (Martinez et al., 2014; Martinez et al., 2015). Therefore, these findings suggest that introduced symbiont strains tend to confer stronger protection but also higher fitness costs, whereas natural symbiont strains provide lower protection with lower associated costs.

Although over the past two decades symbiont-mediated protection and its associated costs have been extensively quantified within individual systems, particularly *Drosophila*/mosquito–*Wolbachia*– virus and aphid–*Hamiltonella*–parasitoid, the overall magnitude of the costs and benefits of host– symbiont associations across terrestrial arthropod systems remains unclear. Here, we conducted a meta-analysis to quantitatively assess the costs and benefits of endosymbiont-mediated protection in terrestrial arthropods. Specifically, we addressed the following questions: (1) what is the magnitude of the protective effect conferred by symbionts to their hosts? (2) what is the magnitude of the costs of symbiont infection for hosts? and (3) what is the magnitude of the costs of symbiont infection for natural enemies? Furthermore, we tested possible sources of variation in symbiont-mediated effects across study systems, such as host family, symbiont origin (i.e., natural or introduced into hosts), fitness components, symbiont species, symbiont number, natural enemy group, and natural enemy killing strategy.

## 2. Methods

### 2.1. Literature search

We conducted a literature search using all databases of ISI *Web of Science* and *Scopus* platforms, and Google Scholar. The search was last updated in January 2026. We used the following combinations of keywords: “(protect* OR defens*) AND (symbio*) AND (fitness)”, “(defenc*) AND (symbio*) AND (fitness)”, and “(endosymbio*) AND (fitness)”. We also searched for grey literature at the advanced search (subject areas: “evolutionary biology” and “ecology”) in the *bioRxiv* platform using the same keywords. Additionally, we conducted a backward search of studies cited in the reference list of the following review articles: Oliver et al. (2014), Vorburger & Perlman (2018) and Zug & Hammerstein (2015). This allowed us to identify 45 additional studies.

### 2.2. Inclusion criteria

For the title and abstract screening, studies must have: (1) been performed on arthropod hosts, (2) mentioned that they measured the fitness of hosts and/or natural enemies, and (3) mentioned the presence of one or more defensive symbionts infecting hosts. For the full text screening, studies must have: (1) been empirical, (2) been performed in whole organisms (cell culture/in vitro experiments were excluded), (3) measured fecundity, egg hatch, longevity, survival or any other fitness measure (i.e., response variable) of the host in the presence and/or absence of a natural enemy, (4) measured viral load, parasitoid survival or any other fitness measure, which varies according to the natural enemy, (5) compared symbiont-infected hosts (treatment group) and symbiont-uninfected hosts (control group). Symbiont-uninfected hosts could be either naturally uninfected or treated with antibiotics, (6) reported sample size, mean and standard deviation (SD) or standard error (SE), or proportion/number of events of the response variables. If descriptive data and graphs were missing, study must have reported inferential statistics (F, t, and degree of freedom) to be included; (7) the symbiont in the study had to be defensive. We considered a symbiont to be defensive when at least one study demonstrated that it conferred a fitness benefit to the host in the presence of a natural enemy. However, the great majority of our datasets consists of well-established study systems (Figure 1, Tables S1, S2, and S3), and (8) the defensive symbiont had to be an endosymbiont. Criteria (3) and (4) did not have to be attended at the same time because we had distinct datasets for host and natural enemy data. After full text screening, our sample included 246 studies. The full screening process is summarized in a PRISMA diagram (Moher et al., 2009) (Figure S1). A list with all studies included in our meta-analyses is provided in the Supplementary information.

**Figure 1.**
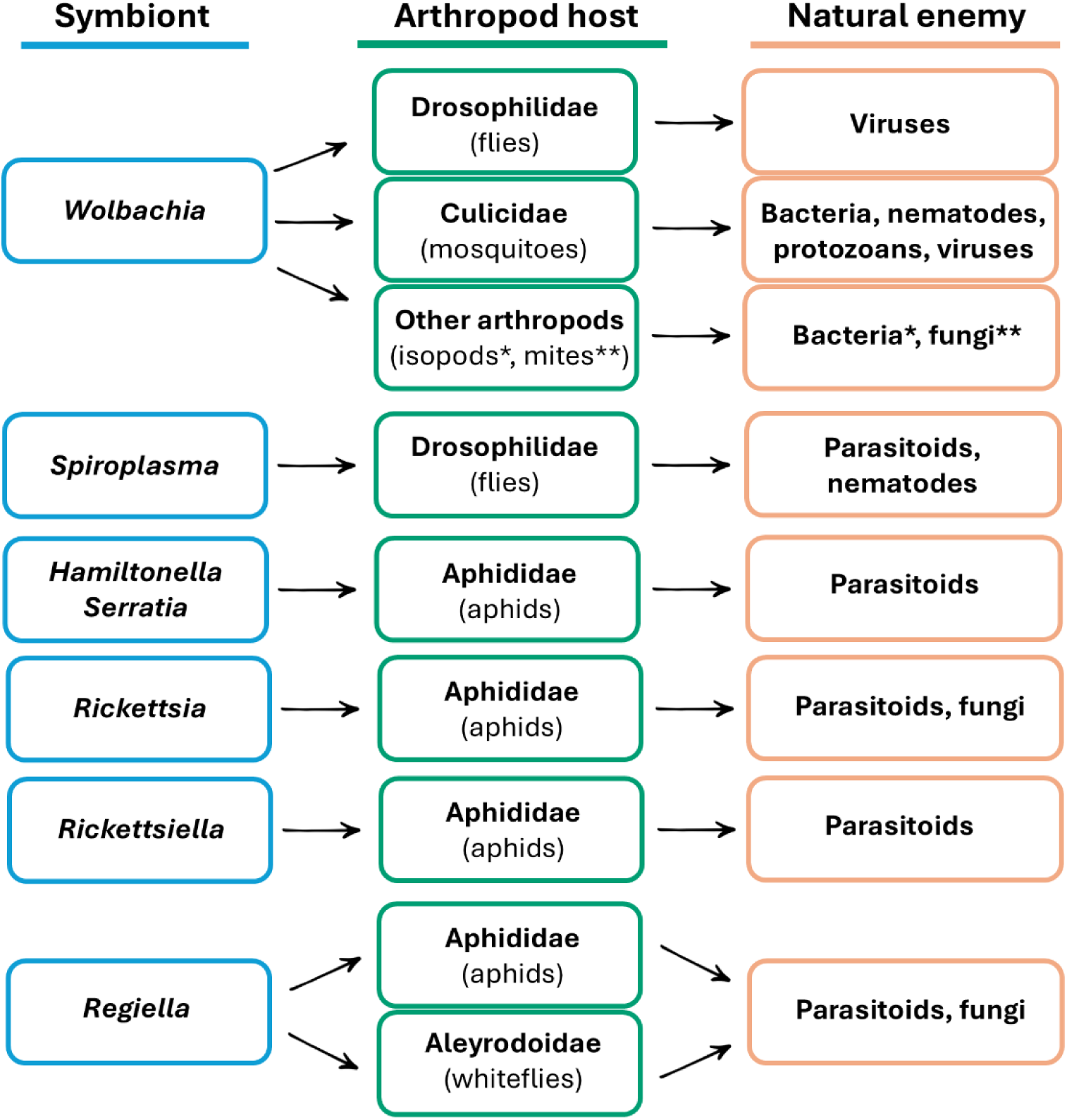
Representative study systems frequently included in this meta-analysis and widely recognized in the literature. The complete list of study systems is provided in Tables S1, S2, and S3.

### 2.3. Data extraction

We extracted different fitness measures and grouped similar data under six broad categories of fitness measure: survival, fecundity, body size, development time, number of uninfected hosts (i.e., the percentage or number of individuals that were not successfully parasitized or infected by natural enemies, used here as a proxy for host fitness, as hosts free of natural enemy infections tend to have higher survival), and natural enemy load (i.e., the abundance of natural enemy replicating inside the host, such as viruses and bacteria) (Table S4). Survival, fecundity, body size, development time, and number of uninfected hosts were used as measures of host fitness. Body size, survival, and natural enemy load were used as measures of natural enemy fitness.

We extracted raw data from tables and figures when available. Data presented in figures were extracted using *PlotDigitizer* (Huwaldt, 2005). When raw data were not available, we extracted summary statistics from inferential tests (e.g., F, t, and degrees of freedom). For studies reporting mean ± SE, we converted SE to SD. When neither SD nor SE was reported, we estimated SD using the metric proposed by Bracken (1992) (Lajeunesse, 2013). For studies reporting results across multiple generations, we extracted data only from the first generation, as it was a restricted number of studies and our analyses did not aim to assess temporal effects (i.e., repeated measures). When the figure reported was a survival curve, we extracted the median and calculated the estimated SD using the Bracken (1992) metric (Lajeunesse, 2013). We considered the median equal to the mean when the sample size was ≥ 25 (Hozo et al., 2005).

We organized the data into three distinct datasets: (1) protection, which included fitness measures of hosts with and without symbionts when exposed to natural enemies; (2) cost to hosts, which included fitness measures of hosts with and without symbionts in the absence of natural enemies; and (3) cost to natural enemies, which included fitness measures of natural enemies after infecting or parasitizing hosts with and without symbionts. In addition to the fitness measures used to calculate effect sizes, we collected characteristics of the original studies that could potentially account for variation in host–symbiont interactions, which we included as moderators in the models: (1) Host family: taxonomic family to which hosts belong to; (2) Fitness measure: the six broad categories of fitness measure (body size, development time, fecundity, survival, number of uninfected hosts, and natural enemy load); (3) Symbiont type: whether the symbiont is naturally associated with the host species or was introduced (i.e., artificially infected into a non-native species); (4) Symbiont species; (5) Symbiont number: whether the host was infected by a single symbiont species or coinfected by two symbiont species; (6) Natural enemy group: type of natural enemy (bacterium, fungus, nematode, parasitoid, protozoan, or virus); (7) Natural enemy killing strategy: whether the natural enemy is obligate lethal (i.e., dependent on host death for successful infection) or facultative lethal (Table S5). We excluded studies with predators as natural enemies because, in this specific scenario, symbionts do not confer a direct fitness benefit to their hosts. Instead, the fitness of the natural enemy is negatively affected only after the predator consumes a symbiont-infected host and becomes infected itself (Noriyuki et al., 2014).

### 2.4. Effect size calculation

For data reported as proportion or numbers of events, we used the odds ratio (OR) to calculate effect sizes, comparing the odds of survival between symbiont-infected hosts (treatment group) and symbiont-uninfected hosts (control group). For data reported as mean ± SD, we calculated effect sizes using Hedge’s g, which is the standardized mean difference between treatment and control groups corrected for bias (Freeman et al., 1986). Thus, Hedges’ g quantified the mean difference in fitness between symbiont-infected and symbiont-uninfected hosts. In the case of natural enemies, we compared the odds of survival and differences in mean fitness between natural enemies that attacked symbiont-infected and symbiont-uninfected hosts. We converted OR estimates to Hedges’ g to obtain a common metric (Polanin and Snilstveit, 2016). The calculation of effect sizes, including the conversion of OR to Hedges’ g, was performed using the function *escalc* in the *metafor* package in R (R Core Team, 2026; Viechtbauer, 2017). We calculated and converted the data collected from summary statistics to Hedges’ g using the Practical Meta-Analysis Effect Size Calculator, which uses pre-established equations to estimate effect sizes from inferential tests (e.g., ANOVA and t-tests) (Wilson, 2014). Therefore, all data types collected were converted into the same metric (Hedges’ g), making them comparable. We calculated and adjusted the direction of effect sizes such that positive values of Hedges’ g indicate that symbionts have a positive effect on the fitness of either the host or the natural enemy. Hence, positive values mean higher fecundity, survival, body size, number of uninfected hosts, and natural enemy load, but shorter development time (i.e., faster development). Conversely, negative values indicate that symbionts have a negative effect on fitness.

### 2.5. Statistical analysis

We divided our analyses into two parts: the main analyses, which included all study groups together, and separate analyses for the two most represented subgroups in the dataset: (1) aphid hosts and (2) hosts harboring *Wolbachia* as a symbiont.

#### 2.5.1. Main analysis

##### Multilevel meta-analytical models

We used multilevel meta-analytic models fitted with restricted maximum likelihood (REML) estimation. Because effect sizes were not independent (i.e., multiple effect sizes originated from the same study), we included several random effects in the models: study ID (identity of each study), effect size ID (identity of each effect size), host species, and host phylogeny, which was incorporated into the models with a covariance matrix. This structure allowed us to account for the between-study variance (i.e., the variation between studies), the within-study variance (i.e., the variation between effect sizes obtained from the same study), the species-specific variance, and the phylogenetic variance. We built phylogenetic trees containing the host species from each of our datasets using the Interactive Tree of Life online tree generator (iTOL) (http://itol.embl.de/) (Letunic and Bork, 2024). We used only the topology of the trees in our meta-analyses. We were unable to include symbiont phylogeny because some studies involved symbiont coinfections, and excluding these data would substantially reduce the scope and strength of our analyses. Furthermore, we did not account for natural enemy phylogeny due to the lack of comprehensive phylogenetic information for viruses.

All meta-analytical models were fitted using the *rma.mv* function in *metafor* package in R (R Core Team, 2026; Viechtbauer, 2017). To investigate the effects of symbiont infection on host fitness, we first fitted overall random-effects models (intercept only) including all effect sizes. We then fitted separate single-predictor models (without intercepts), in which each moderator (host family, fitness measure, symbiont type, symbiont species, symbiont number, natural enemy group, and natural enemy killing strategy) was included as a fixed effect alongside the random effects to estimate parameter values for each moderator level (Table S6). Because at least 10 effect sizes are recommended for each moderator level (Nakagawa et al., 2017), moderator levels with less than 10 effect sizes were excluded from the model. The only exception was the “nematode” level (k = 9; Table S10). For the “host family” and “symbiont species” models, levels with less than 10 effect sizes were grouped as “other families” and “other species”, respectively.

Additionally, to test whether higher levels of protection by symbionts were associated with higher fitness costs to hosts, we used only studies that simultaneously quantified both outcomes (i.e., protection and cost to hosts) in the same host–symbiont systems. We aggregated multiple effect sizes (i.e., fitness measures) into a single effect size per host–symbiont system. We then performed a multilevel meta-regression using the *rma.mv* function in the *metafor* package in R (R Core Team, 2026; Viechtbauer, 2017). We accounted for the non-independence among effect sizes by including study ID, host species, and symbiont species as random factors. We did not include host phylogeny as a random effect because a likelihood ratio test comparing models with and without phylogenetic correlation showed no improvement in model fit (LRT = 0, *p* = 1.00). After conducting a sensitivity analysis (described below) we identified two influential points and ran the analysis with and without the two outliers.

##### Heterogeneity and publication bias

For each model, we quantified total heterogeneity (I^2^) and estimated the proportion of variance. explained by each random effect with the *orchaRd* package in R (R Core Team, 2026).

Additionally, we estimated the potential publication bias for each model. Publication bias occurs when the outcomes reported in the published literature deviate from those observed in all conducted studies and statistical analyses, leading to an incomplete and potentially skewed representation of the available evidence, which can lead to erroneous conclusions (Jennions et al., 2013). We checked for potential publication bias using Egger’s regression (Egger et al., 1997) with the *lm* function in R (R Core Team, 2026), using the residuals from the multilevel meta-analytic models.

##### Sensitivity analysis

To evaluate the influence of individual data points on model estimates, we calculated Cook’s distance (*D*) using the *cooks.distance.rma.mv* function from the *metafor* package (Viechtbauer, 2010). Data points are considered influential when *D* > 1 (Cook & Weisberg, 1982). To evaluate how parameter estimates changed in the presence and absence of influential observations, we calculated DFBETAS using the *dfbetas.rma.mv* function from the *metafor* package (Viechtbauer, 2010). Data points are considered influential when DFBETAS > 1 (Viechtbauer & Cheung, 2010). We ran this analysis for our overall models.

#### 2.5.2. Subgroup analysis

We followed the same methodology and criteria for the subgroup analyses “*Wolbachia*” (i.e., analysis including only datapoints of hosts infected with *Wolbachia*) and “Aphids” (i.e., analysis including only datapoints from hosts belonging to family Aphididae). After fitting the overall models only with random effects, we fitted single-predictor models including the same moderators described above as fixed effects when possible (Tables S7 and S8). We also quantified total heterogeneity, assessed potential publication bias for each model, and ran a sensitivity analysis for overall models as described above.

## 3. Results

### 3.1. Main analysis

#### Protection: effect of symbiont infection on host fitness in the presence of natural enemies

All results presented in this section are specific to contexts with the presence of natural enemies. Mean overall effect size was moderate to high and positive, indicating that symbionts offer high protection against natural enemies, increasing host fitness (Table 1, Figure 2a). Fitness of hosts from Aphididae and Drosophilidae families was increased by symbiont infection, but no significant effect caused by symbiont infection was detected in hosts from Culicidae and other arthropod families (Table 1, Figure 2b). However, interpretation is limited by the small number of effect sizes collected from other arthropod families (k = 20). While symbiont-infected hosts had higher survival and a higher proportion of individuals not infected or not parasitized by natural enemies (uninfected hosts), symbiont infection had no apparent effect on host fecundity (Table 1, Figure 2c). However, these results should be interpreted with caution due to the relatively small number of effect sizes from fecundity group (k = 34). Both natural and introduced symbionts positively affected host fitness (Table 1, Figure 2d), indicating that both types of symbionts offer protection. All symbiont species showed evidence of conferring protection against natural enemies, except for *Regiella* and *Spiroplasma* simultaneously infecting hosts (Table 1, Figure 2e).

**Figure 2.**
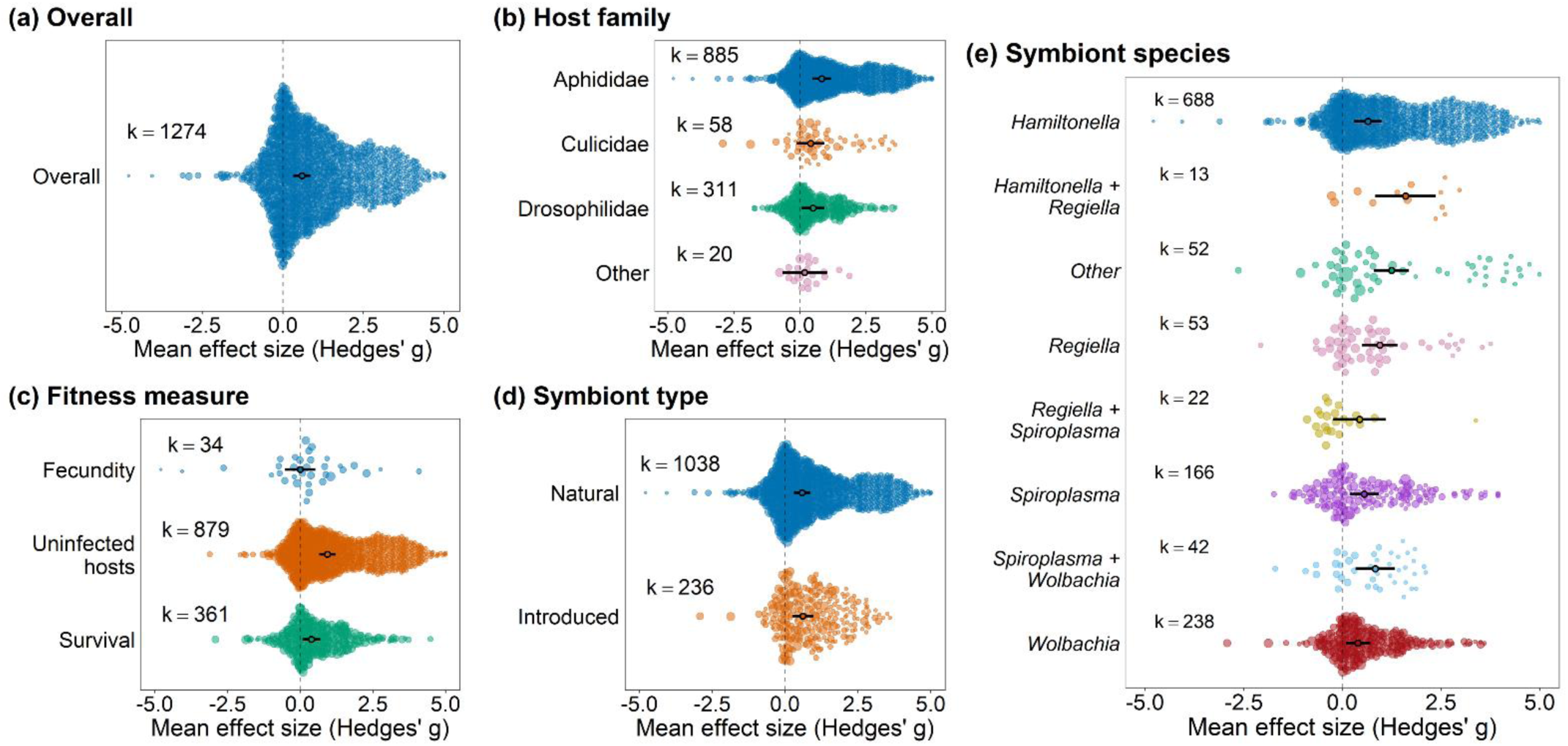
Effect of defensive symbionts on the fitness of hosts in the presence of natural enemies. (a) overall model, (b) model including host family as moderator, (c) model including fitness measure as moderator, (d) model including symbiont type (i.e., if the symbiont naturally infects the host species or if it was artificially introduced) as moderator, and (e) model including symbiont species as moderator. Positive values indicate positive effects on host fitness. Hence, positive values indicate higher fecundity, survival, and proportion of hosts that were uninfected (i.e., uninfected hosts - number of individuals that were unsuccessfully parasitized or infected by natural enemies). Negative values indicate that symbionts have a negative effect on fitness. Points are the weighted mean effect sizes ± 95% confidence intervals. Each colored circle represents an effect size, scaled according to precision (inverse of the sampling error variance). ‘k’ is the number of effect sizes.

**Table 1.**
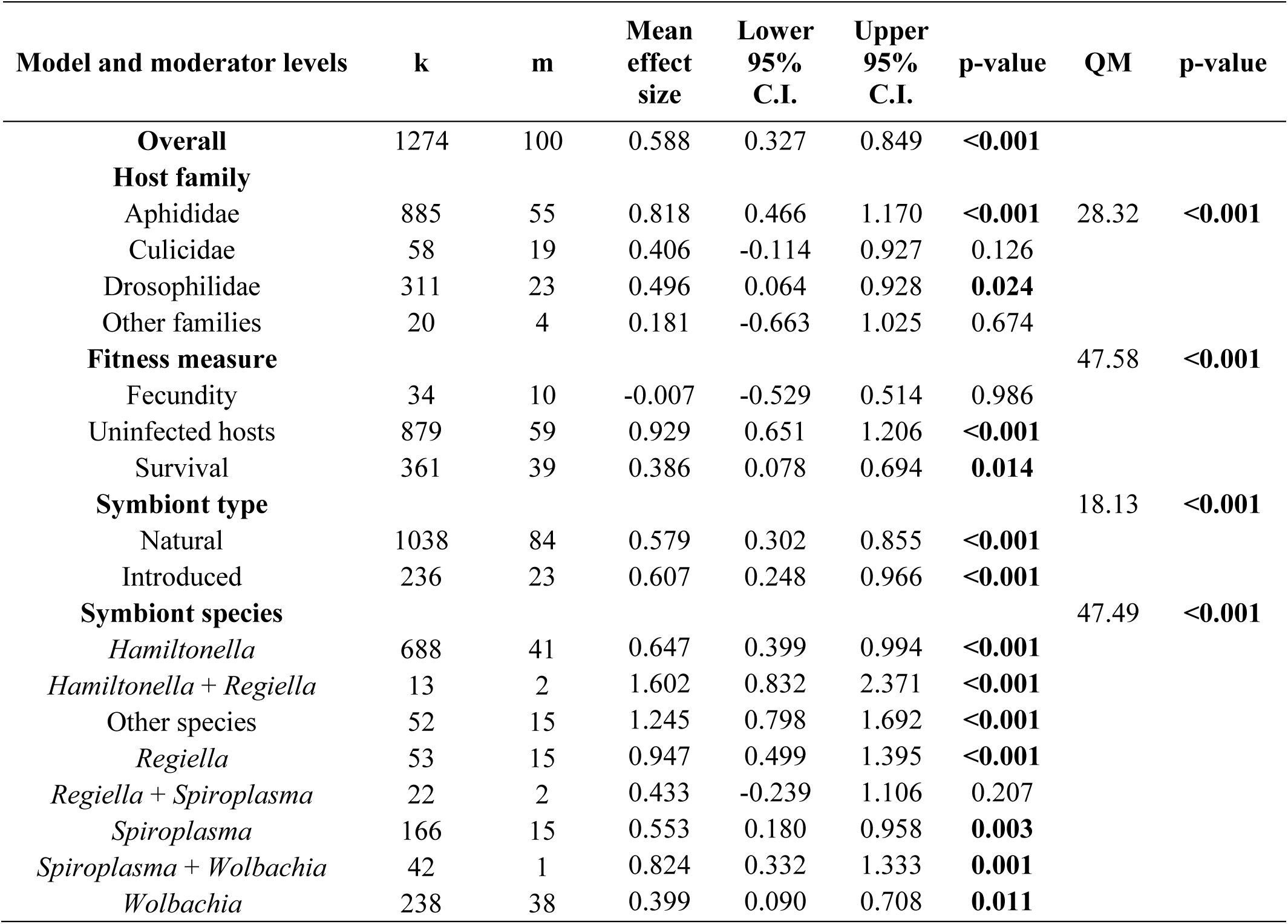
Results of the meta-analysis on the protective effect of symbionts on arthropod hosts in the presence of natural enemies and test of moderators (QM), including the overall model and the models including the following moderators: host family, fitness measure, symbiont type, and symbiont species. ‘k’ is the number of effect sizes and ‘m’ is the number of studies. p-values that are statistically significant are in bold (α =0.05).

Both single infections and coinfections with symbionts improved host fitness, but the fitness increase in coinfected hosts was 1.73 times greater than in single-infected hosts (Table S9, Figure S2a). Symbionts protected hosts against viruses, parasitoids and fungi, but no significant effect was detected on hosts infected with bacteria, nematodes, and protozoans (Table S9, Figure S2b). However, the latter three groups had relatively small sample sizes (k = 20, 17, and 14, respectively). Symbionts protected hosts against both facultative and lethal natural enemies (Table S9, Figure S2c).

#### Cost to hosts: effect of symbiont infection on host fitness in the absence of natural enemies

All results presented in this section are specific to contexts with the absence of natural enemies. Mean overall effect size was negative, small, and non-significant, providing little evidence that symbionts impose significant fitness costs on hosts in the absence of natural enemies. Most effect sizes were clustered close to zero, although both positive and negative effects were observed across studies (Table 2, Figure 3a), indicating that the effects of symbiont infection vary across study systems. Hosts from the family Aphididae had a negative effect on their fitness because of symbiont infection, but no significant effect caused by symbiont infection was detected in hosts from families Culicidae, Drosophilidae, and other arthropod families (Table 2, Figure 3b). Body size and development time were positively affected by symbiont infection, while survival was negatively affected, and no significant effect on fecundity was detected (Table 2, Figure 3c). Both natural and introduced symbionts showed no evidence of negative effects on host fitness (Table 2, Figure 3d). Only *Hamiltonella* and the coinfection of *Hamiltonella* and *Rickettsiella* significantly reduced host fitness (Table 2, Figure 3e). However, these results should be interpreted with caution due to the small number of effect sizes from *Hamiltonella* and *Rickettsiella* group (k = 10). No significant effect on host fitness was detected for either single infections or coinfections (Table S9, Figure S3).

**Figure 3.**
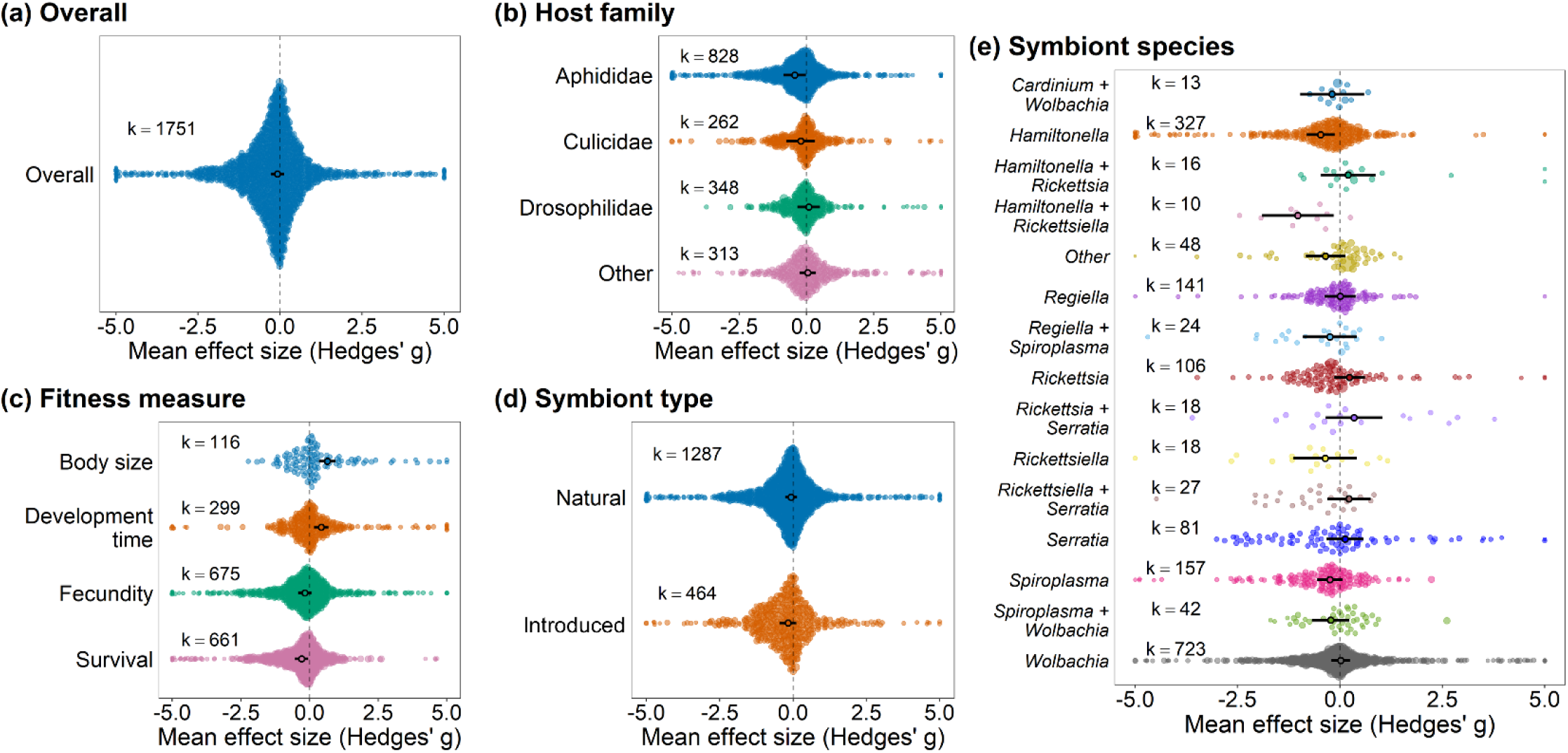
Effect of defensive symbionts on the fitness of hosts that are in the absence of natural enemies. (a) overall model, (b) model including host family as moderator, (c) model including fitness measure as moderator, (d) model including symbiont type (i.e., if the symbiont naturally infects the host species or if it was artificially introduced) as moderator, and (e) model including symbiont species as moderator. Positive values indicate positive effects on host fitness. Hence, positive values indicate higher fecundity, survival, body size and shorter development time. Negative values indicate that symbionts have a negative effect on fitness. Points are the weighted mean effect sizes ± 95% confidence intervals. Each colored circle represents an effect size, scaled according to precision (inverse of the sampling error variance). ‘k’ is the number of effect sizes. Sixteen points fell below −5 and 13 points fell above +5, which expanded the x-axis range and compressed the visual spacing of the remaining points. For visualization purposes only, these points are shown at ±5. All data points were included in the statistical analyses.

**Table 2.** Results of the meta-analysis on the costs of symbiont-infection on arthropod hosts in the absence of natural enemies and test of moderators (QM), including the overall model and the models including the following moderators: host family, fitness measure, symbiont type, and symbiont species. ‘k’ is the number of effect sizes and ‘m’ is the number of studies. p-values that are statistically significant are in bold (α =0.05).

| Model and moderator levels | k | m | Mean<br>effect<br>size | Lower<br>95%<br>C.I. | Upper<br>95%<br>C.I. | p-value | QM | p-value |
| --- | --- | --- | --- | --- | --- | --- | --- | --- |
| **Overall** | 1751 | 174 | -0.083 | -0.285 | 0.119 | 0.420 |  |  |
| **Host family** |  |  |  |  |  |  | 5.19 | 0.269 |
| Aphididae | 828 | 53 | -0.437 | -0.850 | -0.024 | **0.038** |  |  |
| Culicidae | 262 | 29 | -0.211 | -0.740 | 0.319 | 0.436 |  |  |
| Drosophilidae | 348 | 42 | 0.087 | -0.332 | 0.506 | 0.685 |  |  |
| Other families | 313 | 50 | 0.052 | -0.250 | 0.354 | 0.735 |  |  |
| **Fitness measure** |  |  |  |  |  |  | 114.38 | **<0.001** |
| Body size | 116 | 31 | 0.647 | 0.338 | 0.956 | **<0.001** |  |  |
| Development time | 299 | 60 | 0.429 | 0.170 | 0.689 | **0.001** |  |  |
| Fecundity | 675 | 144 | -0.172 | -0.411 | 0.067 | 0.159 |  |  |
| Survival | 661 | 114 | -0.287 | -0.527 | -0.048 | **0.019** |  |  |
| **Symbiont type** |  |  |  |  |  |  | 1.360 | 0.507 |
| Natural | 1287 | 144 | -0.066 | -0.270 | 0.139 | 0.529 |  |  |
| Introduced | 464 | 46 | -0.166 | -0.446 | 0.113 | 0.244 |  |  |
| **Symbiont species** |  |  |  |  |  |  | 34.74 | **0.003** |
| *Cardinium + Wolbachia* | 13 | 4 | -0.192 | -0.972 | 0.588 | 0.630 |  |  |
| *Hamiltonella* | 327 | 35 | -0.469 | -0.809 | -0.129 | **0.007** |  |  |
| *Hamiltonella + Rickettsia* | 16 | 3 | 0.200 | -0.467 | 0.867 | 0.556 |  |  |
| *Hamiltonella + Rickettsiella* | 10 | 2 | -1.029 | -1.904 | -0.154 | **0.021** |  |  |
| Other species | 48 | 9 | -0.349 | -0.829 | 0.130 | 0.153 |  |  |
| *Regiella* | 141 | 20 | 0.009 | -0.370 | 0.389 | 0.962 |  |  |
| *Regiella + Spiroplasma* | 24 | 3 | -0.246 | -0.900 | 0.408 | 0.461 |  |  |
| *Rickettsia* | 106 | 13 | 0.234 | -0.141 | 0.609 | 0.222 |  |  |
| *Rickettsia + Serratia* | 18 | 1 | 0.347 | -0.346 | 1.040 | 0.326 |  |  |
| *Rickettsiella* | 18 | 3 | -0.361 | -1.132 | 0.411 | 0.360 |  |  |
| *Rickettsiella + Serratia* | 27 | 1 | 0.218 | -0.311 | 0.747 | 0.420 |  |  |
| *Serratia* | 81 | 12 | 0.128 | -0.322 | 0.579 | 0.576 |  |  |
| *Spiroplasma* | 157 | 24 | -0.247 | -0.554 | 0.061 | 0.116 |  |  |
| *Spiroplasma + Wolbachia* | 42 | 9 | -0.225 | -0.686 | 0.235 | 0.338 |  |  |
| *Wolbachia* | 723 | 96 | 0.019 | -0.213 | 0.251 | 0.871 |  |  |

#### Trade-off between symbiont protection and fitness cost to hosts

When all effect sizes were included, we found a weak negative association between protective effects and host costs, although this relationship was only marginally significant (β = −0.064, 95% C.I. = −0.128 to 0.0001, *p* = 0.0504; Figure 4a). After removing the two influential observations identified by Cook’s distance sensitivity analysis (*D* > 1), the relationship was no longer apparent (β = −0.024, 95% C.I. = −0.085 to 0.036, *p* = 0.43; Figure 4b). Thus, the estimated negative association between protection and host costs was sensitive to a small number of influential observations, providing no robust evidence for a general protection–cost trade-off across host–symbiont systems.

**Figure 4.**
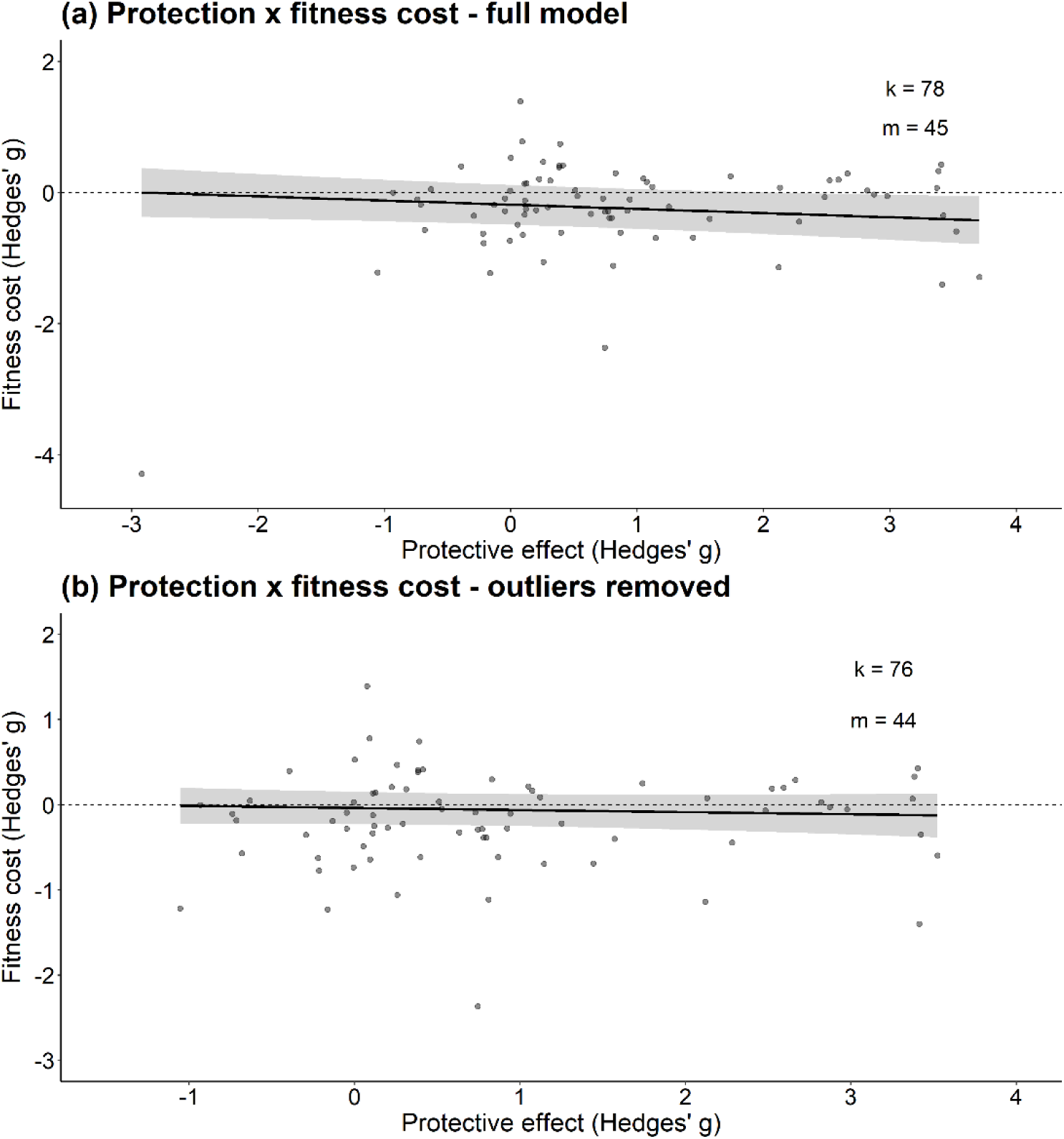
Relationship between protective effects and host fitness costs in different host–symbiont systems. (a) model including all effect sizes, and (b) model with the two influential observations removed. Each point represents an aggregated effect size (i.e., multiple fitness measures) for a given host–symbiont combination from studies that simultaneously measured protection (effect of symbiont infection on host fitness in the presence of natural enemies) and cost (effect of symbiont infection on host fitness in the absence of natural enemies). The meta-regression line is shown in black, and the 95% confidence interval is the grey area around the line. ‘k’ is the number of effect sizes and ‘m’ is the number of studies.

#### Cost to natural enemies: effect of symbionts on the fitness of natural enemies attacking symbiont-infected hosts

Mean overall effect size was negative and strong, indicating that natural enemies had a substantial reduction in overall fitness when attacking symbiont-infected hosts (Table 3, Figure 5a). No significant effect was detected in natural enemies attacking symbiont-infected hosts from families Aphididae, Culicidae, Drosophilidae, and other arthropod families (Table 3, Figure 5b). The lack of significant specific host family effects may reflect the phylogenetic relatedness among host species, as host phylogeny accounted for part of the residual heterogeneity in the host family model (I^2^ host phylogeny = 25.86%, Table S12). The presence of symbionts in hosts had a negative effect on body size, survival and load of natural enemies (Table 3, Figure 5c). Both natural and introduced symbionts imposed a negative effect on natural enemy fitness (Table 3, Figure 5d). Reduction in natural enemy fitness was observed for most symbiont species, but no significant negative effect was detected for *Hamiltonella* or *Wolbachia* (Table 3, Figure 5e). However, this result should be interpreted with caution, as *Hamiltonella* had relatively few effect sizes (k = 20). Furthermore, the lack of a significant effect for both *Hamiltonella* and *Wolbachia* may reflect the variation in protective effects across host species, symbiont strains and natural enemy group/species/strain.

**Figure 5.**
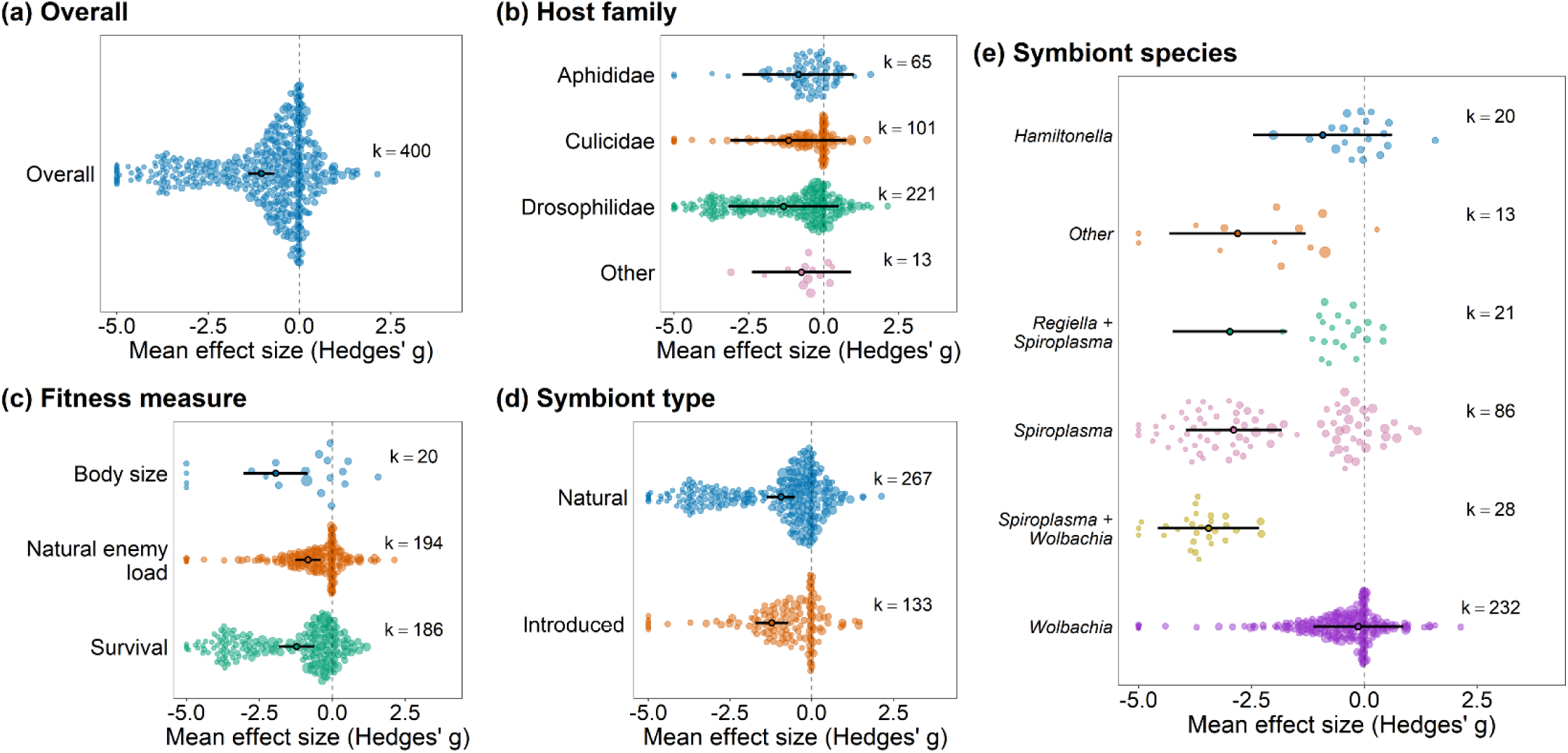
Effect of defensive symbionts on the fitness of natural enemies infecting symbiont-infected hosts. (a) overall model, (b) model including host family as moderator, (c) model including fitness measure as moderator, (d) model including symbiont type (i.e., if the symbiont naturally infects the host species or if it was artificially introduced) as moderator, and (e) model including symbiont species as moderator. Positive values indicate positive effects on host fitness. Hence, positive values indicate bigger body size, higher survival, and higher natural enemy load (quantity of natural enemies replicating inside the host such as viruses and bacteria). Points are the weighted mean effect sizes ± 95% confidence intervals. Each colored circle represents an effect size, scaled according to precision (inverse of the sampling error variance). ‘k’ is the number of effect sizes. Thirteen points had values below −5, which expanded the x-axis range and compressed the visual spacing of the remaining points. For visualization purposes, these points are shown at −5. All data points were included in the statistical analyses.

**Table 3.**
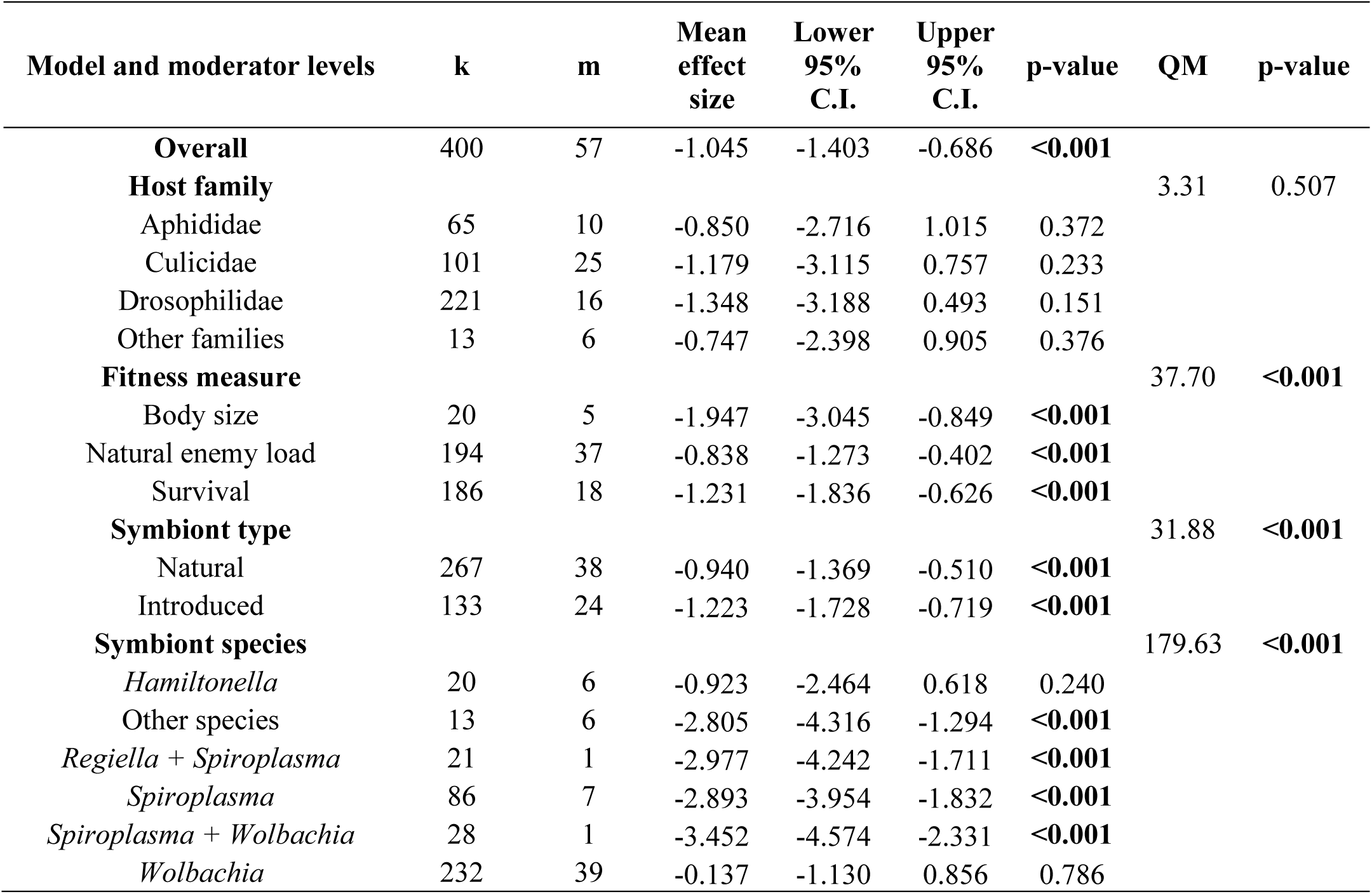
Results of the meta-analysis on the effect of symbionts on the fitness of natural enemies infecting hosts harboring symbionts and test of moderators (QM), including the overall model and the models including the following moderators: host family, fitness measure, symbiont type, and symbiont species. ‘k’ is the number of effect sizes and ‘m’ is the number of studies. p-values that are statistically significant are in bold (α =0.05).

| Model and moderator levels | k | m | Mean<br>effect<br>size | Lower<br>95%<br>C.I. | Upper<br>95%<br>C.I. | p-value | QM | p-value |
| --- | --- | --- | --- | --- | --- | --- | --- | --- |
| **Overall** | 400 | 57 | -1.045 | -1.403 | -0.686 | **<0.001** |  |  |
| **Host family** |  |  |  |  |  |  | 3.31 | 0.507 |
| Aphididae | 65 | 10 | -0.850 | -2.716 | 1.015 | 0.372 |  |  |
| Culicidae | 101 | 25 | -1.179 | -3.115 | 0.757 | 0.233 |  |  |
| Drosophilidae | 221 | 16 | -1.348 | -3.188 | 0.493 | 0.151 |  |  |
| Other families | 13 | 6 | -0.747 | -2.398 | 0.905 | 0.376 |  |  |
| **Fitness measure** |  |  |  |  |  |  | 37.70 | **<0.001** |
| Body size | 20 | 5 | -1.947 | -3.045 | -0.849 | **<0.001** |  |  |
| Natural enemy load | 194 | 37 | -0.838 | -1.273 | -0.402 | **<0.001** |  |  |
| Survival | 186 | 18 | -1.231 | -1.836 | -0.626 | **<0.001** |  |  |
| **Symbiont type** |  |  |  |  |  |  | 31.88 | **<0.001** |
| Natural | 267 | 38 | -0.940 | -1.369 | -0.510 | **<0.001** |  |  |
| Introduced | 133 | 24 | -1.223 | -1.728 | -0.719 | **<0.001** |  |  |
| **Symbiont species** |  |  |  |  |  |  | 179.63 | **<0.001** |
| *Hamiltonella* | 20 | 6 | -0.923 | -2.464 | 0.618 | 0.240 |  |  |
| Other species | 13 | 6 | -2.805 | -4.316 | -1.294 | **<0.001** |  |  |
| *Regiella* + *Spiroplasma* | 21 | 1 | -2.977 | -4.242 | -1.711 | **<0.001** |  |  |
| *Spiroplasma* | 86 | 7 | -2.893 | -3.954 | -1.832 | **<0.001** |  |  |
| *Spiroplasma* + *Wolbachia* | 28 | 1 | -3.452 | -4.574 | -2.331 | **<0.001** |  |  |
| *Wolbachia* | 232 | 39 | -0.137 | -1.130 | 0.856 | 0.786 |  |  |

Both single and coinfections with symbionts reduced natural enemy fitness, but the fitness reduction caused by symbiont coinfection was 2.5 times greater than the reduction caused by single infection (Table S9, Figure S4a). Nematodes, parasitoids, protozoans, and viruses had significant reductions in fitness when infecting symbiont-infected hosts, but no significant effect was detected for bacteria (Table S9, Figure S4b). However, interpretation is limited by the small number of effect sizes of the bacteria group (k = 22). Symbionts reduced the fitness of both facultative and obligate lethal natural enemies (Table S9, Figure S4c)

### 3.2. Subgroup analyses: *Wolbachia* and aphids

#### Wolbachia only

For protection data, mean overall effect size was small and positive, but non-significant (Figure S5a, Table S10). Although most effect sizes were distributed on the positive side (Figure S5a), the variation in the protective effects of *Wolbachia* likely contributed to the non-significant overall effect. *Wolbachia*-infected hosts had a higher proportion of individuals not infected or not parasitized by natural enemies (uninfected hosts), but no significant effect was detected for fecundity or survival (Figure S5c, Table S10). However, this result should be interpreted with caution, as fecundity had few effect sizes (k = 11). Furthermore, while most effect sizes were distributed on the positive side (Figure S5c), the variation in the effects of different *Wolbachia* strains on host survival likely contributed to the non-significant overall effect. Among natural enemy groups, only hosts infected with virus showed a significant fitness increase due to *Wolbachia* presence (Figure S5e, Table S10). *Wolbachia* increased the fitness of hosts being attacked by facultative lethal natural enemies, but not obligate lethal (Figure S5f, Table S10). No significant effect was detected for host family or symbiont type (Figures S5b and S5d, Table S10).

Regarding the cost to hosts data, *Wolbachia* showed little evidence of affecting host fitness in the absence of natural enemies, as the mean overall effect size was close to zero and non-significant. Most effect sizes were clustered close to zero, although both positive and negative effects were observed across studies (Figure S6a, Table S10). *Wolbachia*-infected hosts had bigger body size, but no significant effect was detected in development time, fecundity, and survival (Figure S6c, Table S10). No significant effect was detected for host family or symbiont type (Figures S6b and S6d, Table S10).

Regarding the fitness of natural enemies attacking *Wolbachia*-infected hosts, mean overall effect size was moderate and negative but non-significant. Although most effect sizes were distributed on the negative side, variation in the effects of different *Wolbachia* strains on natural enemy fitness may have contributed to the non-significant overall effect (Figure S7a, Table S10). No significant effects were detected for host family, fitness measure, symbiont type, natural enemy group, or natural enemy killing strategy (Figures S7b-S7f, Table S10).

#### Aphids only

For protection data, mean overall effect size was moderate to high and positive, indicating that symbiont-infected aphids had positive effect on their fitness when they were under attack of natural enemies (Figure S8a, Table S11). Symbiont-infected aphids had higher survival and a higher proportion of individuals not infected or not parasitized by natural enemies (uninfected hosts), but no significant effect was detected for fecundity (Figure S8b, Table S11). However, this result should be interpreted with caution, as fecundity had a small sample size (k = 10). Both natural and introduced symbionts increased aphid fitness (Figure S8c, Table S11). All symbiont species showed evidence of protecting aphids against natural enemies, except for *Spiroplasma* single infection and *Regiella* and *Spiroplasma* coinfection (Figure S8d, Table S11). Both single infections and coinfections with symbionts improved aphid fitness (Figure S8e, Table S11). Symbionts protected aphids against fungi (facultative lethal) and parasitoids (obligate lethal; Figure S8f and S8g, Table S11).

For cost to hosts data, mean overall effect size was moderate and negative, indicating that symbionts impose fitness costs on aphids in the absence of natural enemies (Table S11). Most effect sizes were clustered on the negative side, although both positive and negative effects were observed across studies (Figure S9a). Fecundity and survival were negatively affected by symbiont infection, but little evidence of negative effects on body size and development time were detected (Figure S9b, Table S11). Natural symbionts significantly reduced host fitness, but no significant effect was detected for introduced symbionts (Figure S9c, Table S11). Single symbiont infections reduced host fitness, but no significant effect was detected for coinfections (Table S11). However, for coinfection, most effect sizes were distributed on the negative side, and the upper confidence interval only slightly crossed zero (Figure S9d). Among symbiont species, only *Hamiltonella* and the coinfection of *Hamiltonella* and *Rickettsiella* showed evidence of reducing host fitness (Figure S9e, Table S11). However, these results should be interpreted with caution as most symbiont species had few effect sizes (Table S11).

Regarding the cost to natural enemies, mean overall effect size was negative but non-significant (Figure S10a, Table S11). No significant differences were detected among fitness measures, symbiont species, or symbiont number (Figures S10b-d, Table S11). Survival of natural enemies showed a negative and non-significant effect, but most effect sizes were clustered on the negative side with the upper confidence interval only slightly crossing zero (Figure S10b). The same was observed for symbiont coinfection (Figure S10d). However, most moderator levels were represented by few effect sizes, and therefore these results should be interpreted with caution. Furthermore, these results likely reflect the high variability in the protective effects of *Hamiltonella*, which are known to depend on specific combinations of host genotype, symbiont strain, and natural enemy species and strain.

### 3.3. Heterogeneity and publication bias

All models presented high heterogeneity (I^2^_total_ > 80%, Tables S12 and S13). In most models, effect size ID (within-study variance) explained the largest proportion of variance, followed by study ID (between-study variance), and then host species ID. Host phylogeny explained very little of the variance in most cases (I^2^_host phylogeny_ < 1%). An exception to this was observed in the models ‘host family’ and ‘symbiont species’ from the cost to natural enemies’ data of the main analysis, in which host phylogeny explained approximately 26% and 22% of the variance, respectively (Table S12). Similarly, in all six models from the cost to natural enemies’ data of the ‘*Wolbachia* only’ subgroup analysis (Table S13), host phylogeny explained 29-35% of model variance (Table S13).

We found evidence of publication bias in 18 of the 22 models in the main analysis (Table S14). In the ‘*Wolbachia only*’ subset analysis, we detected evidence of publication bias in six of the 16 models (Table S15). However, evidence of publication bias was found in 15 of the 16 models in the ‘aphids’ subset analysis (Table S15). Although the presence of publication bias does not invalidate our findings, these results should be interpreted with caution.

### 3.4. Sensitivity analysis

Influence diagnostics did not identify any influential observations in the overall models from the main analysis and the subgroup analysis (Cook’s distance: *D* < 0.72, DFBETAS < 0.24; Table S16), suggesting that no individual data point had a disproportionate effect on model estimates. However, we found two influential points (Cook’s distance: *D* > 1; Table S16) in the trade-off between protection and fitness costs analysis.

## 4. Discussion

In this meta-analysis we collected data from 246 studies, comprising around 3,425 effect sizes across many different study systems, which allowed us to make a broad assessment of the generality of symbiont effects on host fitness. The overall models from our main analyses revealed that the protection provided by symbionts (mean effect size = 0.588) is approximately 7.1 times stronger than the cost in the absence of natural enemies (mean effect size = −0.083), indicating strong protective effects and little evidence of substantial fitness costs in symbiont-infected hosts. We also found that the outcome of host–symbiont interactions varies among host families, symbiont species, and natural enemy group. Moreover, not all fitness components contribute equally to the overall conclusions. For example, in the presence of natural enemies, symbionts have a substantial positive effect on host survival, but no detectable effect on fecundity. In the following section, we provide a more comprehensive discussion of these results.

Zytynska et al. (2021) performed a meta-analysis on the costs and benefits of harboring facultative symbionts in aphids, whiteflies and heteropterans. Overall, they found that whiteflies have higher fecundity and shorter development time whereas aphids have increased resistance to natural enemies, but lower fecundity when infected by symbionts. This is in accordance with our findings where aphids harboring symbionts have strong protection against natural enemies such as parasitoids and fungi, and a substantial cost to their fitness, especially their fecundity and survival. By comparing our dataset, we identified 48 overlapping studies, representing approximately 20% of our total dataset (246 studies) meaning that around 80% of our study is composed of new data on defensive symbiosis. This is expected given our inclusion of terrestrial arthropods, particularly two major study systems: *Drosophila–Wolbachia–*virus and mosquito–*Wolbachia*–virus. Together, these additions expand their important findings on host-symbiont interactions.

We found that, in the presence of natural enemies, symbiont-infected hosts have higher fitness than uninfected hosts, indicating a general pattern of symbiont-mediated protection against natural enemies in insects. Interestingly, in the absence of natural enemies, we found little evidence of substantial fitness costs in symbiont-infected hosts, and observed a high variability across systems, with interactions ranging from highly beneficial to highly costly. Furthermore, when we analyzed the trade-off between protection and cost to host fitness, we found no clear pattern of higher levels of symbiont protection being associated with higher fitness costs. Collectively, these results suggest that over the coevolutionary course, selection may lead to the evolution of symbiont strains that provide moderate to high protection against natural enemies at low costs, at least in certain systems. Our findings support results and predictions from prior studies. In pea aphids, it has been reported mild fitness costs associated with *Hamiltonella* infection (Leclair et al., 2016; Martinez et al., 2018; Oliver et al., 2006), which have strains that provide moderate to high protection against parasitoids (Doremus et al., 2018; Martinez et al., 2018; Oliver et al., 2005). Martinez et al. (2015) observed a natural strain of *D. simulans, w*Au, that is highly protective while imposing little cost on egg hatch rates in the absence of viruses, suggesting that selection may sometimes reduce the trade-off between protection and cost. Therefore, our findings suggest that strong symbiont-mediated protection combined with low fitness costs may, in part, explain the persistence of symbionts in natural populations.

Our results showed no evidence of protection against natural enemies in other arthropod families, with protection detected only in Aphididae and Drosophilidae. Protection against natural enemies may be one of the reasons for symbionts being so widely spread among arthropods (Pimentel et al., 2021; Zug and Hammerstein, 2015). However, studies on protection against natural enemies are still limited mostly to aphid and dipteran hosts. Besides studies on different species from families Aphididae, Culicidae and Drosophilidae, we only found studies that tested for symbiont protection against natural enemies in Diptera: Glossinidae (Schneider et al., 2019), Hemiptera: Aleyrodidae (Ghosh et al., 2018), Trombidiformes: Tetranychidae (Zélé et al., 2020), Isopoda: Armadillidiidae (Braquart-Varnier et al., 2015), and Isopoda: Porcellionidae (Braquart-Varnier et al., 2015). Thus, further studies across diverse arthropod families are essential to better understand how protection against natural enemies affects symbiont prevalence in natural populations of terrestrial arthropods.

We found that the interaction is substantially costly for aphids. This is in accordance with Zytynska et al. (2021), that found that the interaction with symbionts reduced fecundity and lifespan of aphids, but not of whiteflies. In the subgroup analysis restricted to hosts from the family Aphididae, the cost was specifically associated with host fecundity, survival, and development time in hosts infected with *Hamiltonella*, *Spiroplasma*, and *Hamiltonella + Rickettsiella* co-infection. Although defensive symbionts have successfully spread among insects, the level of colonization vary between symbiont and host species. For example, *Wolbachia* frequency in natural insect populations varies from 10% to 100% (Kajtoch and Kotásková, 2018; Kriesner et al., 2016). Aphid symbiont *Serratia* occurs at low frequencies (approximately 20%) in nature, despite offering protection against parasitoids (Pons et al., 2022), while *Hamiltonella* frequency can vary between approximately 20 to 70% (Smith et al., 2021). This suggests that harboring certain symbionts species and strains involves a moderate to high cost to aphid fitness in the absence of selection by parasitoids. In addition to the balance between cost and protection, some symbionts have other strategies to increase in frequency in nature. For instance, certain strains of *Wolbachia* and *Spiroplasma* have successfully colonized natural populations via reproductive parasitism (Himler et al., 2011; Martinez et al., 2015; Zug & Hammerstein, 2015). Therefore, in the absence of natural enemies, some symbiont strains can cease to be a facultative mutualist and act exclusively as a reproductive parasite. This is the case for some *Wolbachia* strains that cause cytoplasmic incompatibility (CI) while providing antiviral protection (Martinez et al., 2015). Thus, our results suggest that one possible explanation for aphid symbionts having lower frequencies in natural host populations might be due to relatively moderate to high costs associated with the interaction, and the lack of other strategies to spread among and within populations, such as reproductive parasitism.

We found that both natural and introduced symbionts confer protection against natural enemies, but neither showed evidence of an overall pattern of negative effects on host fitness in the absence of natural enemies. These results are contrary to our initial hypothesis. Among fly and mosquito hosts, there is an overall trend in which symbiont strains that provide stronger protection occur in higher densities within host cells (Martinez et al., 2014; Martinez et al., 2015; Zug & Hammerstein, 2015). Therefore, it is common to transinfect high density strains into new hosts to test for protection against natural enemies (Moreira et al., 2009; van den Hurk et al., 2012; Zug & Hammerstein, 2015), especially in the Culicidae family to control insect-born human diseases (Bian et al., 2010; Hughes et al., 2011; Moreira et al., 2009; van den Hurk et al., 2012; Werren et al., 2008; Xue et al., 2018). However, hosts have a reduction in their fitness when they are not infected with natural enemies but are carrying these high-density strains (Martinez et al., 2014; Martinez et al. 2015; Martinez et al., 2016). For example, Martinez et al. (2014) transferred 19 *Wolbachia* strains with different levels of density into *D. simulans* to test them for antiviral protection. After transinfecting *Wolbachia* strains into a new recipient species, usually they keep the same level of density as in their original host. Flies infected with high density strains *w*Mel and *w*MelCS, natural from *D. melanogaster*, had higher survival and lower viral load when infected with virus. However, when flies were not infected with virus, these same strains caused reduction in fecundity and lifespan of flies. Interestingly, when flies were infected with natural and highly protective strain *w*Au, fitness costs were lower (Martinez et al., 2015). Another interesting study using *w*Au showed that despite this strain not presenting a reproductive parasitism phenotype – which is one of the known mechanisms that *Wolbachia* uses to spread in arthropod populations – this strain is still able to spread in field populations of its natural host, *D. simulans* (Kriesner et al., 2013). This alone suggests that symbionts have different mechanisms to propagate, one of them potentially being protection against natural enemies. Thus, the level of protection and the cost associated depends on the symbiont strain that is infecting the host, and this might influence the ability of symbionts to spread in nature. Moreover, one possible explanation is that early on the host-symbiont interaction the cost of harboring symbionts is high, but it decreases over evolutionary time, facilitating the propagation of symbionts. This could provide a possible explanation for our results. However, it is important to point out that our dataset is very diverse in terms of symbiont density – there are symbiont strains which have high density, strains with low density, and strains with no information on symbiont density. We were unable to conduct a more detailed analysis comparing specific symbiont strains, including those commonly used in transinfection experiments, because of limited statistical power. Therefore, although variation in symbiont density and the evolutionary history of host–symbiont associations may contribute to the patterns we found, our data do not allow us to directly assess these effects.

Looking at the overall results for costs to hosts, we found a small, negative, and non-significant mean effect size, with symbiont effects ranging from highly negative to highly positive. Similarly, for costs to natural enemies, we found no significant effects on the fitness of natural enemies infecting hosts harboring *Hamiltonella* or *Wolbachia.* These results likely reflect the variation in protective effects across specific combinations of host species-symbiont strain-natural enemy group/species/strain, as well as the variation in the associated costs across specific host species-symbiont strain combinations. In *Hamiltonella*, for example, this variation can be explained by genotype-by-genotype (G×G) interactions between *Hamiltonella* and natural enemy strains – i.e., the fitness of different parasitoid strains can vary depending on the *Hamiltonella* strain infecting the host (Cayetano and Vorburger, 2015). For example, in *Aphis fabae*, different *Lysiphlebus fabarum* strains varied in their ability to parasitize aphids infected with different *Hamiltonella* strains, with some parasitoid strains able to parasitize aphids infected with one *Hamiltonella* strain but not with another strain (Gimmi and Vorburger, 2024). Therefore, while broad studies such as our meta-analysis reveal general patterns, studies of specific host–symbiont–natural enemy systems are essential for uncovering context-dependent effects that may be obscured at broader scales.

We found that, in the presence of natural enemies, symbionts increased host survival and their likelihood of avoiding infection, but were unable to recover host fecundity. This was expected, since the great majority of our dataset consists of survival and parasitism data, with only a few data points that measured host fecundity (k = 34). Given our limited sample size, additional studies on host fecundity are needed to clarify how symbionts influence this fitness component in the presence of natural enemies. However, one interesting case, is that *Spiroplasma* strains that infect *D. neotestacea* restore host fecundity when flies are infected by nematodes of the genus *Howardula*, which sterilize the female flies (Haselkorn et al., 2013; Jaenike et al., 2010). We also found that symbiont infection in the absence of natural enemies increased body size and shortened development time, while survival was decreased, and no significant effect on fecundity was detected. This may have a positive impact on overall fitness, since larger individuals are expected to have higher reproductive success (Beukeboom, 2018), and individuals that develop faster are exposed to natural enemies such as parasitoids for shorter periods of time (Nijhout et al., 2010). Thus, despite the cost in survival, the benefit in development time and body size may be reducing the overall cost of the interaction.

Our results showed both natural and introduced symbionts reduced the survival, load, and body size of natural enemies. Symbionts can compete for resources within the host, release toxins or activate the host’s immune response, directly affecting the chance of success of the natural enemy (Vorburger and Perlman, 2018). In aphids, *Hamiltonella* releases toxins that are produced by bacteriophages present in its genome and kill the early life stages of parasitoids (Vorburger and Gouskov, 2011; Weldon et al., 2013). Although we do not know exactly which mechanism is adopted by *Wolbachia*, *Spiroplasma* and other aphid symbionts (Zug and Hammerstein, 2015), our results suggest a general pattern of symbionts reducing the fitness of natural enemies that attack their hosts. However, it is important to notice that there is a wide functional diversity among natural enemies – i.e., natural enemies can be either obligate lethal (such as parasitoids) or facultative lethal (such as viruses, for example). We found that symbionts reduced the fitness of both facultative and obligate lethal natural enemies. However, viruses are a common pathogen in nature, but their virulence (i.e., the capacity to cause damage in a host) varies greatly. Looking specifically at the viruses that infect *Drosophila* for example, DCV is frequently studied in the laboratory but rarely found in nature due to its virulence, killing the flies within days (Kapun et al., 2010). Other viruses that are more frequently found in nature such as DAV and Nora similarly affect the fitness of their hosts, although to a lesser extent than DCV. DAV reduces the fecundity of flies (Brosh et al., 2022), and both Nora and DAV kill their host, but at a much slower rate than DCV (Ambrose et al., 2009; Habayeb et al., 2006). If natural enemies exert only mild effects on host fitness, the fitness benefits of symbiont-mediated protection may consequently be smaller. Hence, the level of virulence of natural enemies may directly influence the outcome of host-symbiont-natural enemy interactions.

### Caveats

The great majority of studies addressing symbiont protection against natural enemies are performed under laboratory conditions, minimizing biotic and abiotic factors that could influence the outcome of the host-symbiont-natural enemy interactions in nature. For aphids, it has been shown that under laboratory conditions, *Hamiltonella* is able to increase host resistance against parasitoids and that the level of protection varies among host and parasitoid genotypes (Cayetano and Vorburger, 2015). Few studies have tested *Hamiltonella* protection against parasitoids in the field, and some suggested that *Hamiltonella* reduces aphid mortality against specific parasitoid strains but there was no overall fitness benefit for the hosts possibly because of the cost of the host-symbiont interaction (Hrček et al., 2016; Rothacher et al., 2016). A recent study has shown no significant relationship between the presence of parasitoid wasps and *H. defensa* prevalence in the field as well as no strong evidence for symbiont cost in the absence of parasitoids. Instead, an important factor modulating symbiont prevalence in the field could be the temperature (Smith et al., 2021). In dipterans, only very recently has it been shown that *Wolbachia* protects flies against virus infection in nature (Cogni et al., 2021). This study showed that in a natural *D. melanogaster* population, flies infected with *Wolbachia* are less likely to be infected by viruses (Cogni et al., 2021). Regarding the effect of temperature, it has been demonstrated in the laboratory that the *Wolbachia* antiviral effect observed when flies develop at high temperatures is reduced when flies develop at low temperatures (Chrostek et al., 2021; Martins et al., 2023). These studies give a hint that not only defensive symbiosis can be a factor modulating symbiont prevalence in the field, but temperature might be an important factor as well. The lack of studies performed in the field is a major caveat and more studies investigating protection against natural enemies in natural populations are essential for understanding the ecological importance of defensive symbiosis and what factors modulate the prevalence of symbionts in field populations.

Concerning heterogeneity and publication bias, all of our meta-analytical models had high heterogeneity, which is expected in biological meta-analysis. However, even with the inclusion of moderators in the models, the amount of heterogeneity remained high, suggesting that there are other factors (i.e., moderators not included in our meta-analysis) that may influence the results. Moreover, we found evidence for publication bias in most of our models, which requires caution in interpreting the results. However, it does not invalidate our findings, since adding more effect sizes to our data could reduce the mean effect sizes but maintain the same results. Such bias can occur because studies with positive or statistically significant results are more likely to be published than studies with negative or inconclusive findings. Several factors can lead to publication bias: (1) difficult access to all existing studies due to the language of the publication, impact factor of the journal or number of citations of the article (Jennions et al., 2013), (2) failure to present the results, since researchers can report only data based on the significance of the results and direction of effect sizes that corroborate their hypothesis (Jennions et al., 2013; Thornton and Lee, 2000), (3) reviewers and editors of journals who tend to deny negative results (Thornton and Lee, 2000), and (4) there is a possibility that these studies simply do not exist yet. Thus, we encourage researchers, reviewers, and editors to publish any results obtained in high-quality studies, regardless of their significance or direction of effect size. The dissemination of this practice can help to reduce publication bias in systematic and narrative reviews.

### Conclusion

In summary, our study shows that: (1) in the presence of natural enemies, symbionts provide strong protection to their hosts, (2) in the absence of natural enemies, there is little evidence of substantial fitness costs associated with symbiont infection, and (3) natural enemies attacking symbiont-infected hosts have substantial reduction in their fitness. Our results reveal broad symbiont-mediated protection with little evidence of substantial costs associated, which may help explain the successful spread of symbionts among terrestrial arthropods, especially insects. This suggests that symbionts can play an important role in natural host populations, acting as a complement to the endogenous defenses of hosts. However, the level of protection and cost for both hosts and natural enemies varies between the fitness components, host families, symbiont species, and natural enemy group.

Defensive symbionts vary from low to high frequencies in wild insect populations, suggesting that there are many factors that can influence the spread of symbionts in nature such as if the symbiont strain causes reproductive parasitism, as well as the strength of the reproductive parasitism phenotype and the efficiency of maternal transmission; how strong is the protection that symbionts offer against natural enemies (which may vary with symbiont density and natural enemy virulence, for example); how high is the cost of the interaction with symbionts; and if the natural enemies’ frequencies vary temporally in nature. Furthermore, temperature and nutritional provisioning can also be an important factor in modulating the prevalence of symbionts in nature.

We hope our study stimulates researchers to expand the investigation on defensive symbiosis, collecting information on protection in other terrestrial arthropod families. Hopefully, in the future, this will allow us to investigate if these findings expand to other arthropods.

## Acknowledgments

We would like to thank all researchers that responded to our requests for data. We are grateful to André C. Pimentel, Laura C. Leal, Camila T. Castanho, Ana Paula A. Assis, Brandon S. Cooper, Julien Martinez, Karla Yotoko and Maria Vibranovski for the suggestions on the manuscript. Cássia S. Cesar is funded by Fundação de Amparo à Pesquisa do Estado de São Paulo (FAPESP; 2019/03997-2 and 2025/03375-2). Rodrigo Cogni is funded by FAPESP (2013/25991-0 and 2021/06874-9), Conselho Nacional de Desenvolvimento Científico e Tecnológico (CNPq; 307015/2015-7 and 307447/2018-9), and a Newton Advanced Fellowship from the Royal Society.

## Data availability statement

All data collected and R script are available at Open Science Framework (OSF), doi:10.17605/OSF.IO/CT9ME.

## Author Contributions

Designed research: Rodrigo Cogni, Cássia S. Cesar

Performed research: Cássia S. Cesar

Analyzed data: Cássia S. Cesar, Eduardo S. A. Santos

Wrote the paper: Cássia S. Cesar, Rodrigo Cogni

## Supplementary information

## Supplementary figures

**Figure S1.**
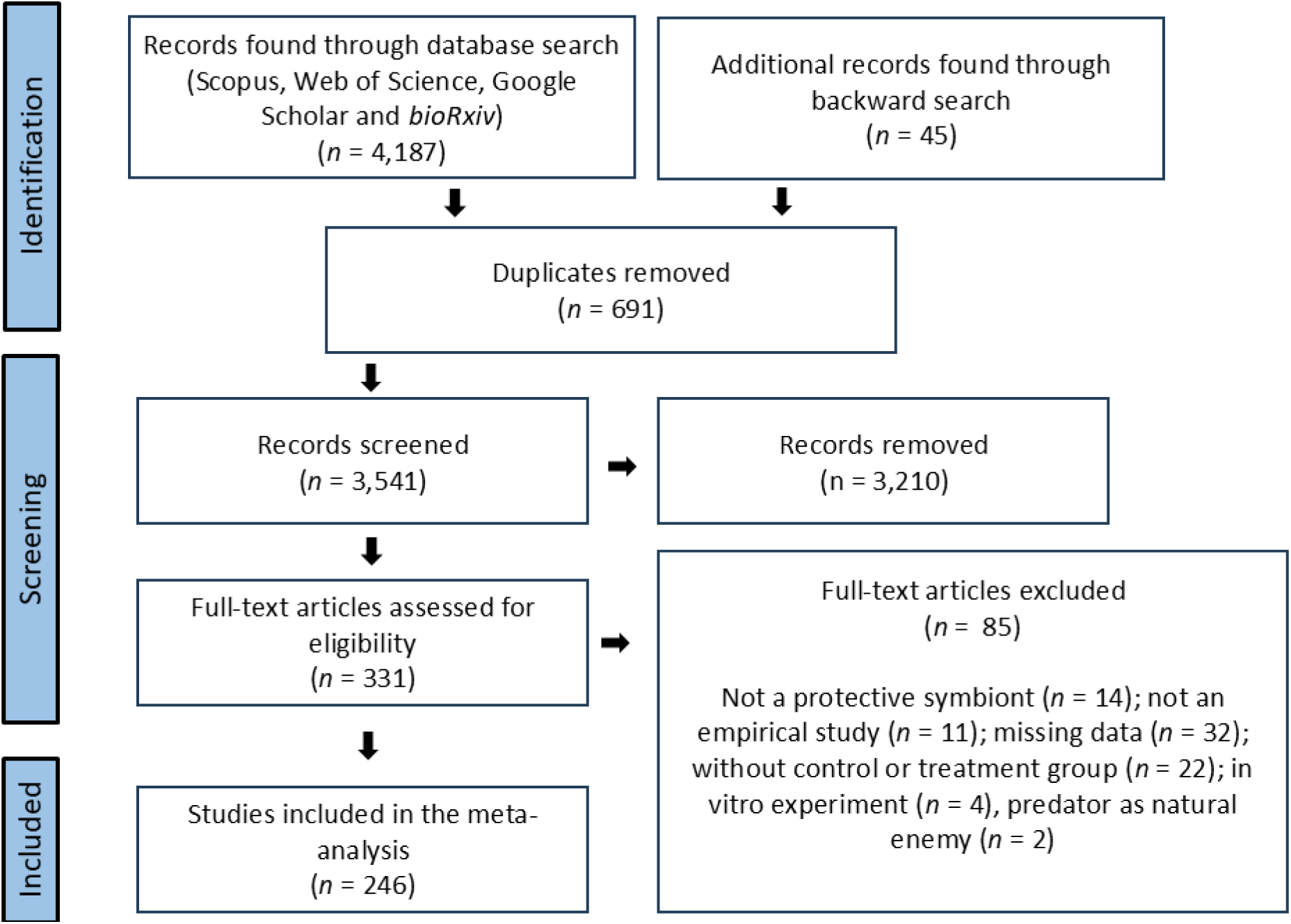
Full screening process of inclusion and exclusion of studies following the PRISMA protocol.

**Figure S2.**
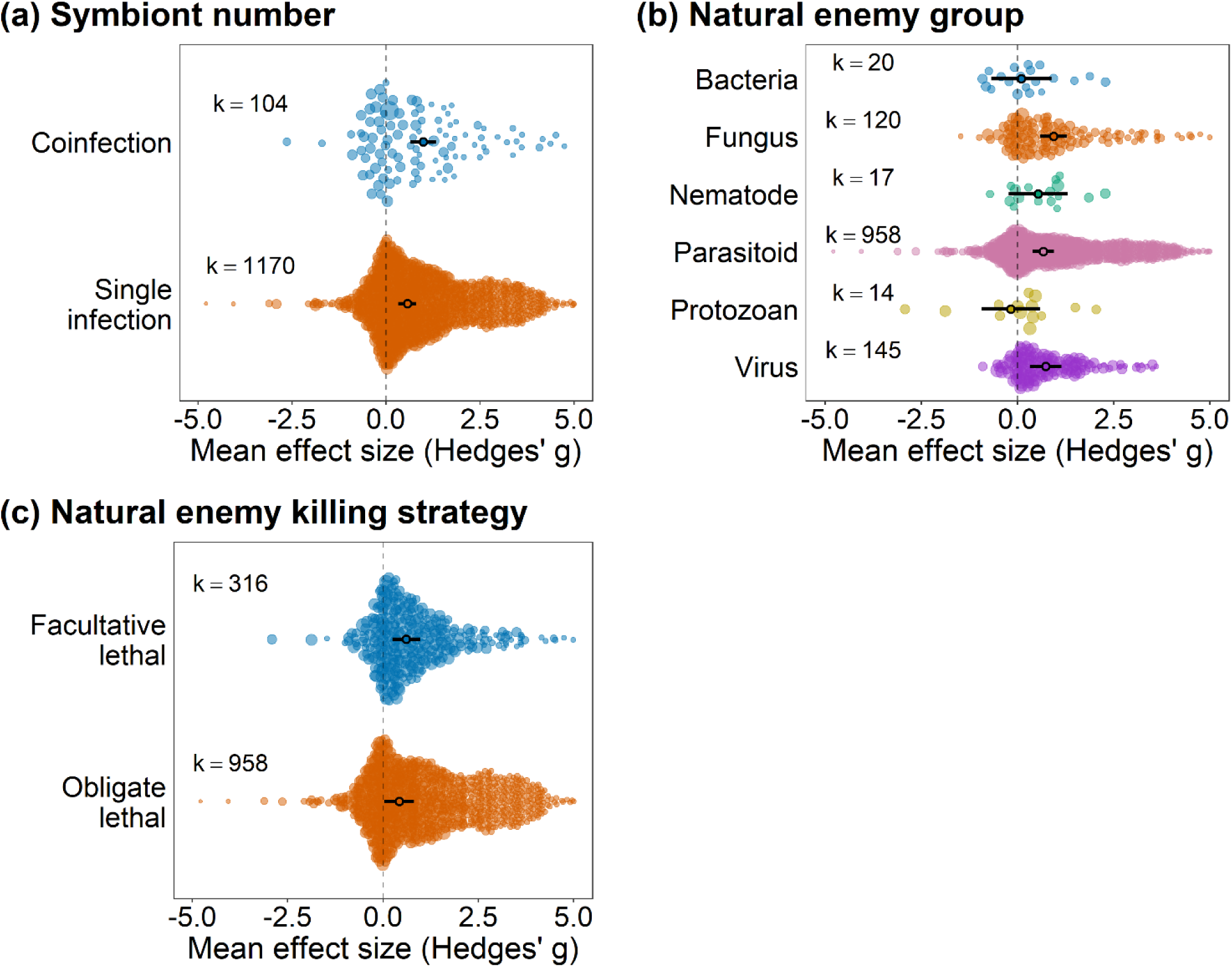
Effect of defensive symbionts on the fitness of hosts that are under attack of natural enemies. (a) model including symbiont number as moderator, (b) model including natural enemy group as moderator, and (c) model including natural enemy killing strategy as moderator. Points are the weighted mean effect sizes ± 95% confidence intervals. Each colored circle represents an effect size, scaled according to precision (inverse of the sampling error variance). ‘k’ is the number of effect sizes.

**Figure S3.**
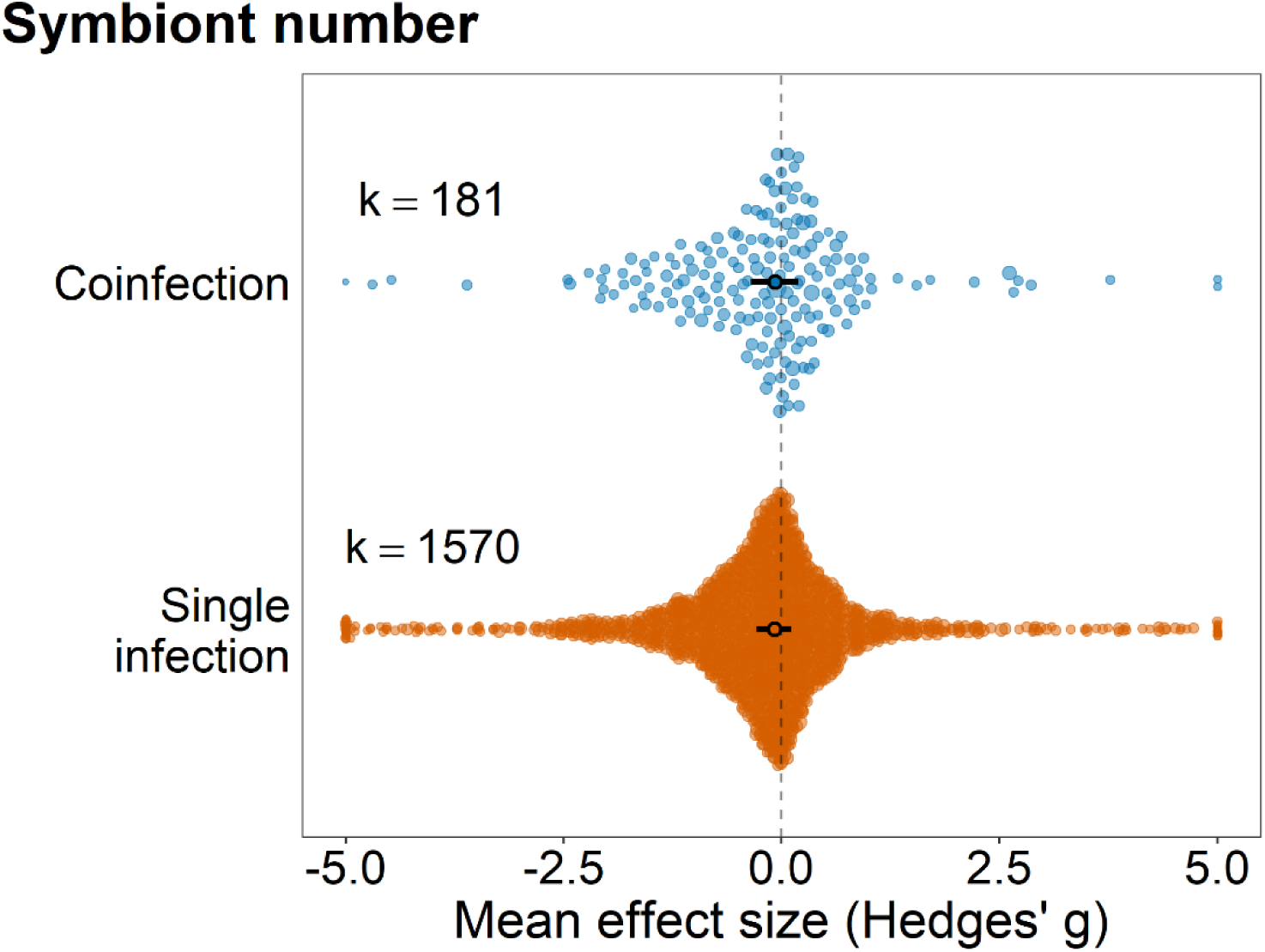
Effect of defensive symbionts on the fitness of hosts that are in the absence of natural enemies, including symbiont number as moderator. Points are the weighted mean effect sizes ± 95% confidence intervals. Each colored circle represents an effect size, scaled according to precision (inverse of the sampling error variance). ‘k’ is the number of effect sizes. Sixteen points fell below −5 and 13 points fell above +5, which expanded the x-axis range and compressed the visual spacing of the remaining points. For visualization purposes only, these points are shown at ±5. All data points were included in the statistical analyses.

**Figure S4.**
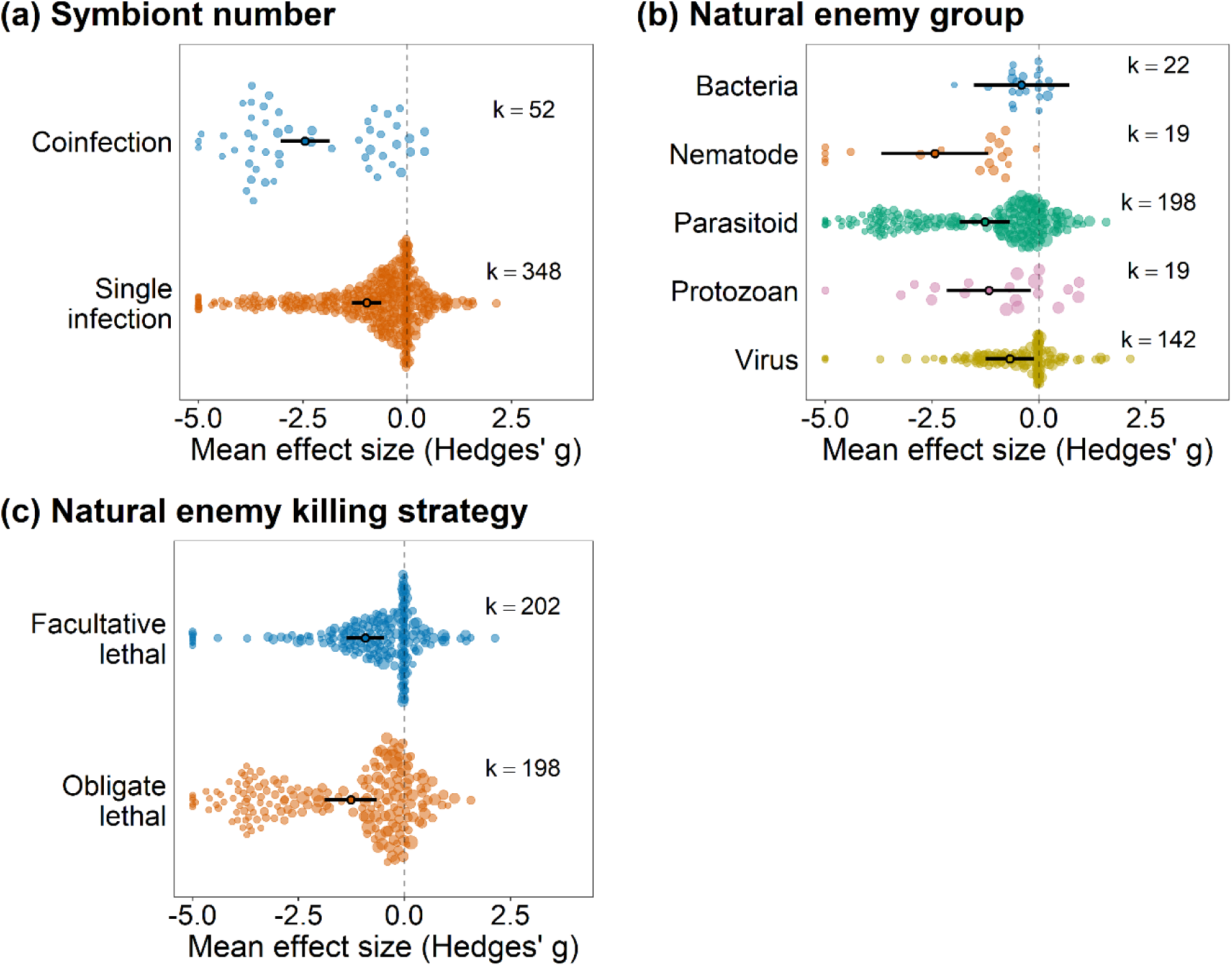
Effect of defensive symbionts on the fitness of natural enemies attacking symbiont-infected hosts. (a) model including symbiont number as moderator, (b) model including natural enemy group as moderator, and (c) model including natural enemy killing strategy as moderator. Points are the weighted mean effect sizes ± 95% confidence intervals. Each colored circle represents an effect size, scaled according to precision (inverse of the sampling error variance). ‘k’ is the number of effect sizes. Thirteen points had values below −5, which expanded the x-axis range and compressed the visual spacing of the remaining points. For visualization purposes, these points are shown at −5. All data points were included in the statistical analyses.

**Figure S5.**
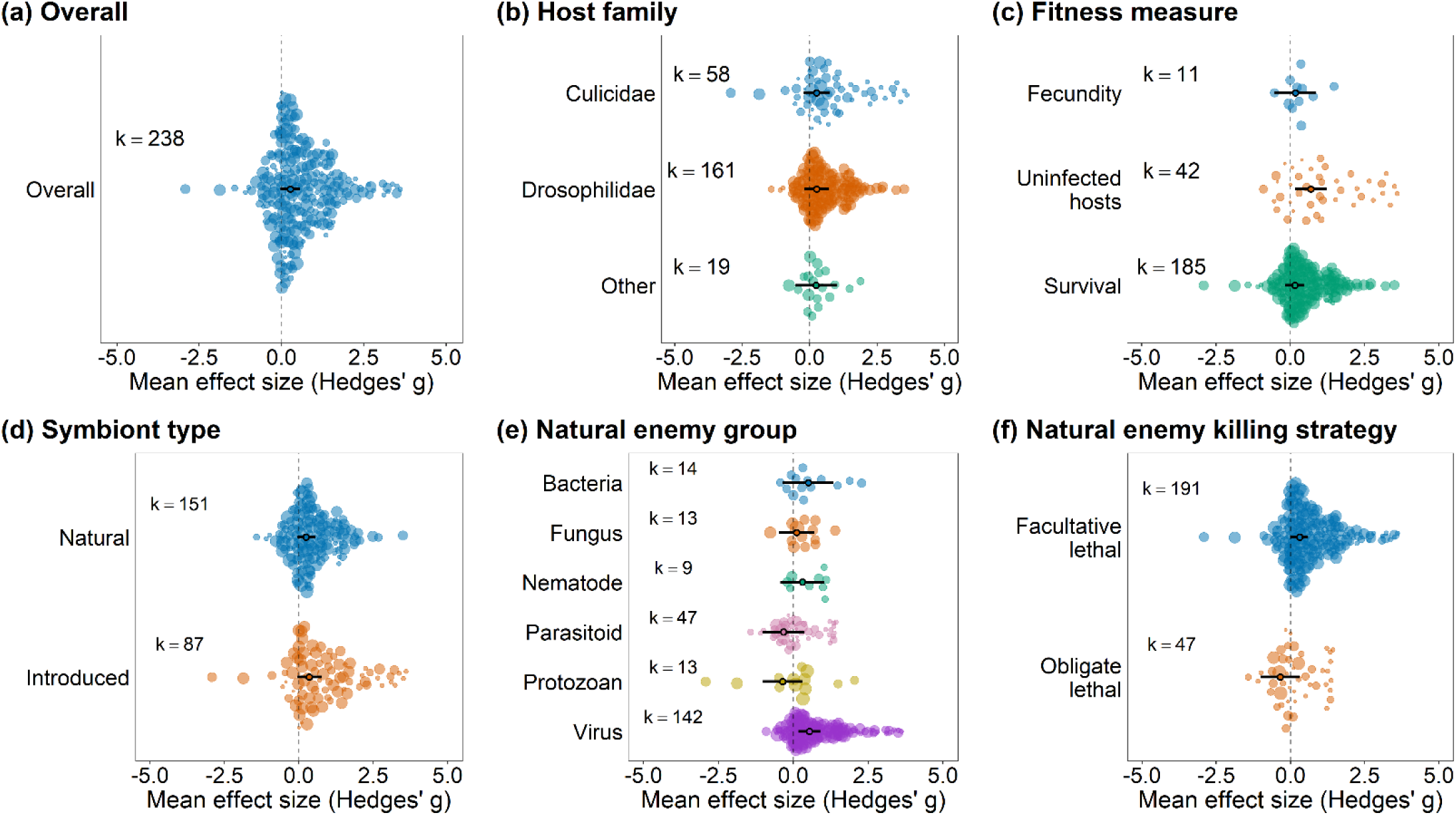
Sub-analysis: *Wolbachia* only. Effect of defensive symbionts on the fitness of hosts that are under attack of natural enemies. (a) overall model, (b) model including host family as moderator, (c) model including fitness measure as moderator, (d) model including symbiont type (i.e., if the symbiont naturally infects the host species or if it was artificially introduced) as moderator, (e) model including natural enemy group as moderator, and (f) model including natural enemy killing strategy as moderator. Positive values indicate positive effects on host fitness. Hence, positive values indicate higher fecundity, survival, and proportion of hosts that were uninfected (i.e., uninfected hosts - number of individuals that were unsuccessfully parasitized or infected by natural enemies). Negative values indicate that symbionts have a negative effect on host fitness. Points are the weighted mean effect sizes ± 95% confidence intervals. Each colored circle represents an effect size, scaled according to precision (inverse of the sampling error variance). ‘k’ is the number of effect sizes.

**Figure S6.**
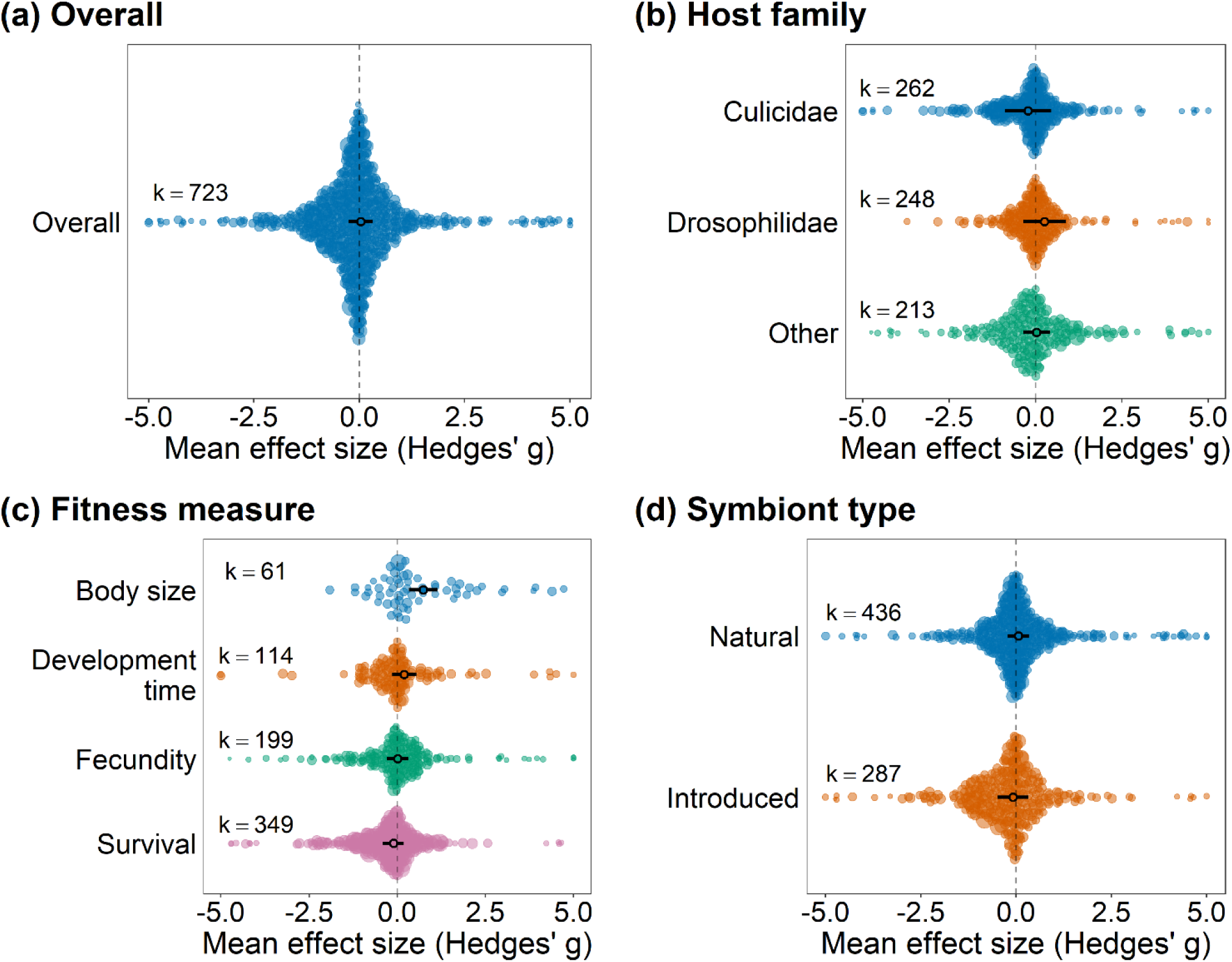
Sub-analysis: *Wolbachia* only. Effect of defensive symbionts on the fitness of hosts that are in the absence of natural enemies. (a) overall model, (b) model including host family as moderator, (c) model including fitness measure as moderator, and (d) model including symbiont type (i.e., if the symbiont naturally infects the host species or if it was artificially introduced) as moderator. Positive values indicate positive effects on host fitness. Hence, positive values indicate higher fecundity, survival, body size and shorter development time. Negative values indicate that symbionts have a negative effect on fitness. Points are the weighted mean effect sizes ± 95% confidence intervals. Each colored circle represents an effect size, scaled according to precision (inverse of the sampling error variance). ‘k’ is the number of effect sizes. Two points fell below −5 and four points fell above +5; for visualization purposes only, these points are shown at ±5. All data points were included in the statistical analyses.

**Figure S7.**
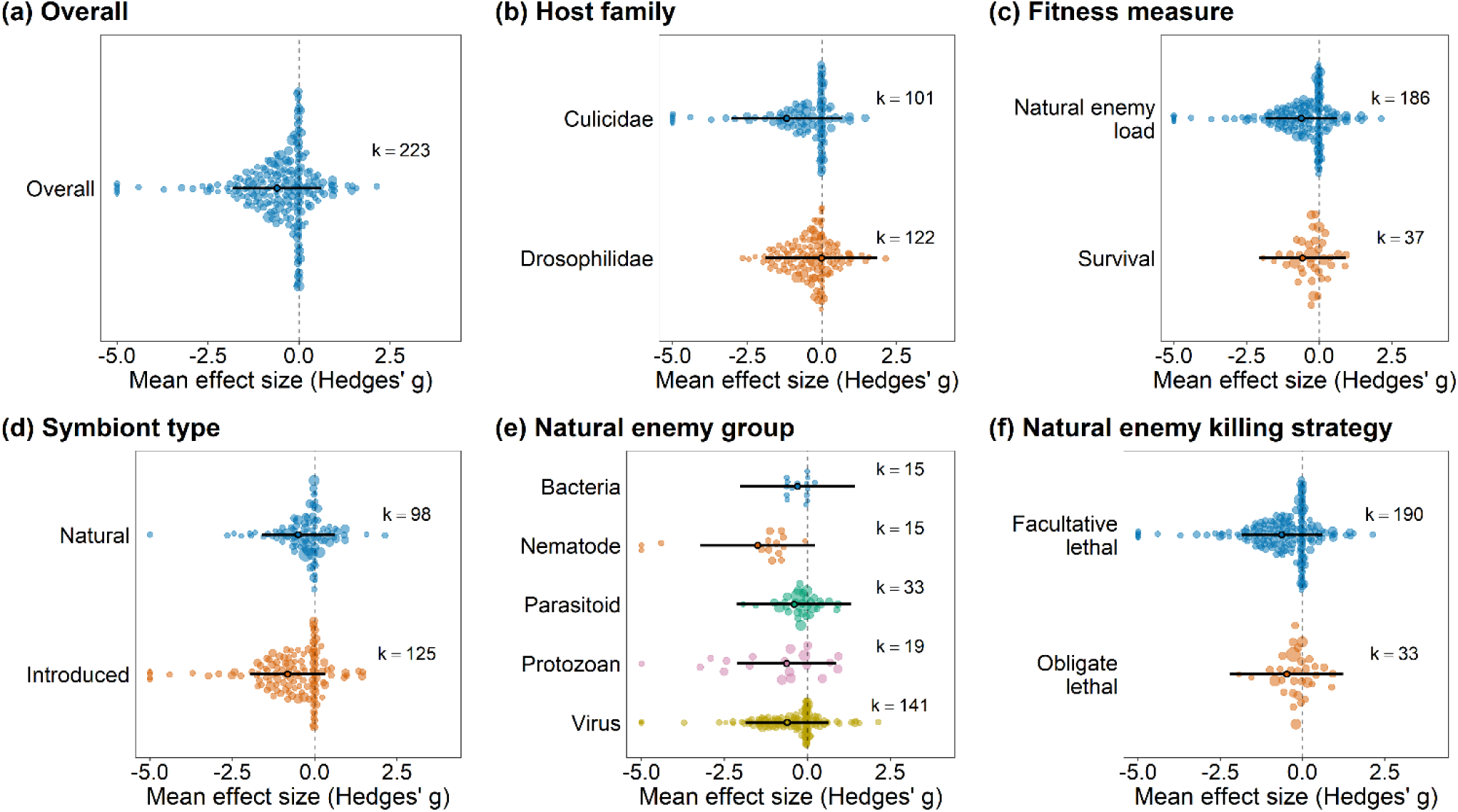
Sub-analysis: *Wolbachia* only. Effect of defensive symbionts on the fitness of natural enemies infecting symbiont-infected hosts. (a) overall model, (b) model including host family as moderator, (c) model including fitness measure as moderator, (d) model including symbiont type (i.e., if the symbiont naturally infects the host species or if it was artificially introduced) as moderator, (e) natural enemy group as moderator, and (f) model including natural enemy killing strategy as moderator. Positive values indicate positive effects on host fitness. Hence, positive values indicate higher survival, and higher natural enemy load (quantity of natural enemies replicating inside the host such as viruses and bacteria). Points are the weighted mean effect sizes ± 95% confidence intervals. Each colored circle represents an effect size, scaled according to precision (inverse of the sampling error variance). ‘k’ is the number of effect sizes. Six points fell below −5; for visualization purposes only, these points are shown at -5. All data points were included in the statistical analyses.

**Figure S8.**
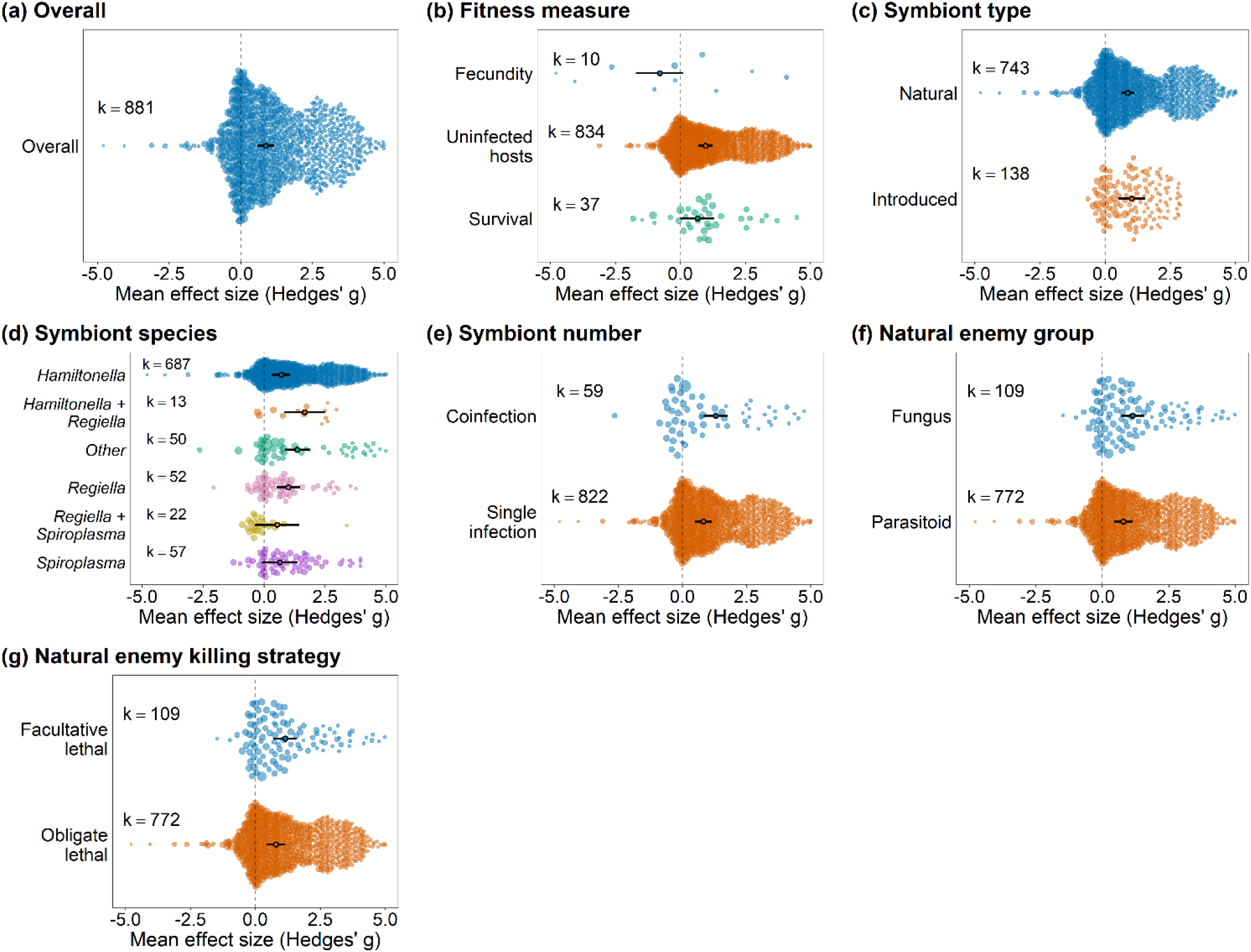
Sub-analysis: Aphids only. Effect of defensive symbionts on the fitness of aphids that are under attack of natural enemies. (a) overall model, (b) model including fitness measure as moderator, (c) model including symbiont type (i.e., if the symbiont naturally infects the host species or if it was artificially introduced) as moderator, (d) model including symbiont species as moderator, (e) model including symbiont number as moderator, (f) model including natural enemy group as moderator, and (g) model including natural enemy killing strategy as moderator. Positive values indicate positive effects on host fitness. Hence, positive values indicate higher fecundity, survival, and proportion of hosts that were uninfected (i.e., uninfected hosts - number of individuals that were unsuccessfully parasitized or infected). Negative values indicate that symbionts have a negative effect on host fitness. Points are the weighted mean effect sizes ± 95% confidence intervals. Each colored circle represents an effect size, scaled according to precision (inverse of the sampling error variance). ‘k’ is the number of effect sizes.

**Figure S9.**
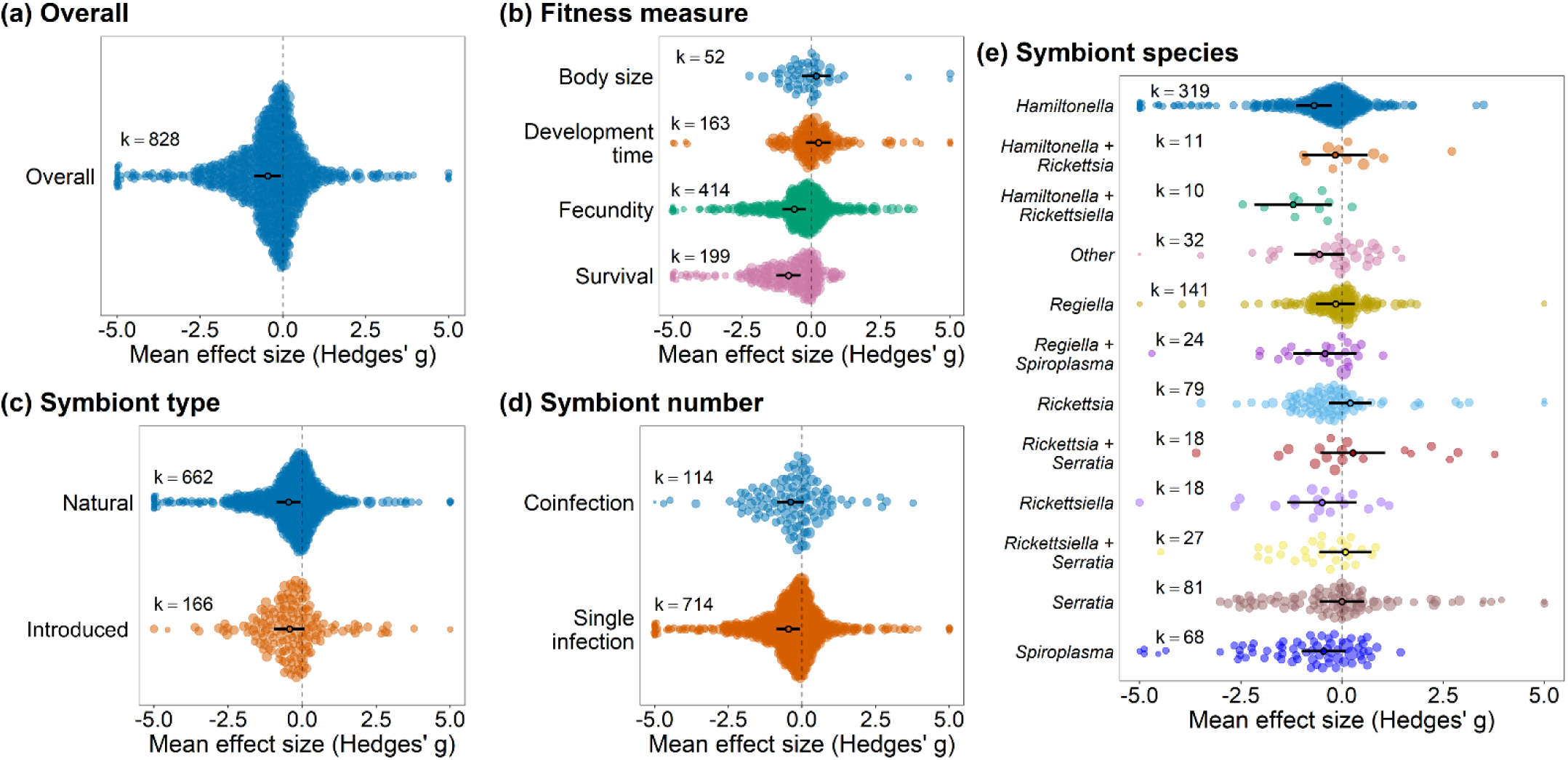
Sub-analysis: Aphids only. Effect of defensive symbionts on the fitness of aphids that are in the absence of natural enemies. (a) overall model, (b) model including fitness measure as moderator, (c) model including symbiont type (i.e., if the symbiont naturally infects the host species or if it was artificially introduced) as moderator, (d) model including symbiont number as moderator, and (e) model including symbiont species as moderator. Positive values indicate positive effects on host fitness. Hence, positive values indicate higher fecundity, survival, body size and shorter development time. Negative values indicate that symbionts have a negative effect on fitness. Points are the weighted mean effect sizes ± 95% confidence intervals. Each colored circle represents an effect size, scaled according to precision (inverse of the sampling error variance). ‘k’ is the number of effect sizes. Fourteen points fell below −5 and four points fell above +5; for visualization purposes only, these points are shown at ±5. All data points were included in the statistical analyses.

**Figure S10.**
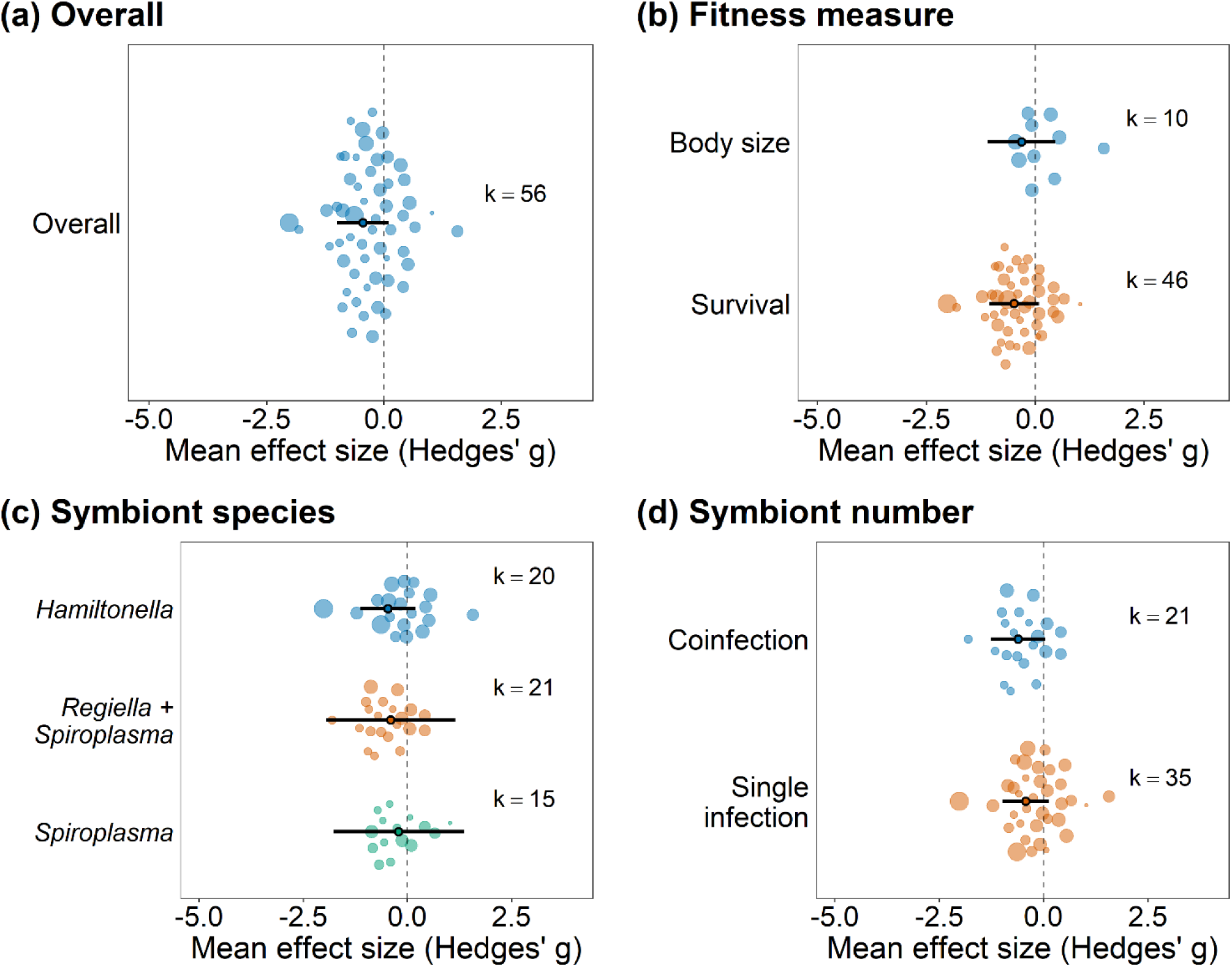
Sub-analysis: Aphids only. Effect of defensive symbionts on the fitness of natural enemies infecting symbiont-infected hosts. (a) overall model, (b) model including fitness measure as moderator, (c) model including symbiont species as moderator, and (d) model including symbiont number as moderator. Positive values indicate positive effects on host fitness. Hence, positive values indicate bigger body size and higher survival. Points are the weighted mean effect sizes ± 95% confidence intervals. Each colored circle represents an effect size, scaled according to precision (inverse of the sampling error variance). ‘k’ is the number of effect sizes.

## Supplementary tables

**Table S1.**
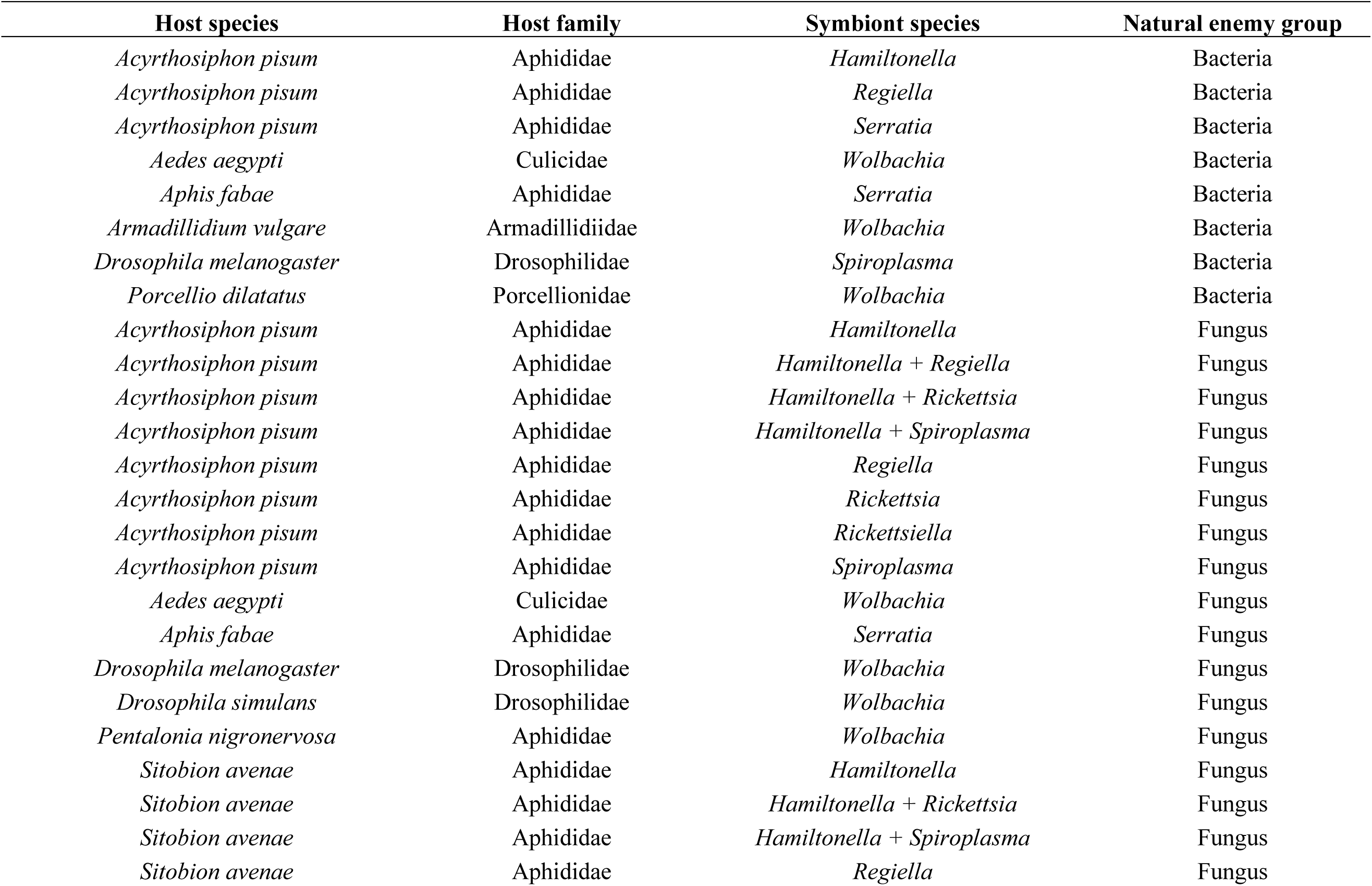
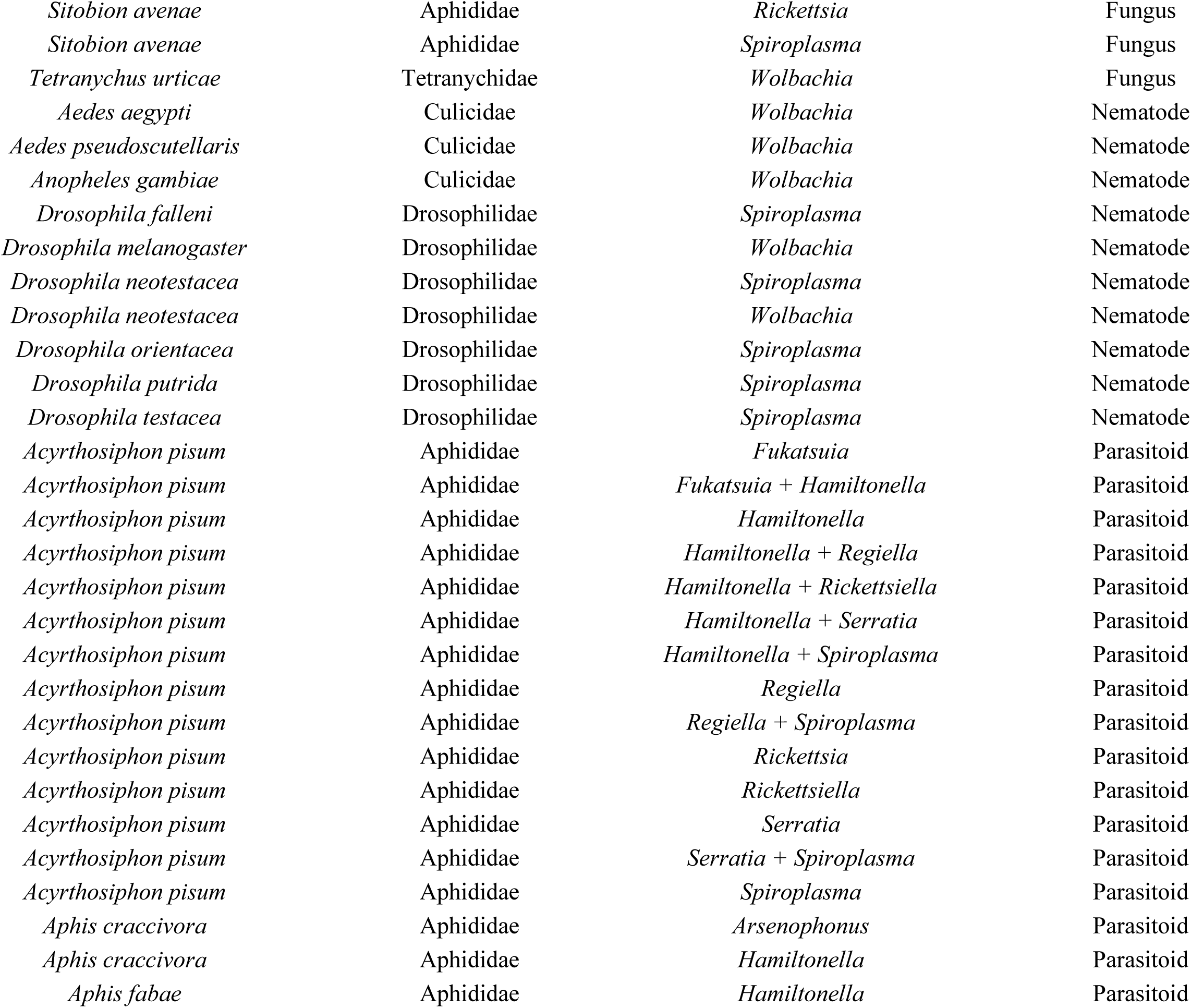
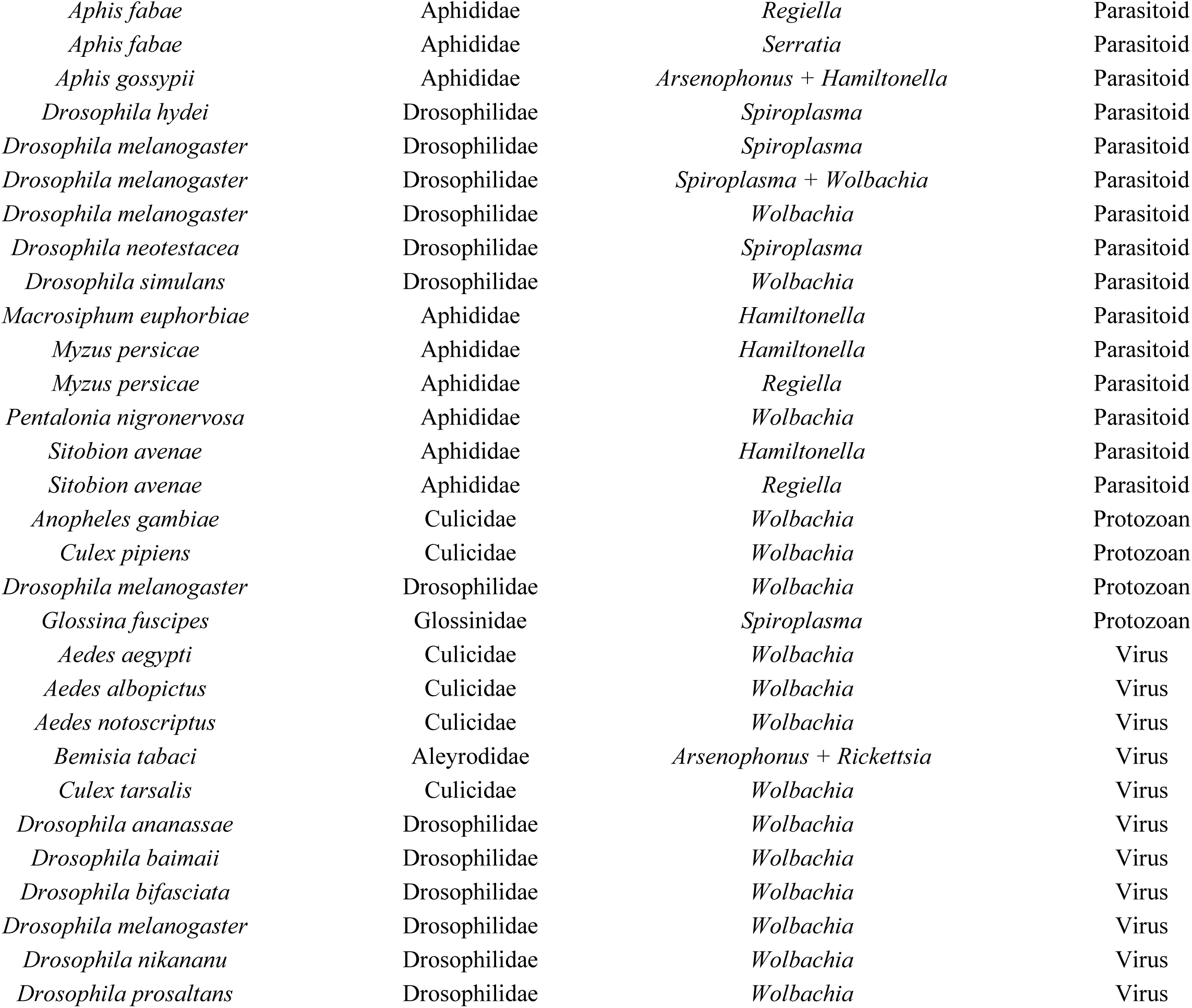
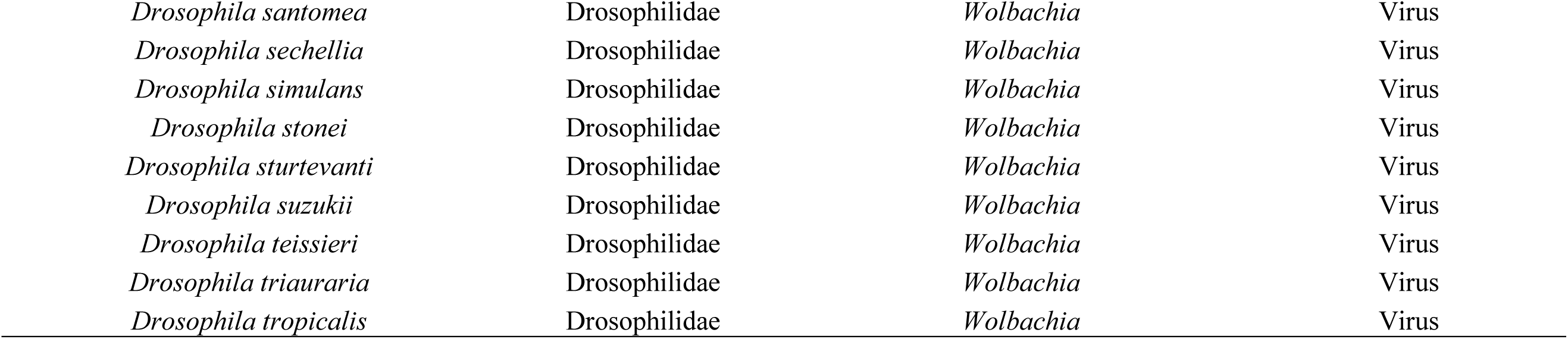
Study systems from “protection” dataset that were included in the meta-analysis.

**Table S2.**
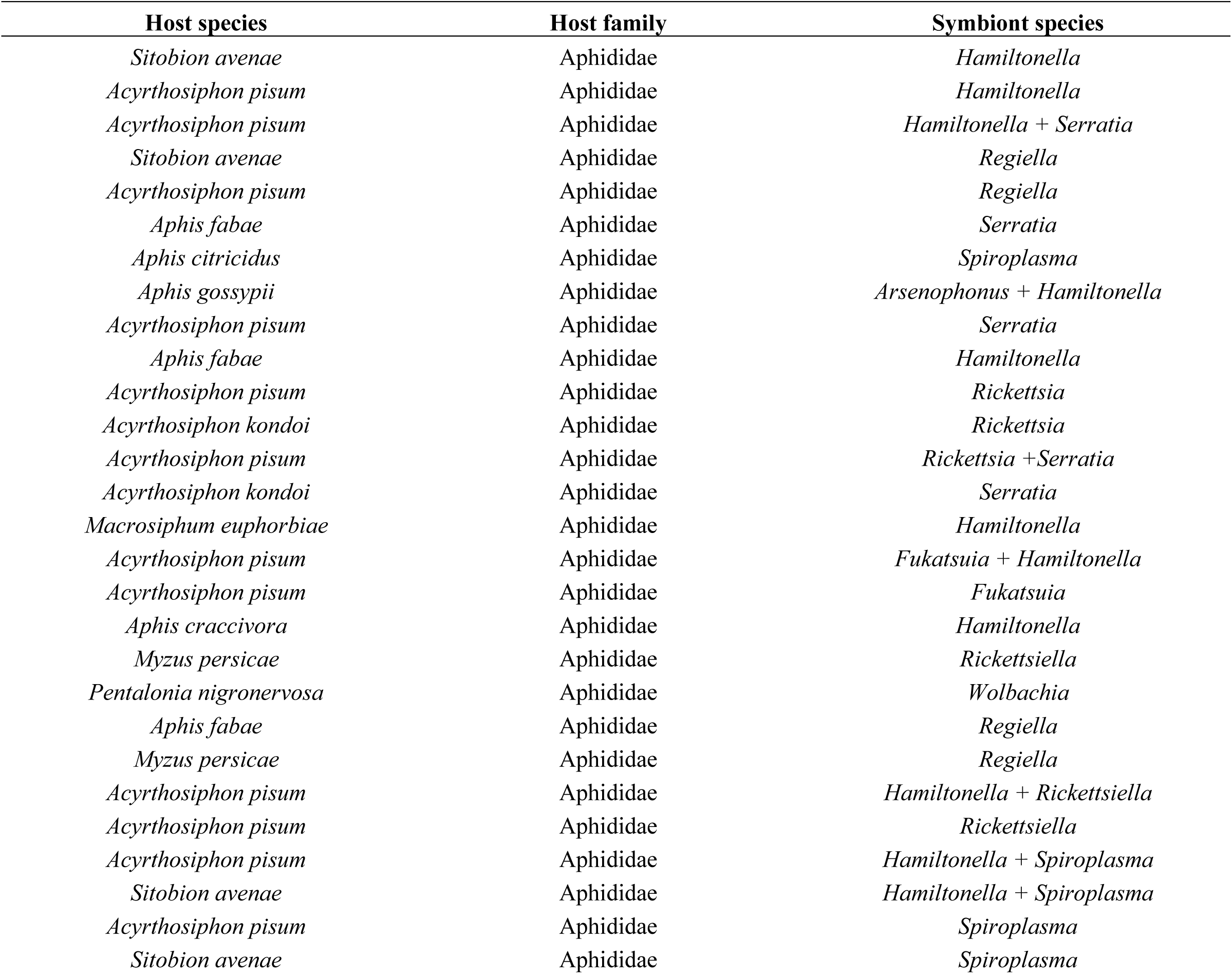
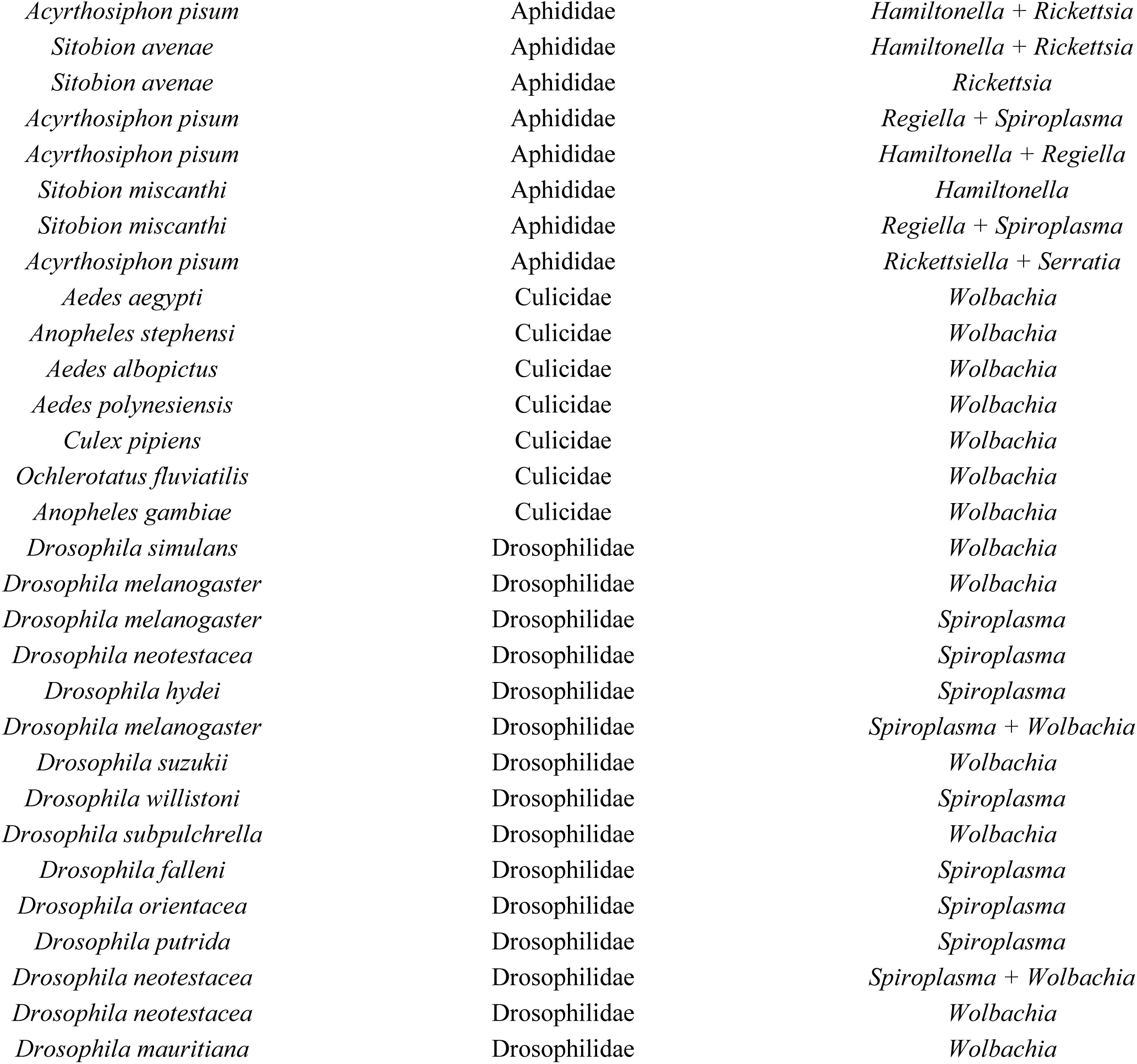
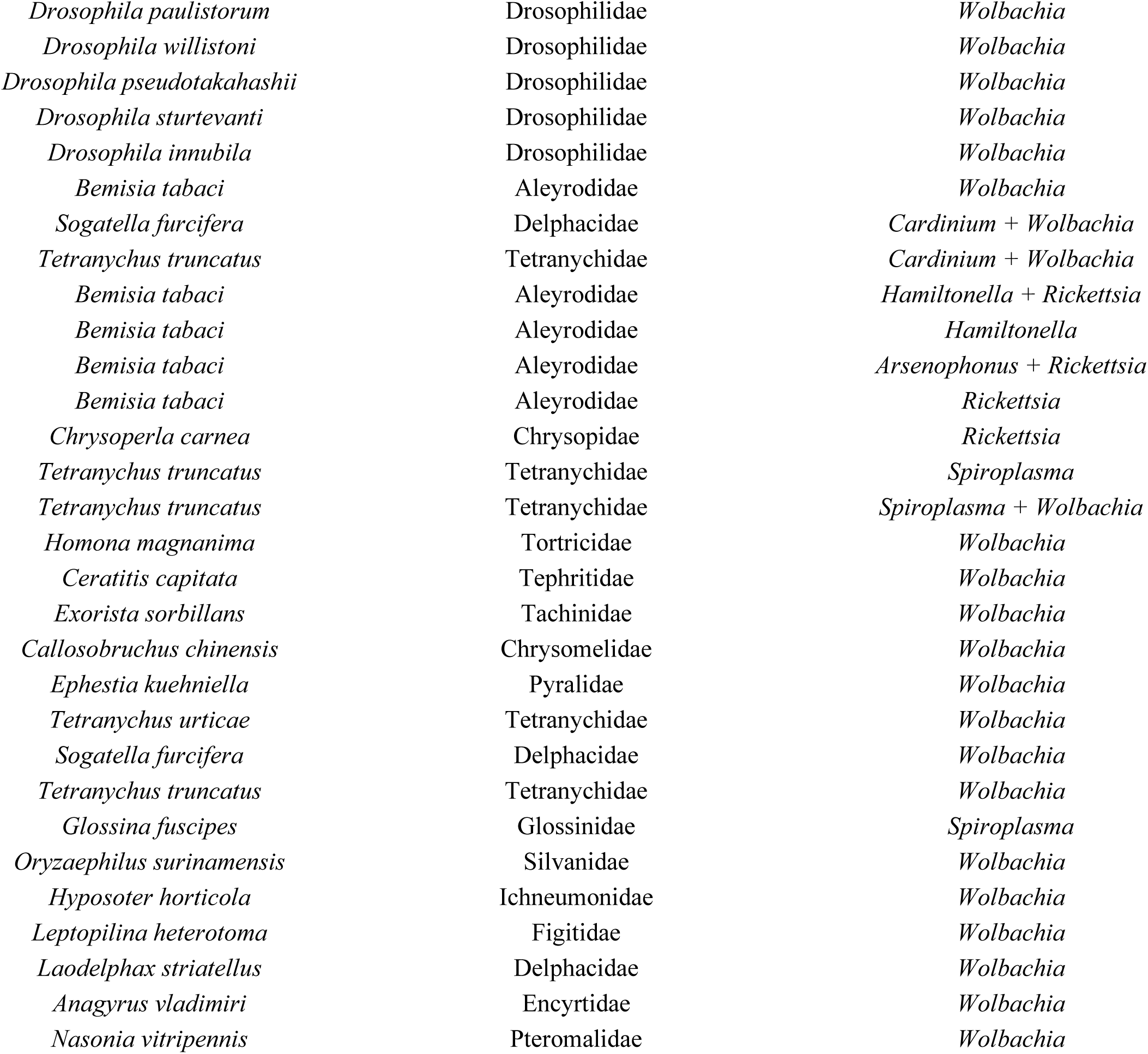
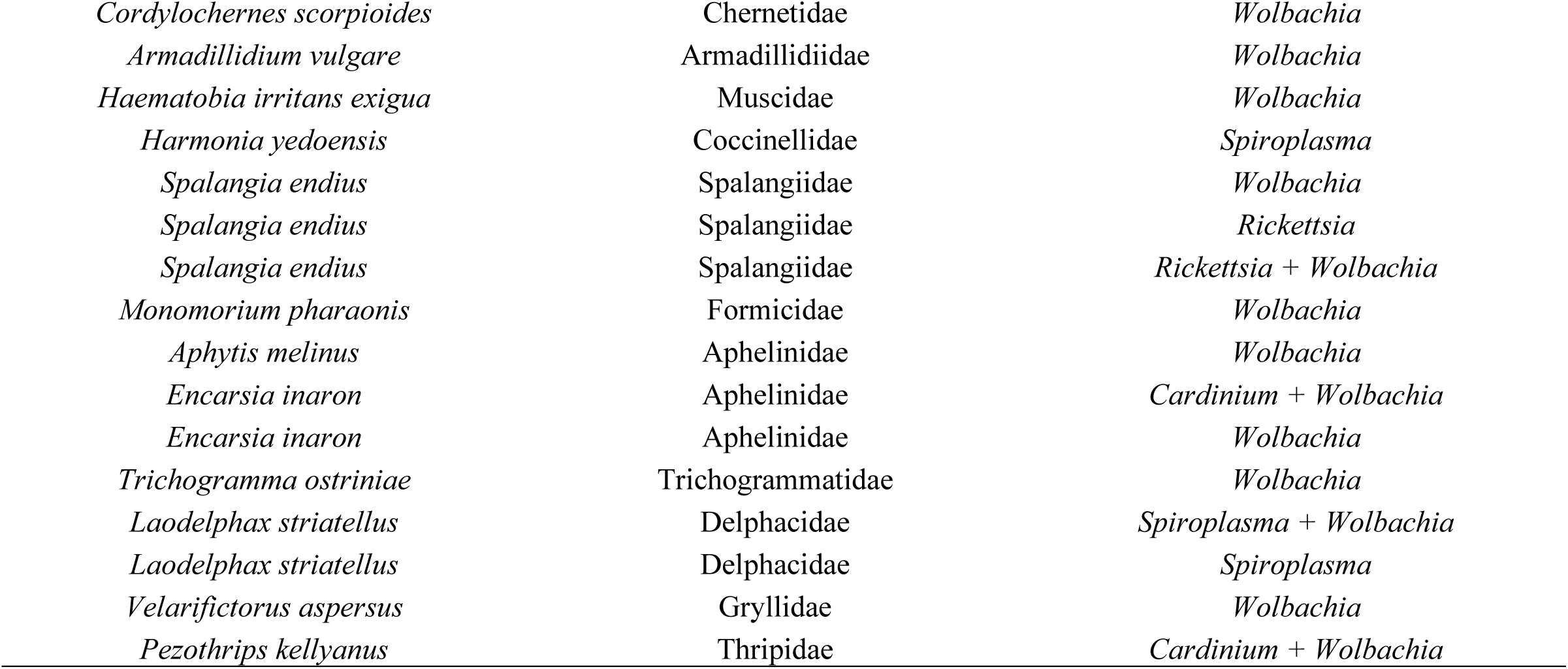
Study systems from “cost to hosts” dataset that were included in the meta-analysis.

**Table S3.**
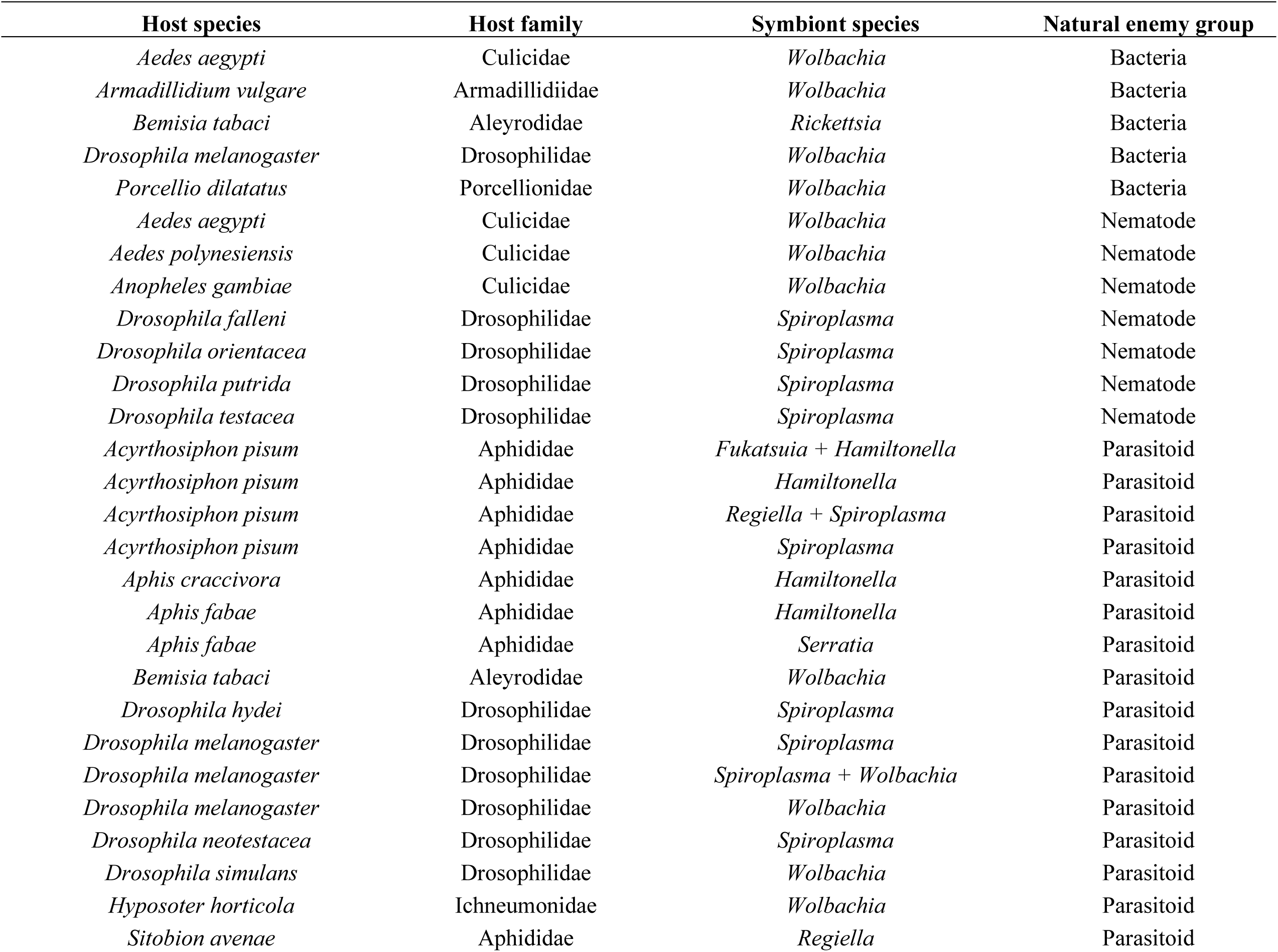
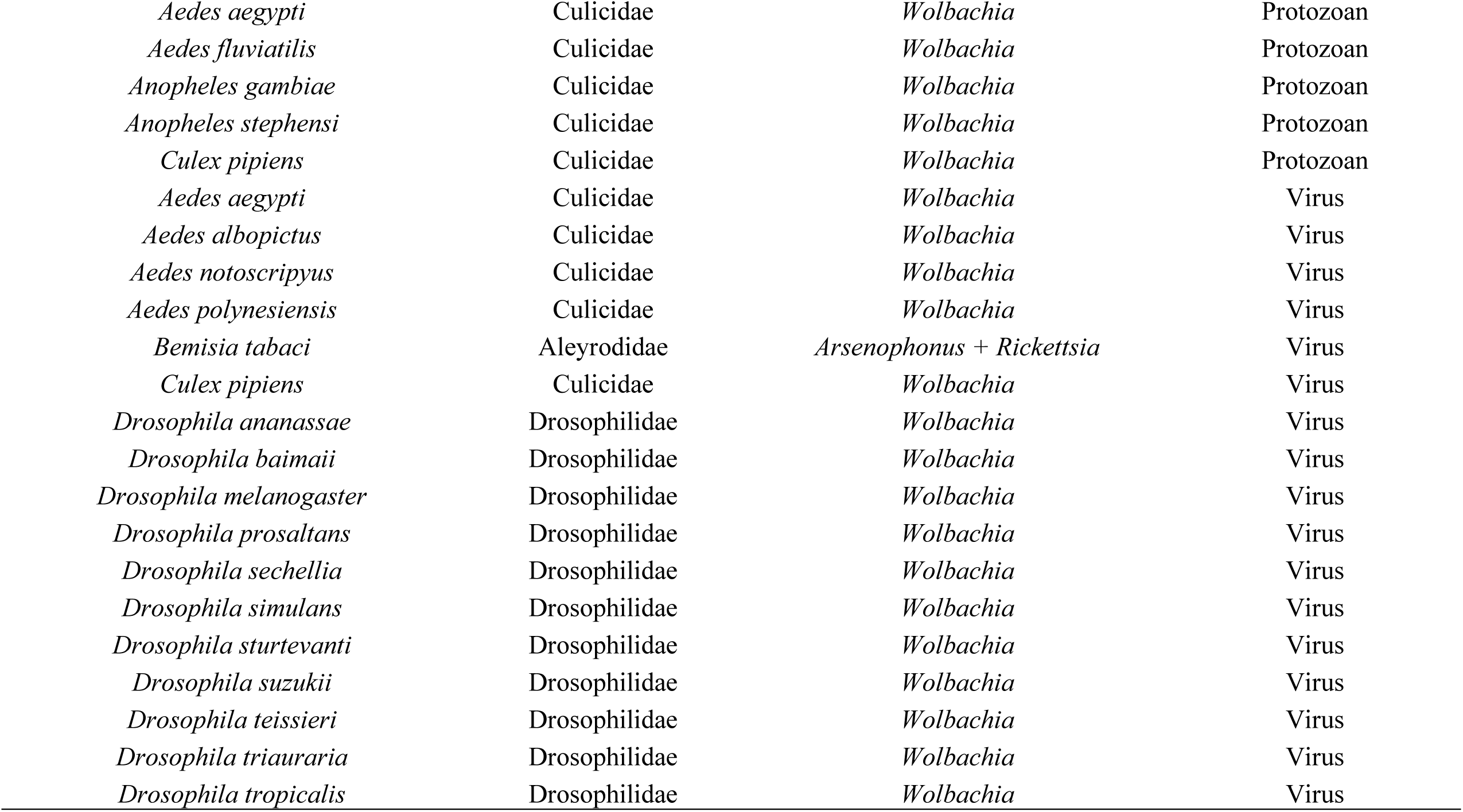
Study systems from “cost to natural enemy” dataset that were included in the meta-analysis.

**Table S4.**
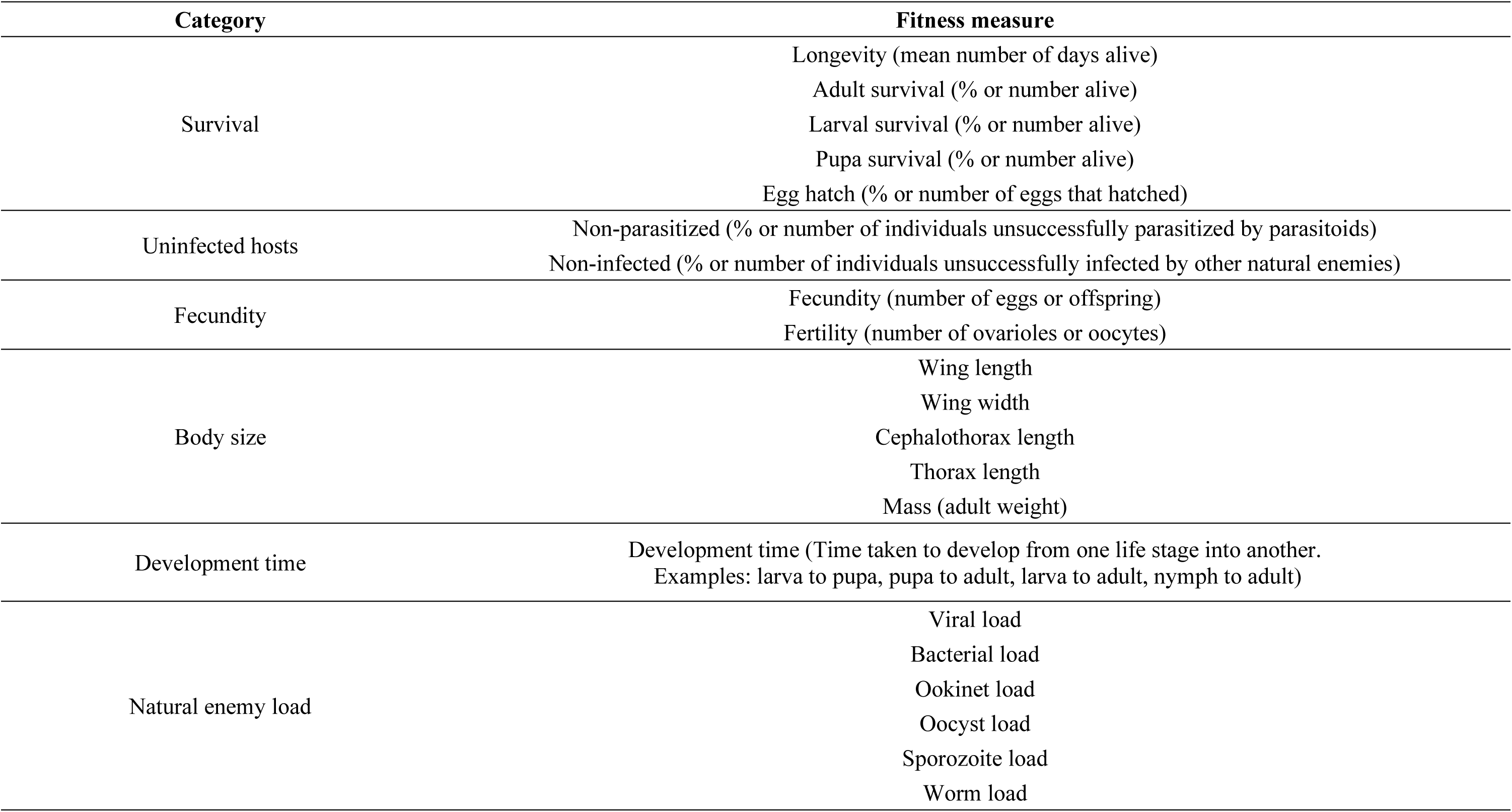
List of all fitness measures extracted from the studies to calculate the mean effect sizes. Measures are grouped in six broad categories.

**Table S5.**
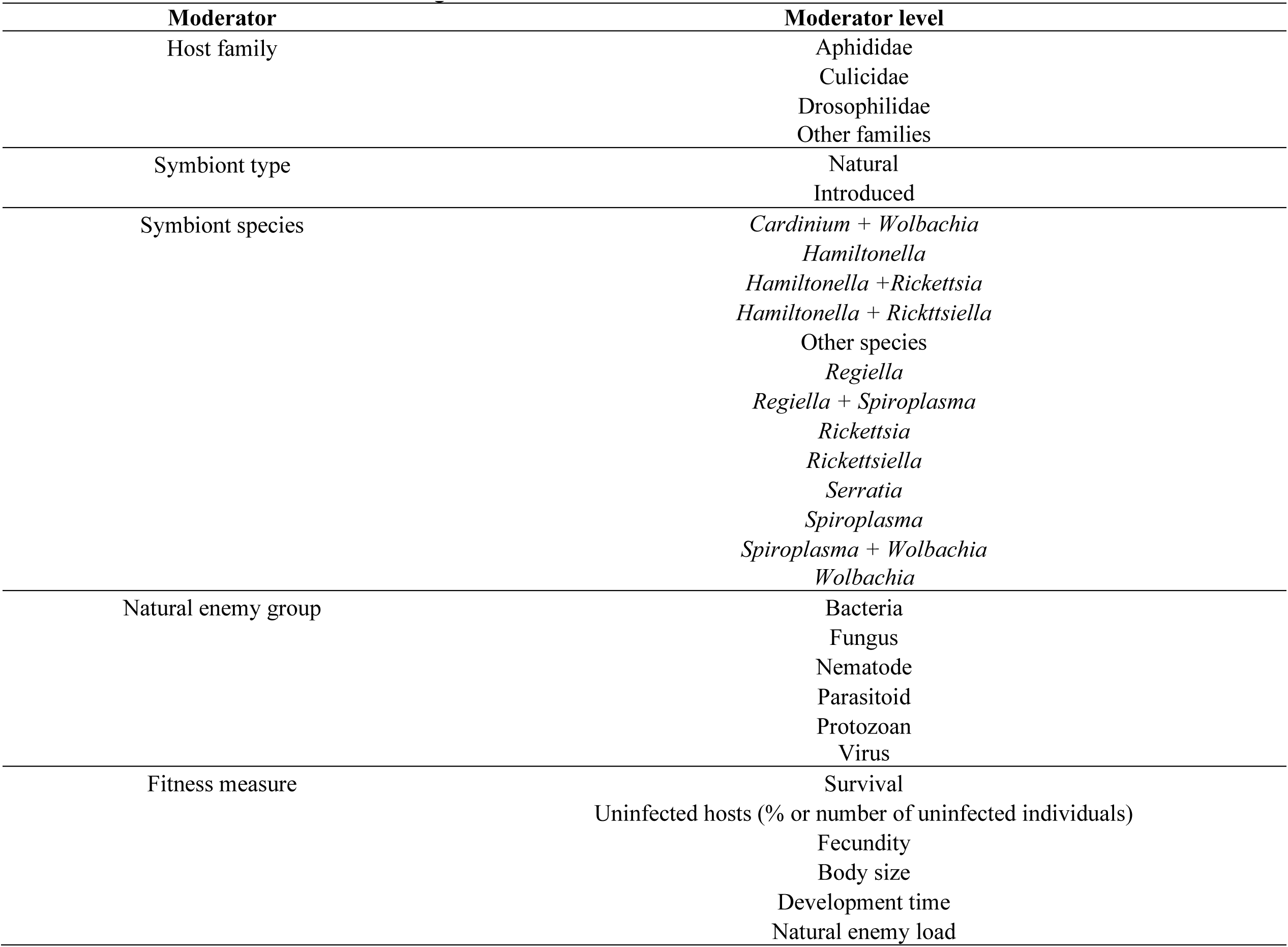
Variables collected from the original studies to be used as moderators in the models.

**Table S6.**
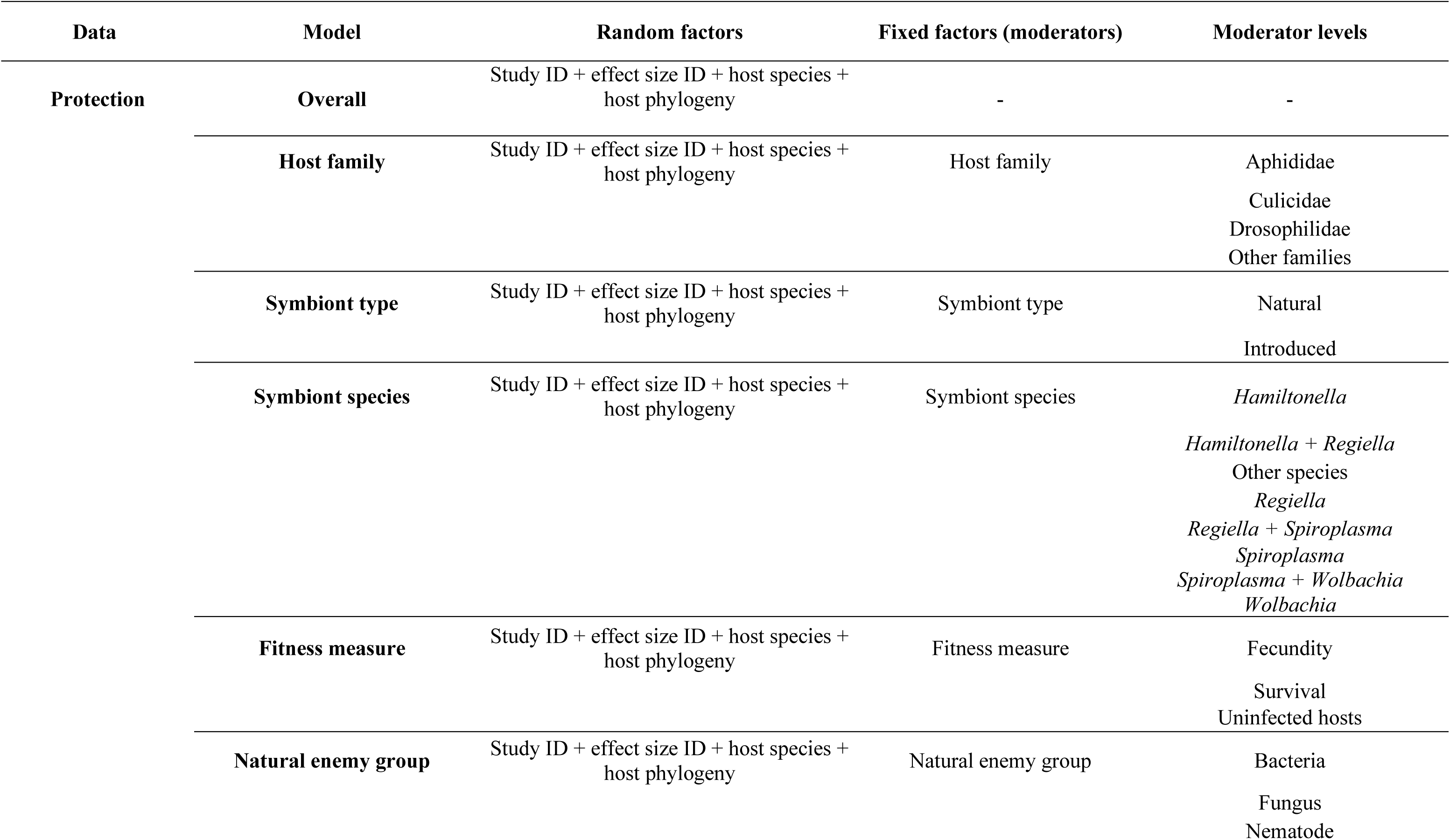
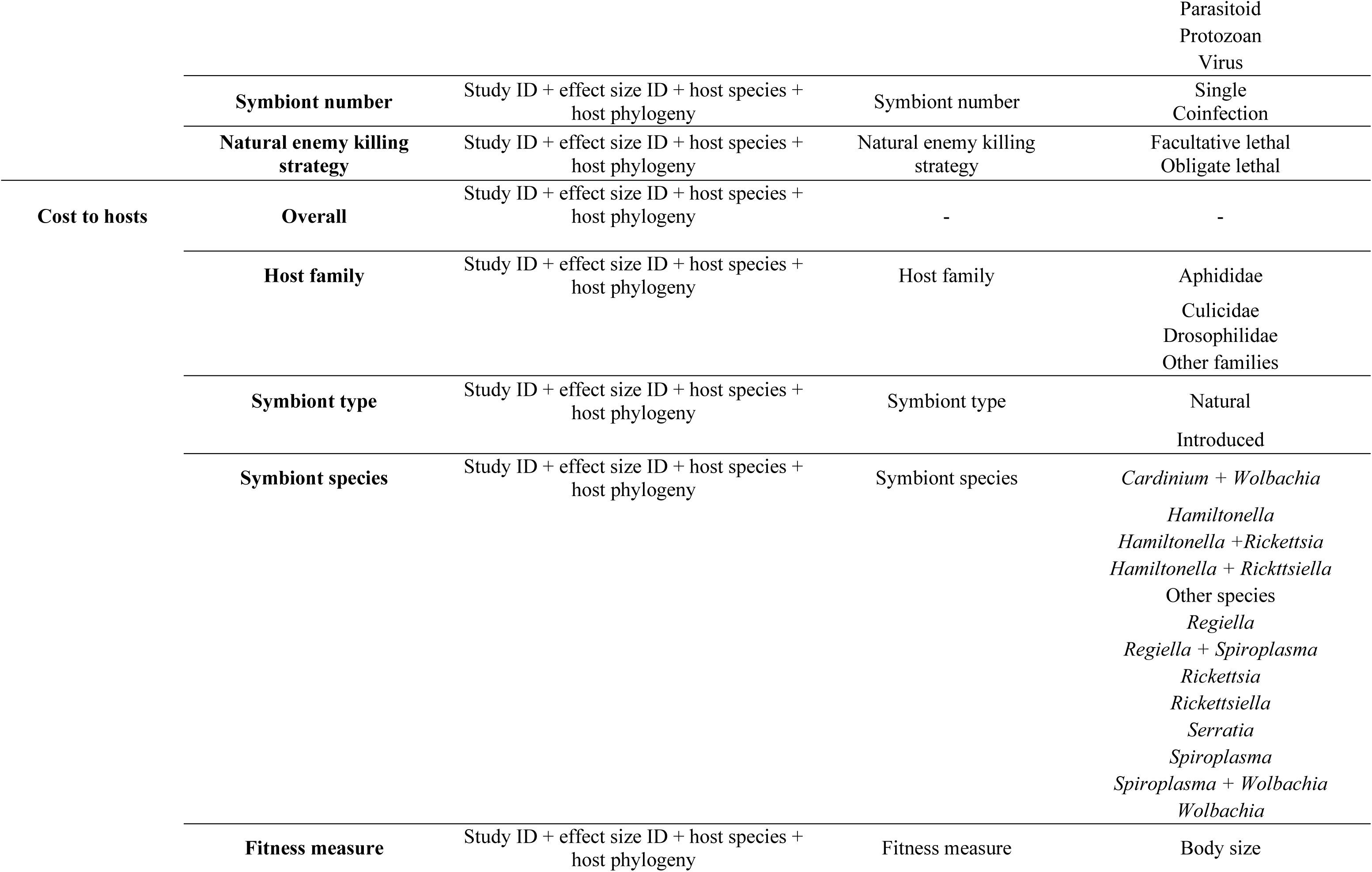
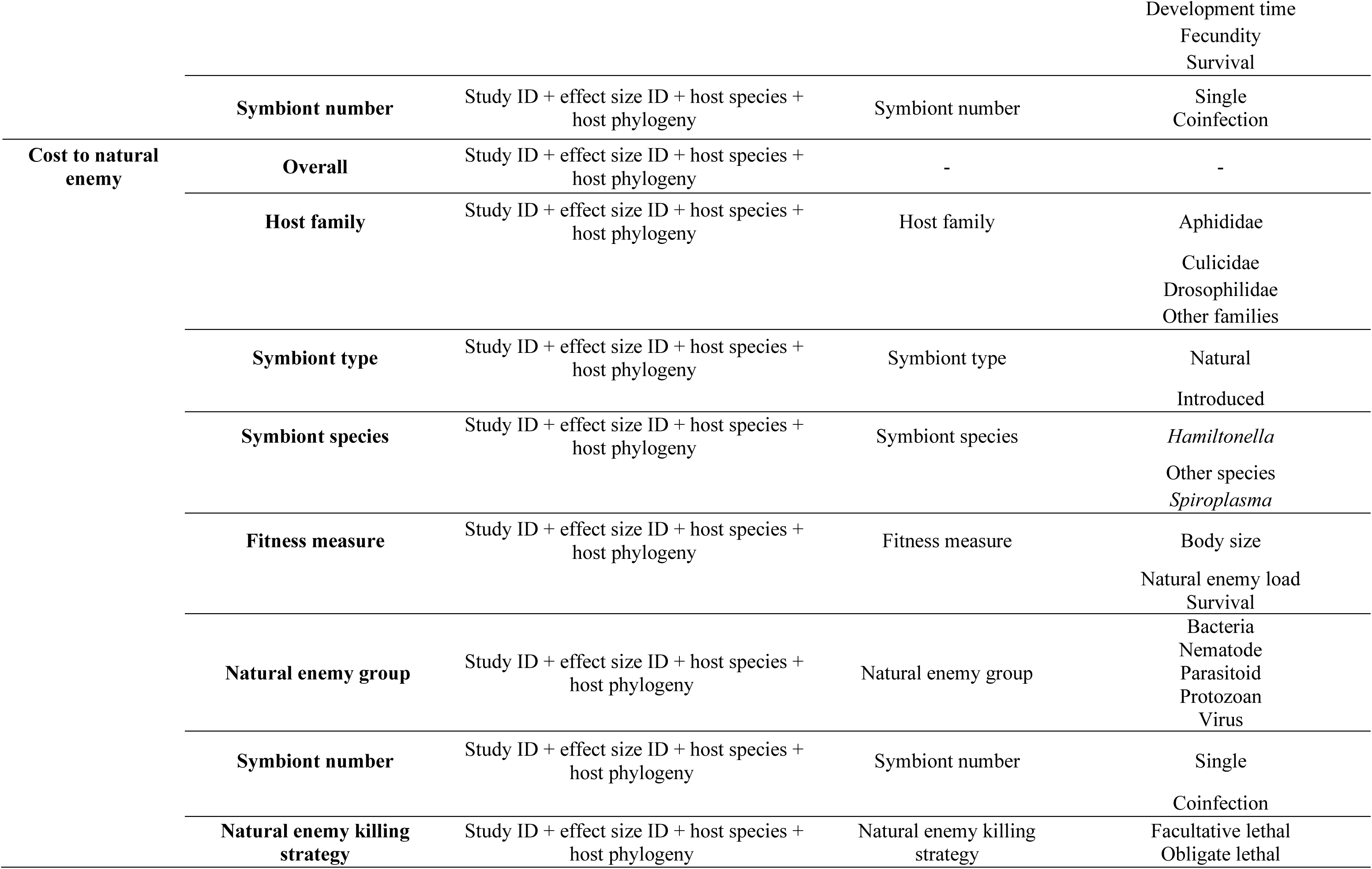
Main analysis: multilevel meta-analytical models performed in this study, specifying their random and fixed factors (moderators) and moderator levels.

**Table S7.**
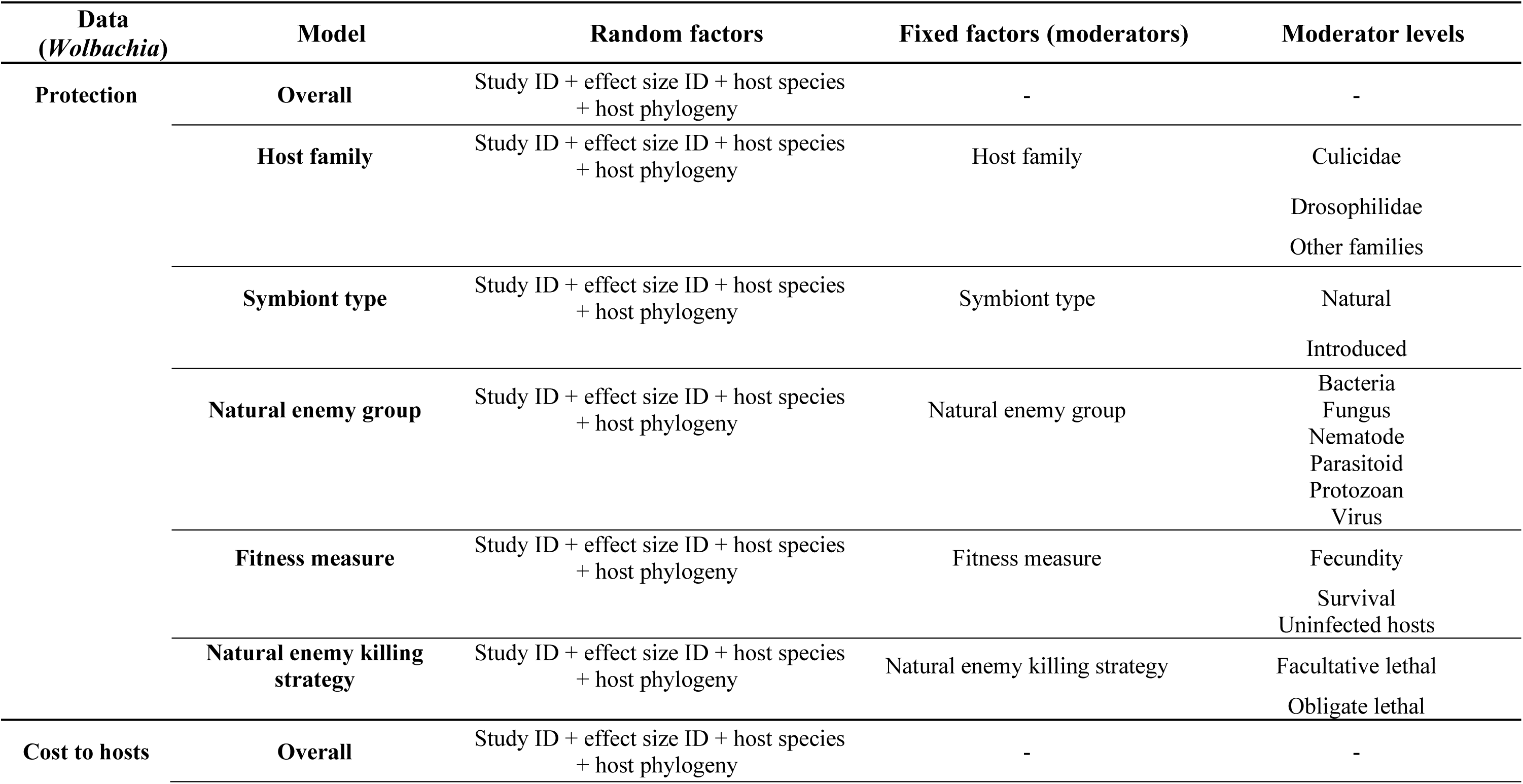
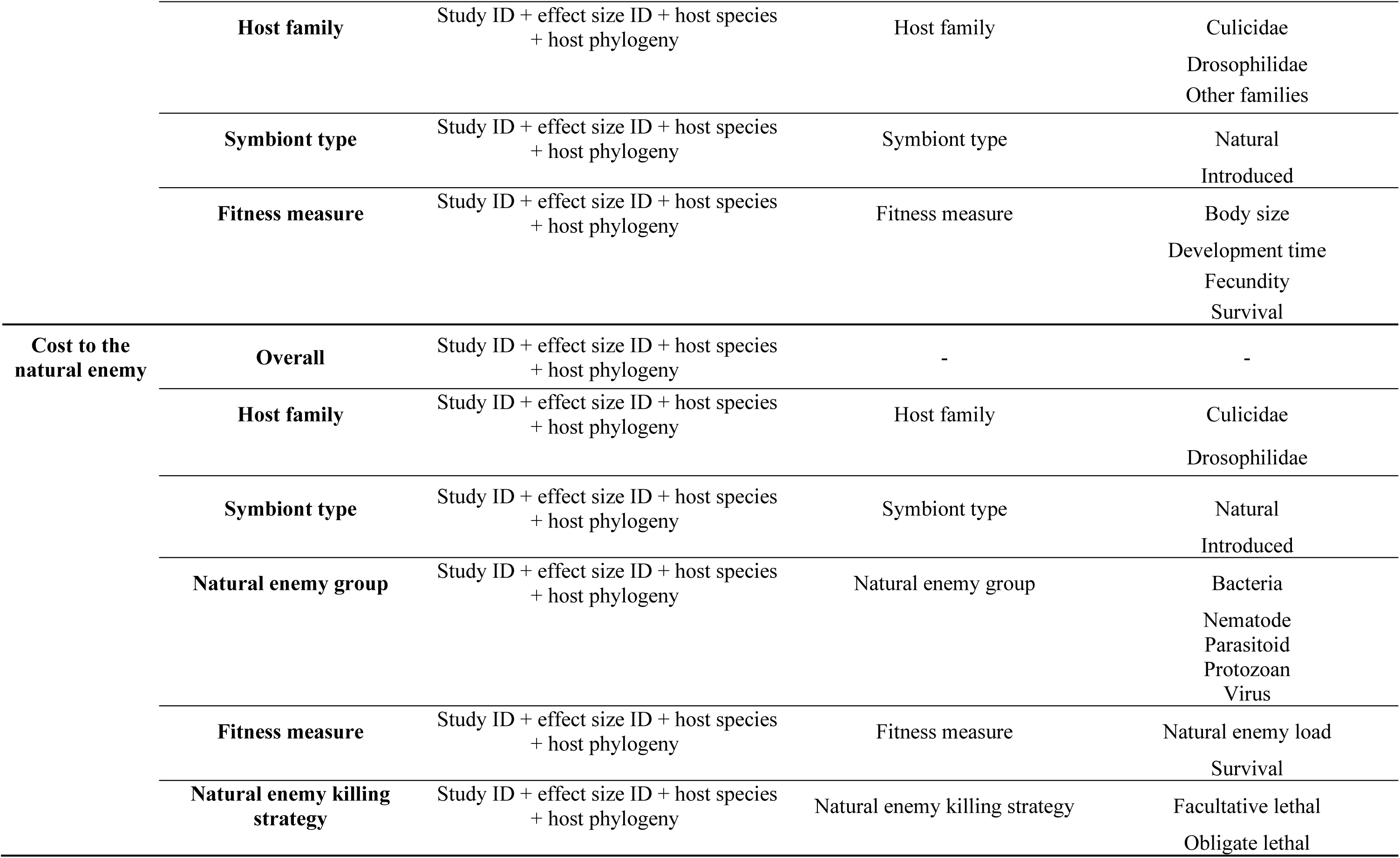
Subgroup analysis: analyses performed only with *Wolbachia* data. Multilevel meta-analytical models performed in this study, specifying their random and fixed factors (moderators) and moderator levels.

**Table S8.**
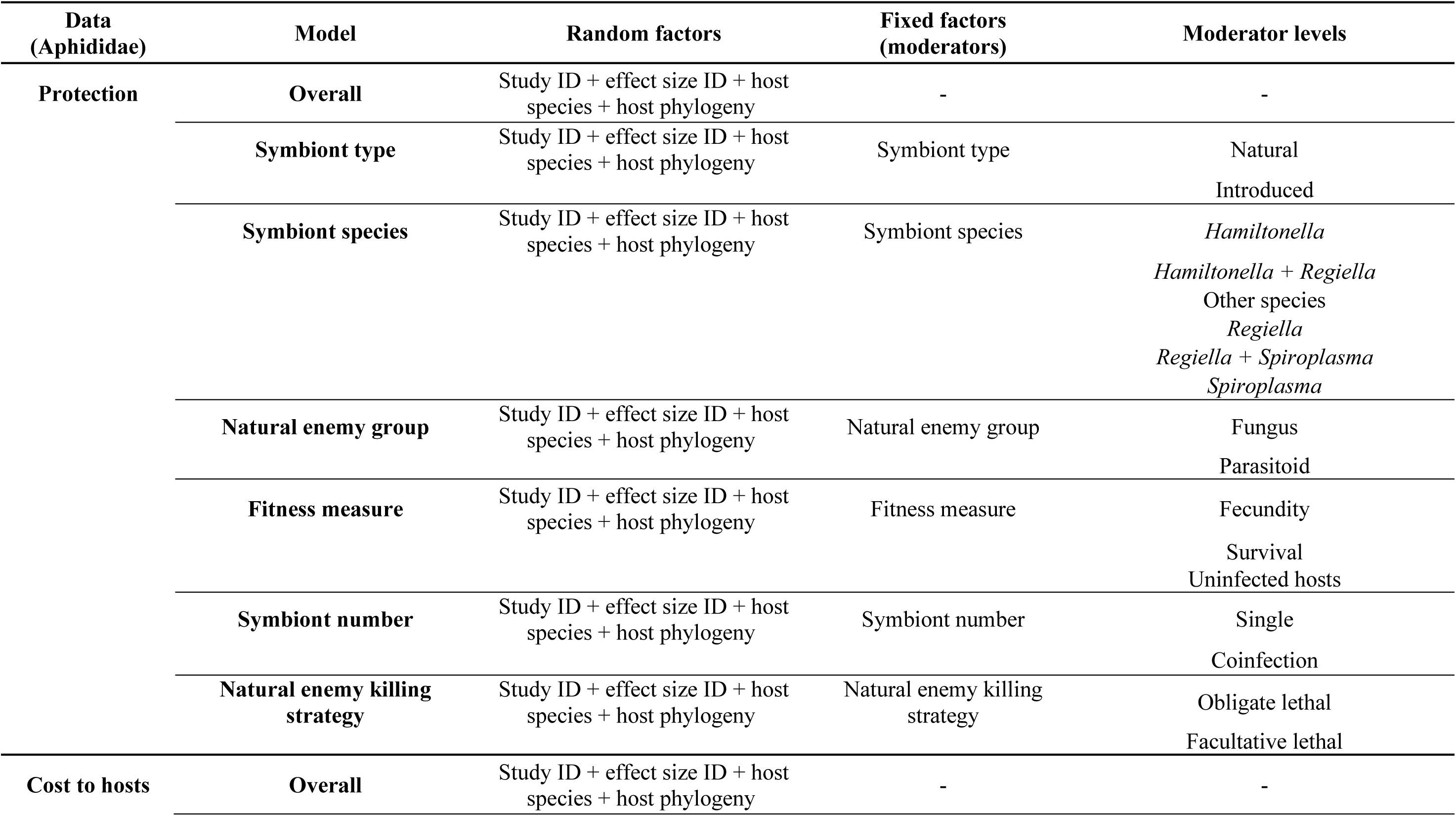
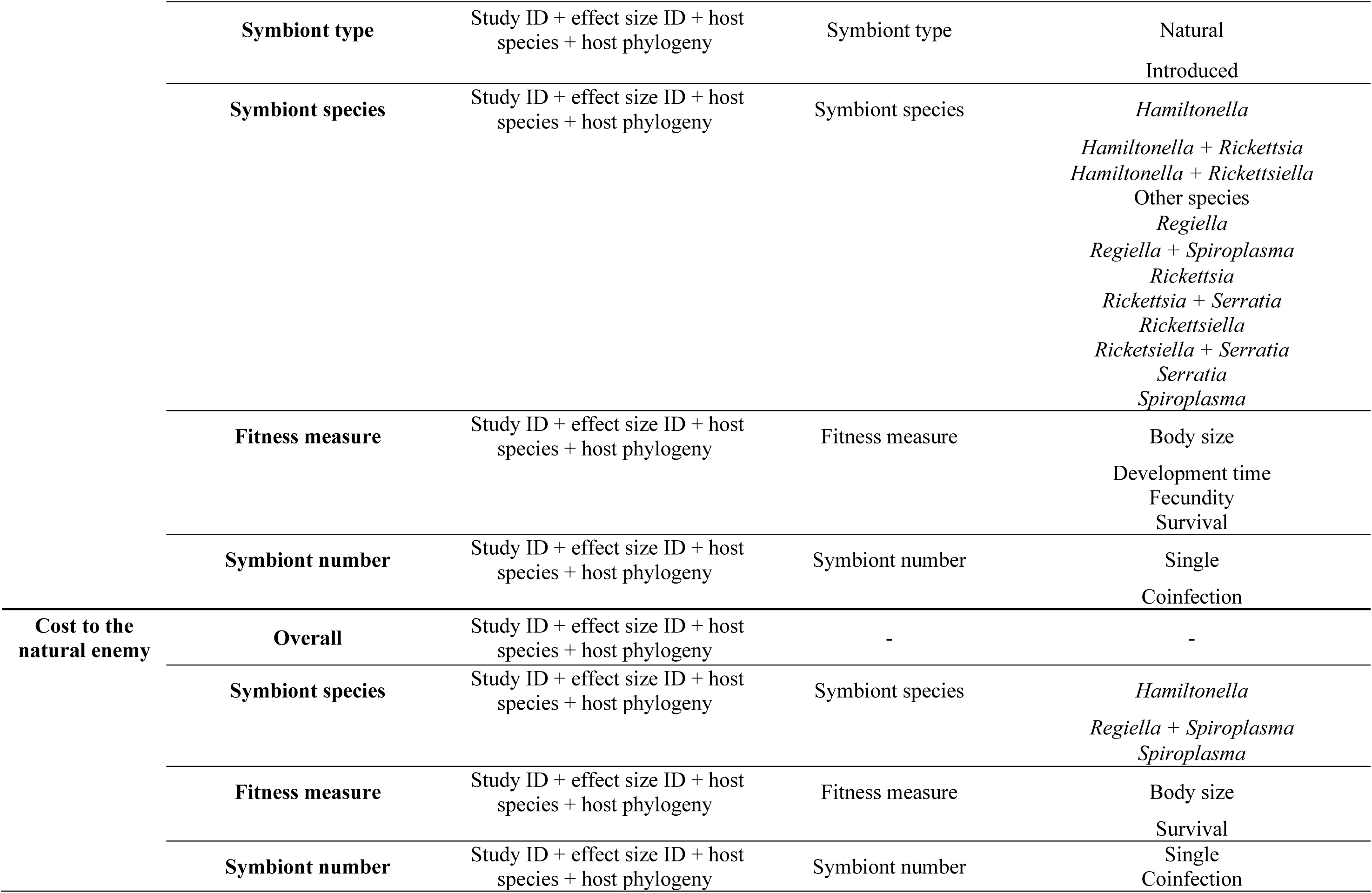
Subgroup analysis: analyses performed only with Aphididae data. Multilevel meta-analytical models performed in this study, specifying their random and fixed factors (moderators) and moderator levels. Moderators “symbiont type” and “natural enemy group” were not included in the cost to the natural enemy data analysis because they presented only one moderator level (“natural” and “parasitoid”, respectively).

**Table S9.**
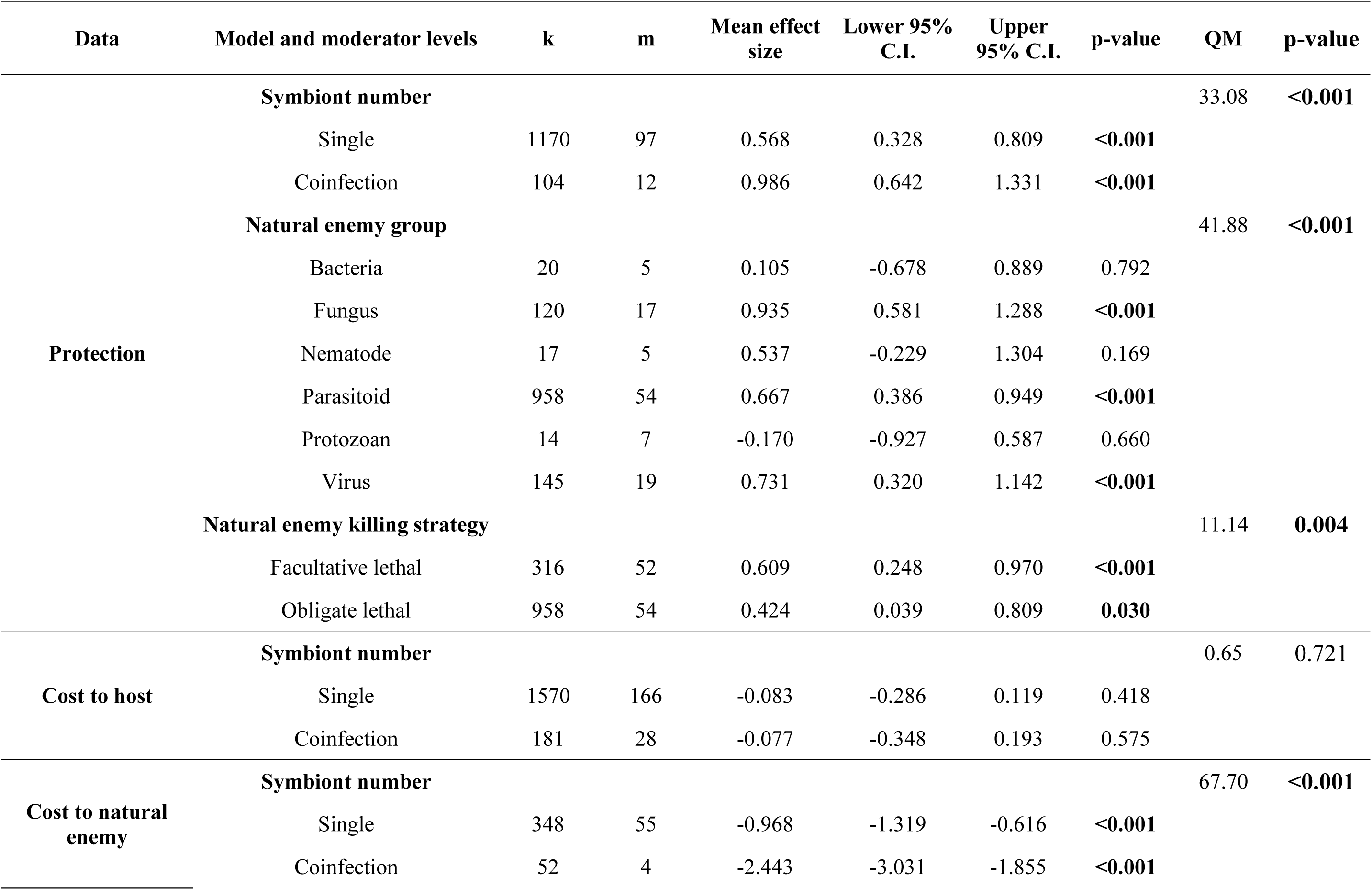
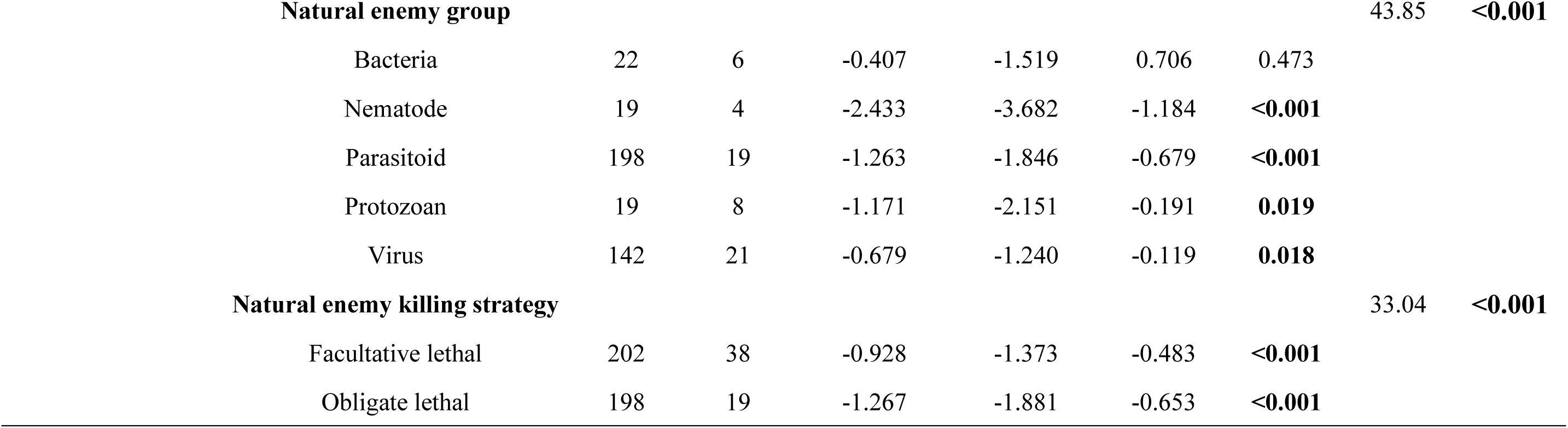
Main analysis: results of the meta-analyses on the costs and benefits of hosting defensive symbionts including symbiont number, natural enemy group and natural enemy killing strategy as moderators.

**Table S10.**
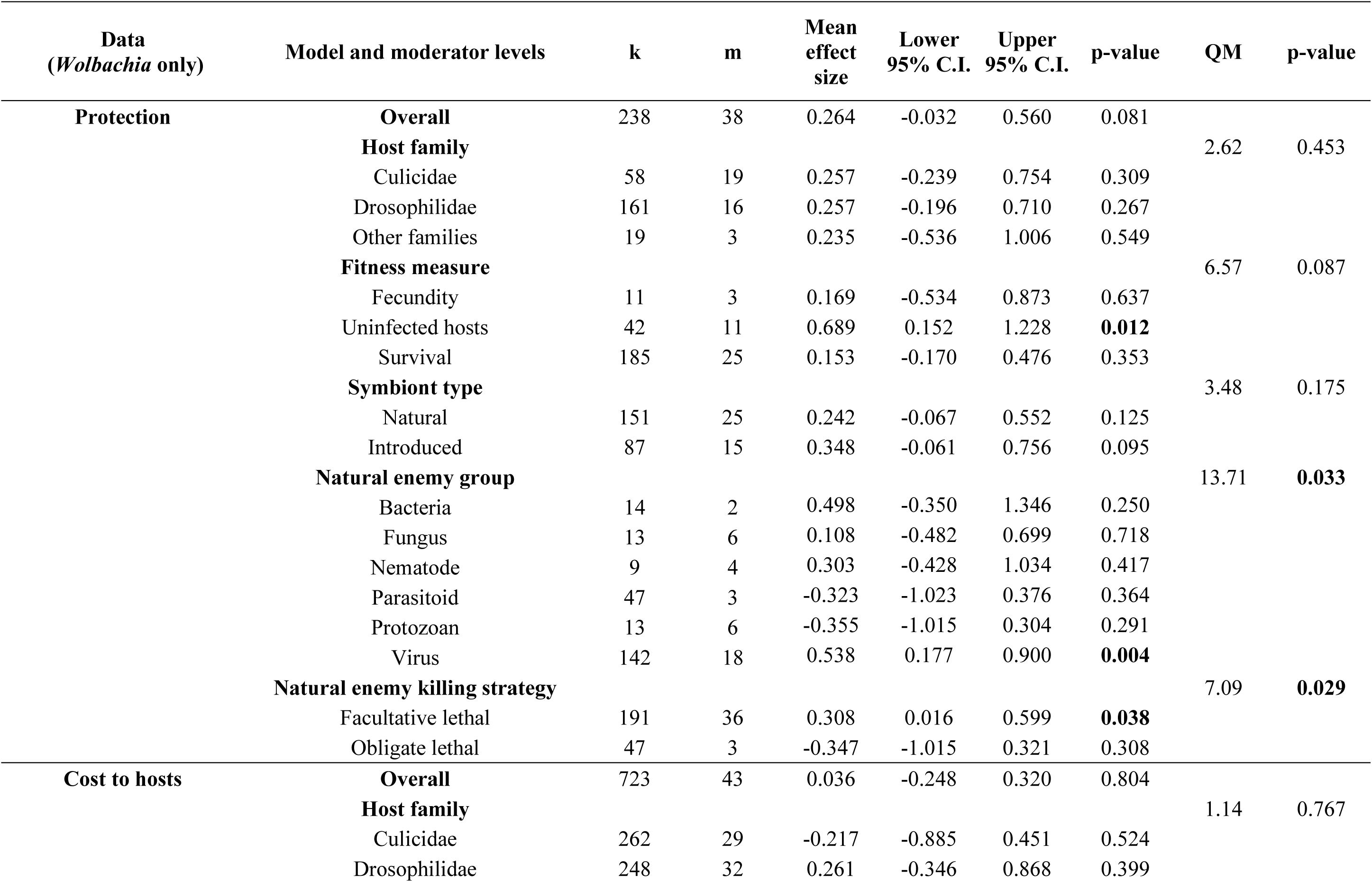
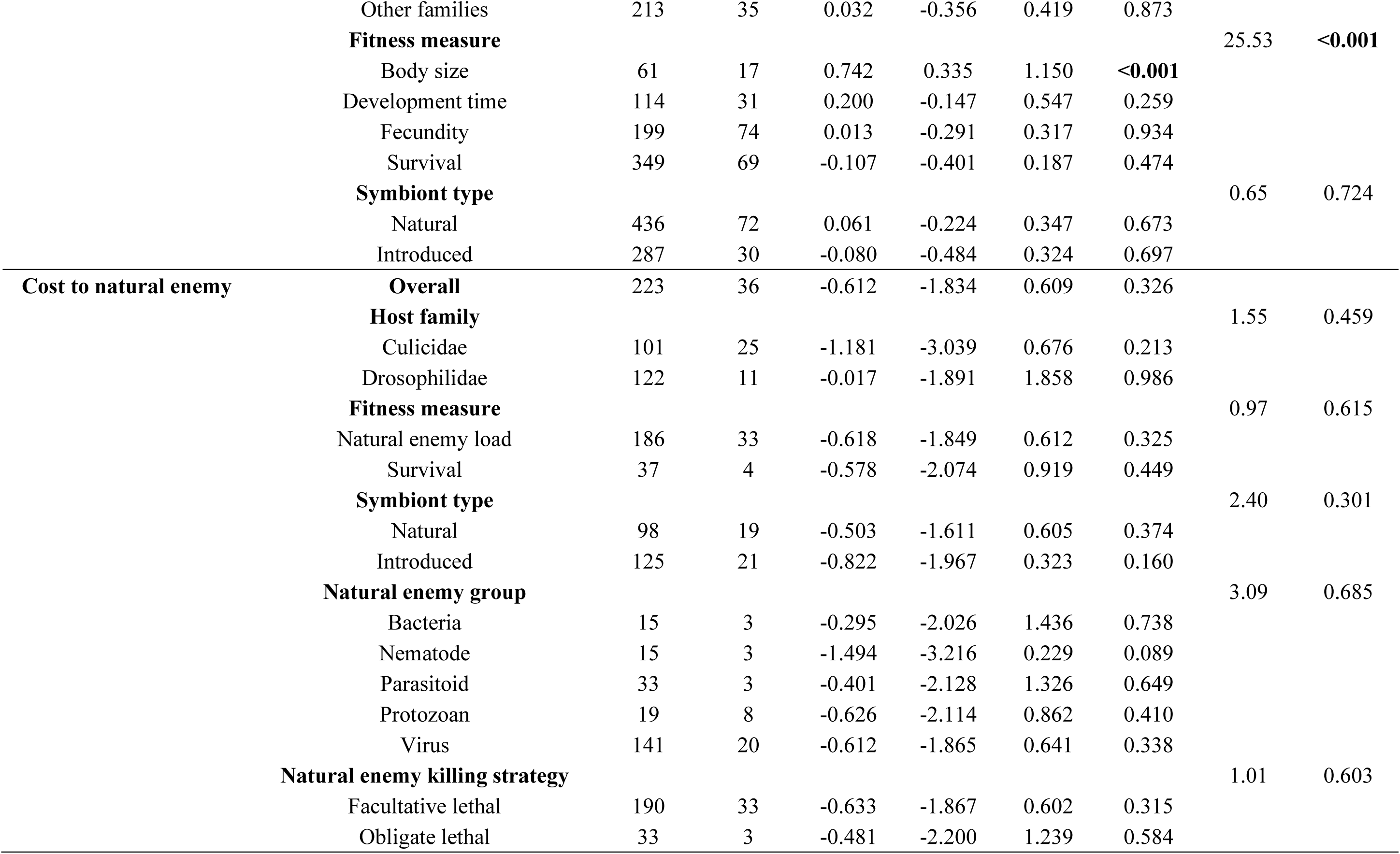
Subgroup analysis: results of the analyses including only effect sizes from hosts harboring bacterial symbiont *Wolbachia*.

**Table S11.**
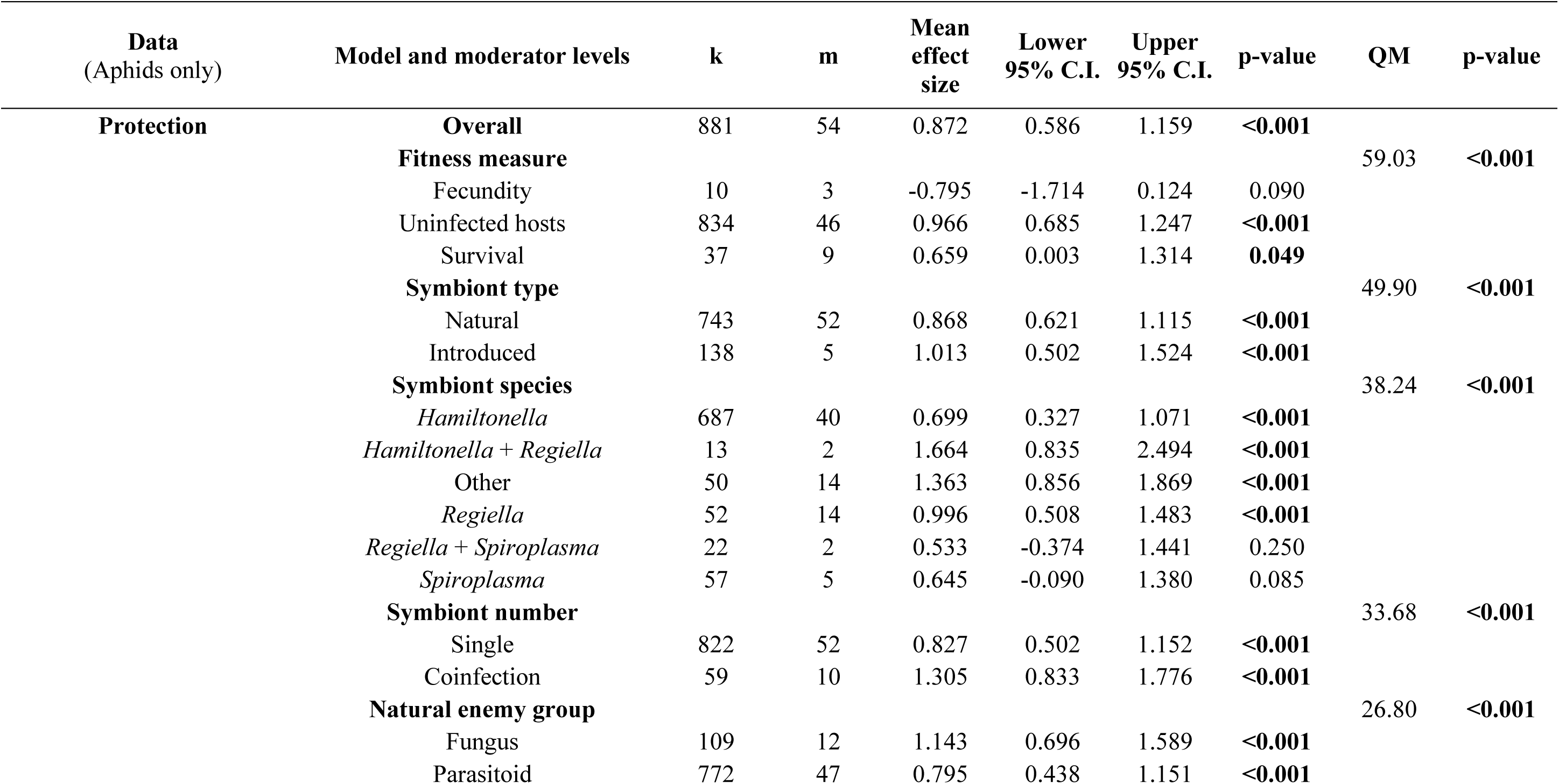
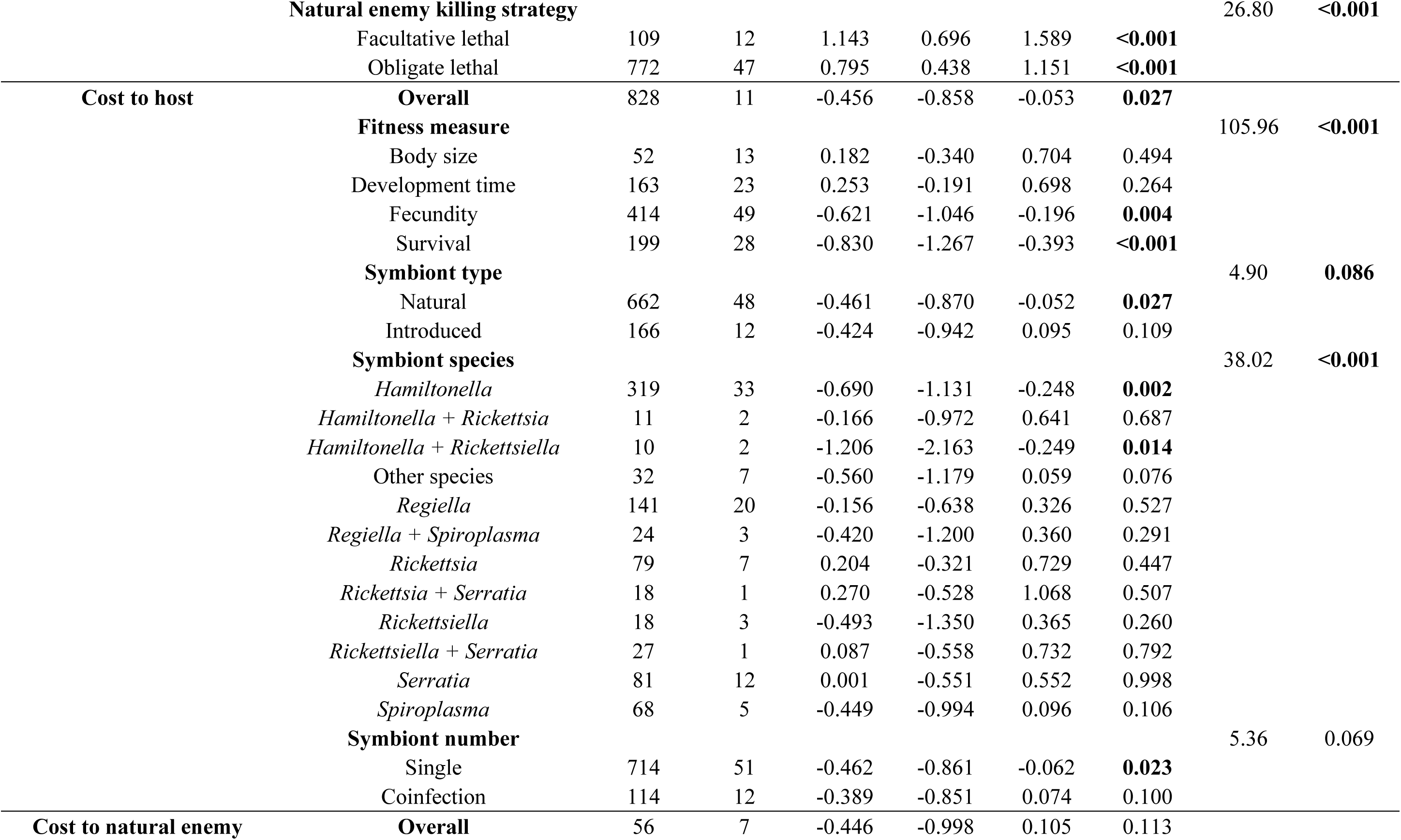
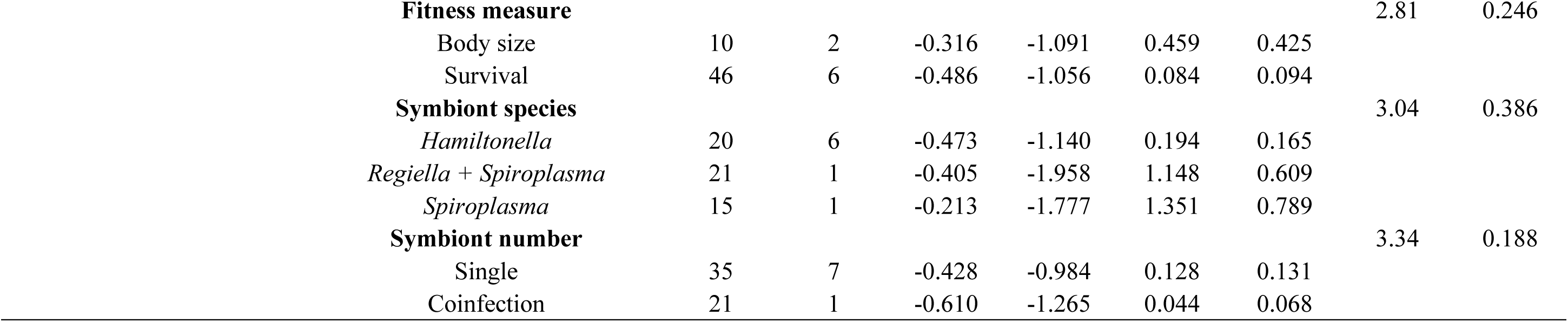
Subgroup analysis: results of the analyses including only effect sizes from hosts from the Aphididae family. Moderators “symbiont type” and “natural enemy group” were not included in the cost to the natural enemy data analysis because they presented only one moderator level (“natural” and “parasitoid”, respectively).

**Table S12.**
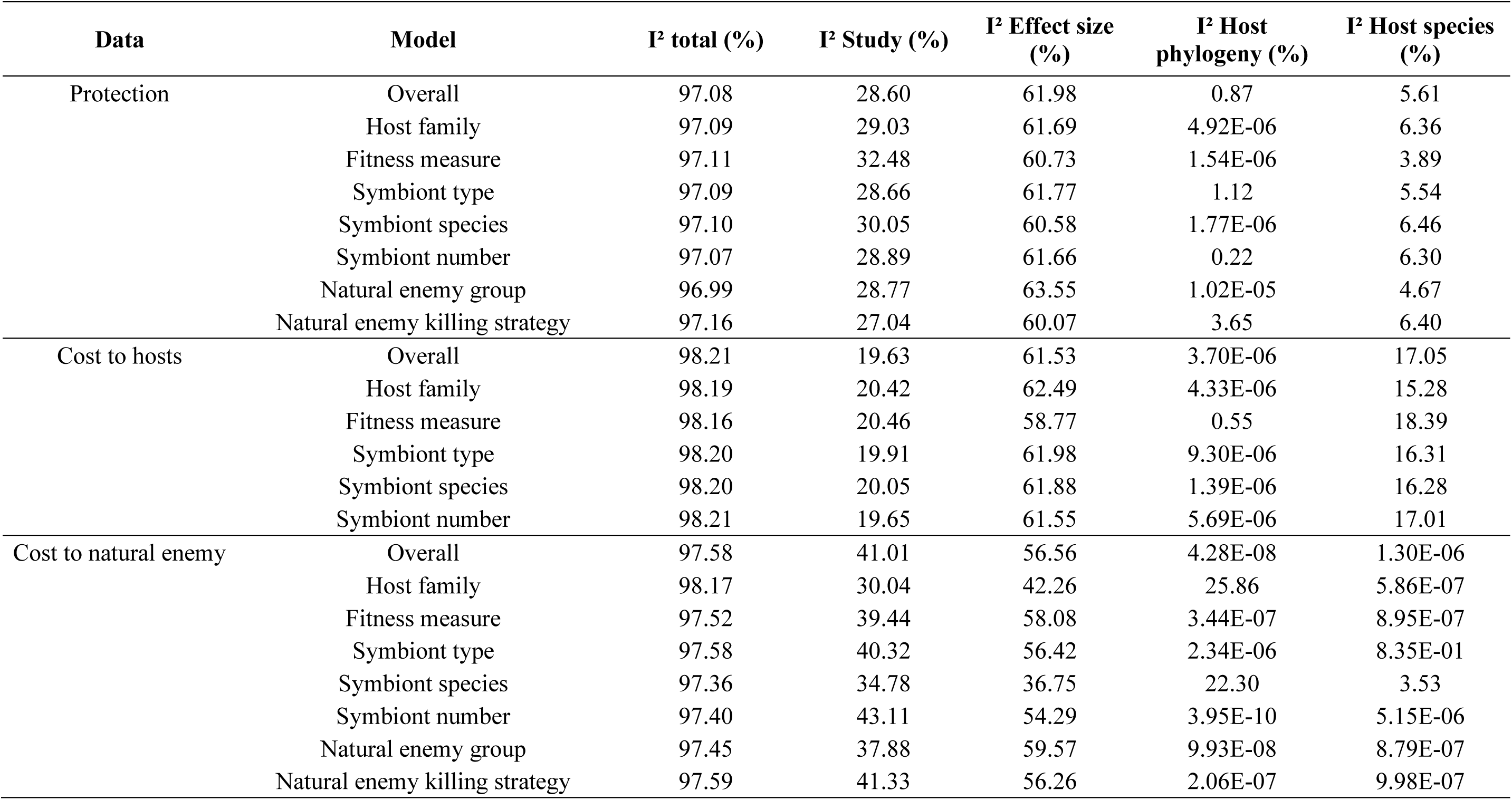
Main analysis: results of the heterogeneity test (I^2^) for each meta-analytical model. For each model, we quantified the total amount of heterogeneity (I^2^) and the percentage of contribution of each random effect for the variance in the models: study, effect size, host phylogeny and host species.

**Table S13.**
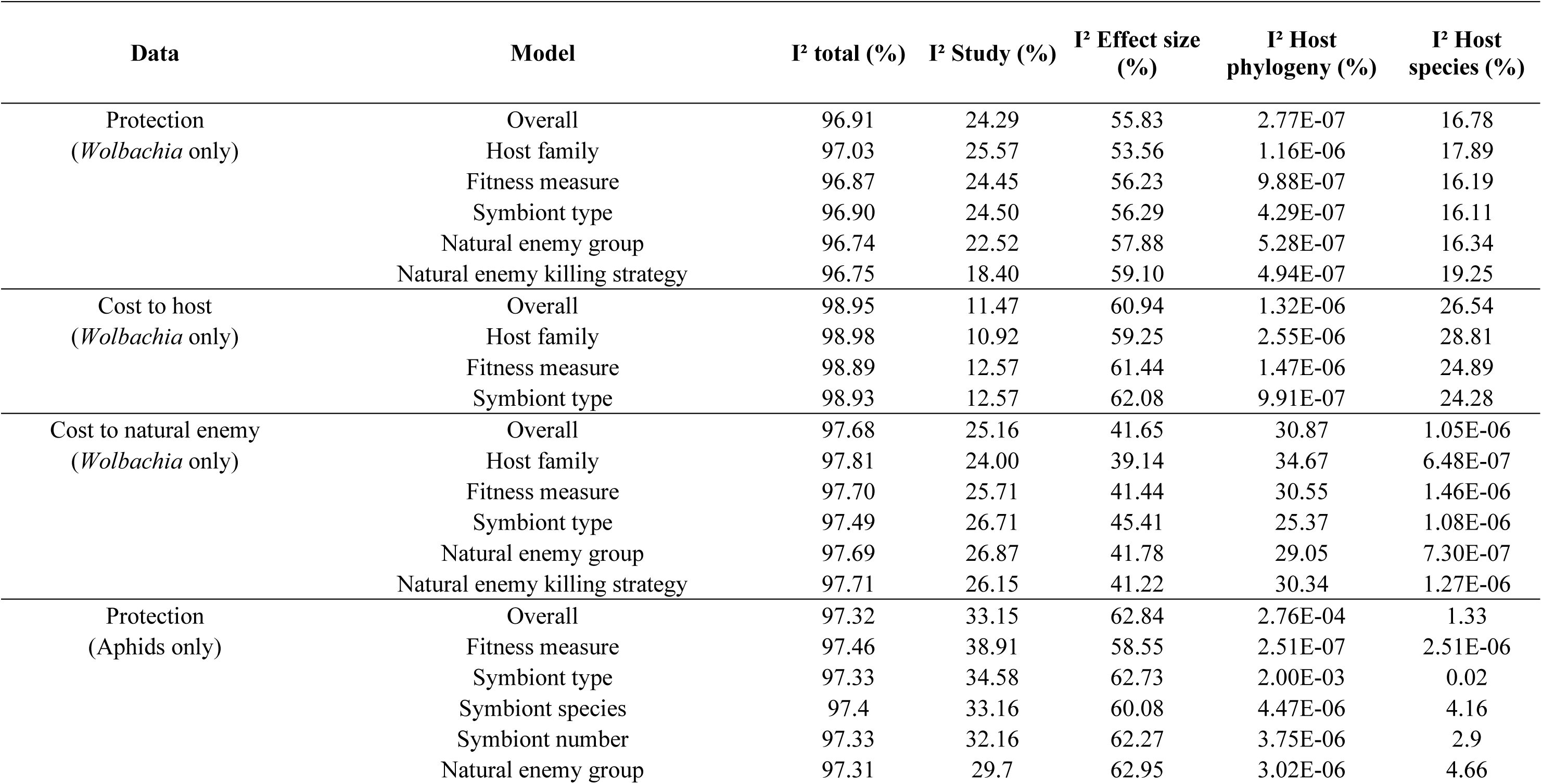
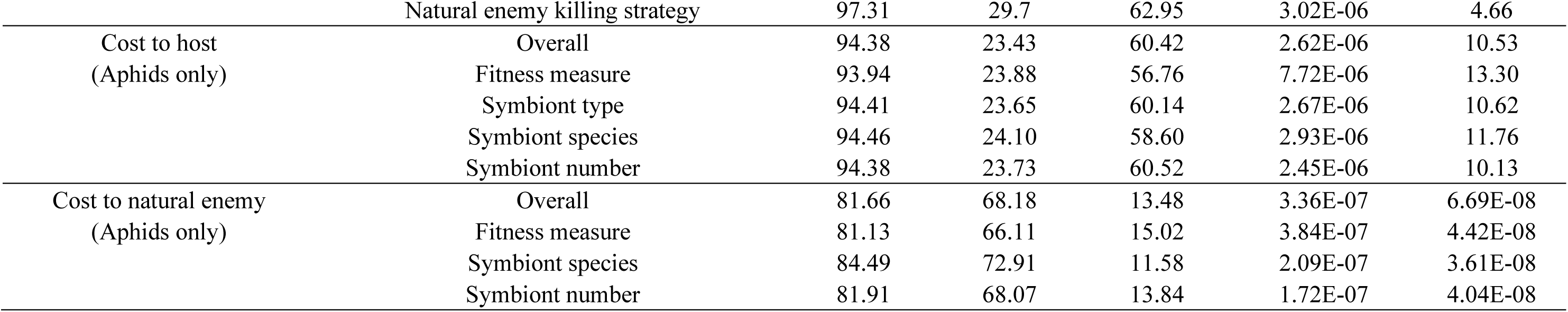
Subgroup analysis: results of the heterogeneity test (I^2^) for each meta-analytical model. For each model, we quantified the total amount of heterogeneity (I^2^) and the percentage of contribution of each random effect for the variance in the models: study, effect size, host phylogeny and host species.

**Table S14.**
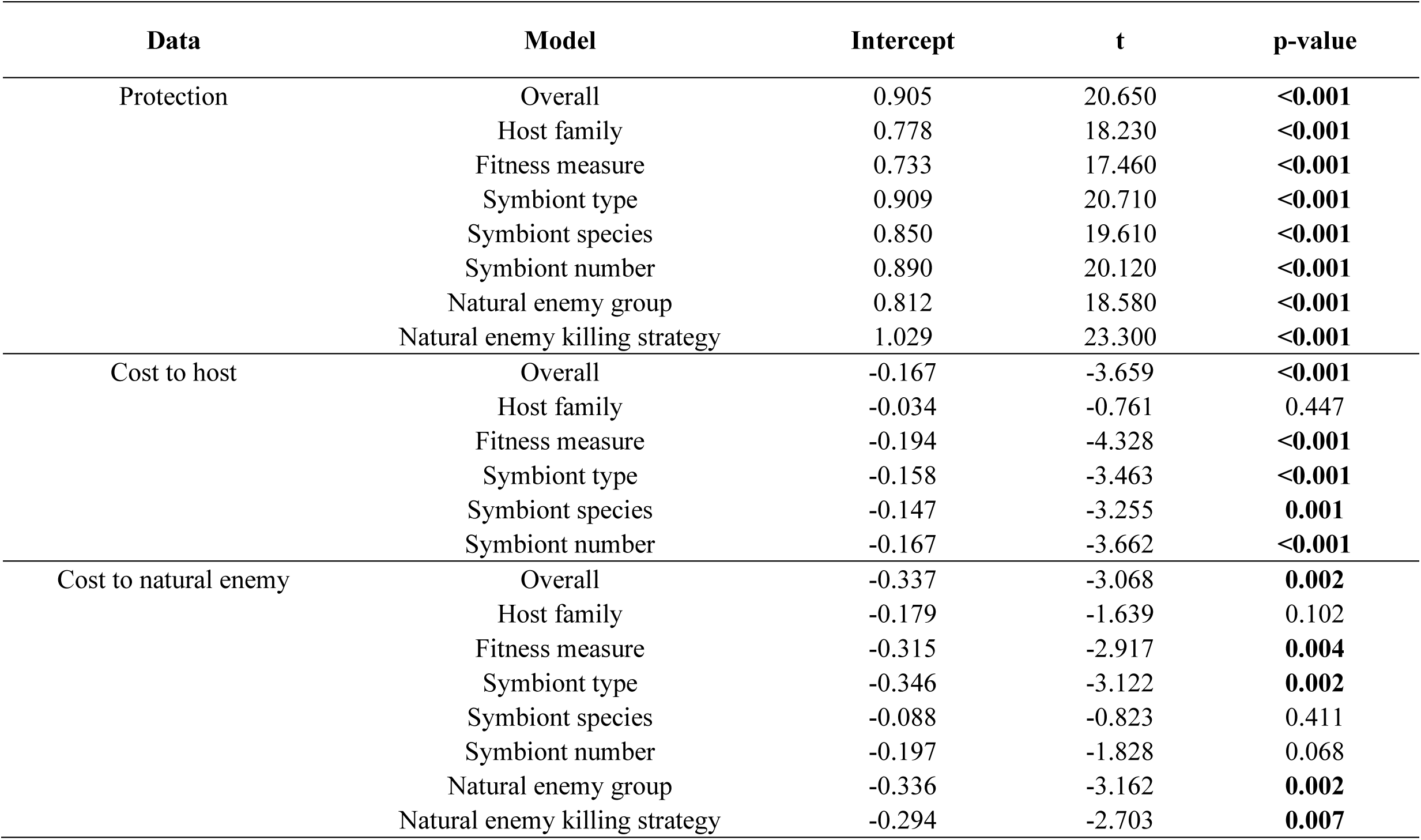
Main analysis: results of the Egger’s regression test for publication bias for each meta-analytical model. Models with p-values in bold represent the models with potential publication bias.

**Table S15.**
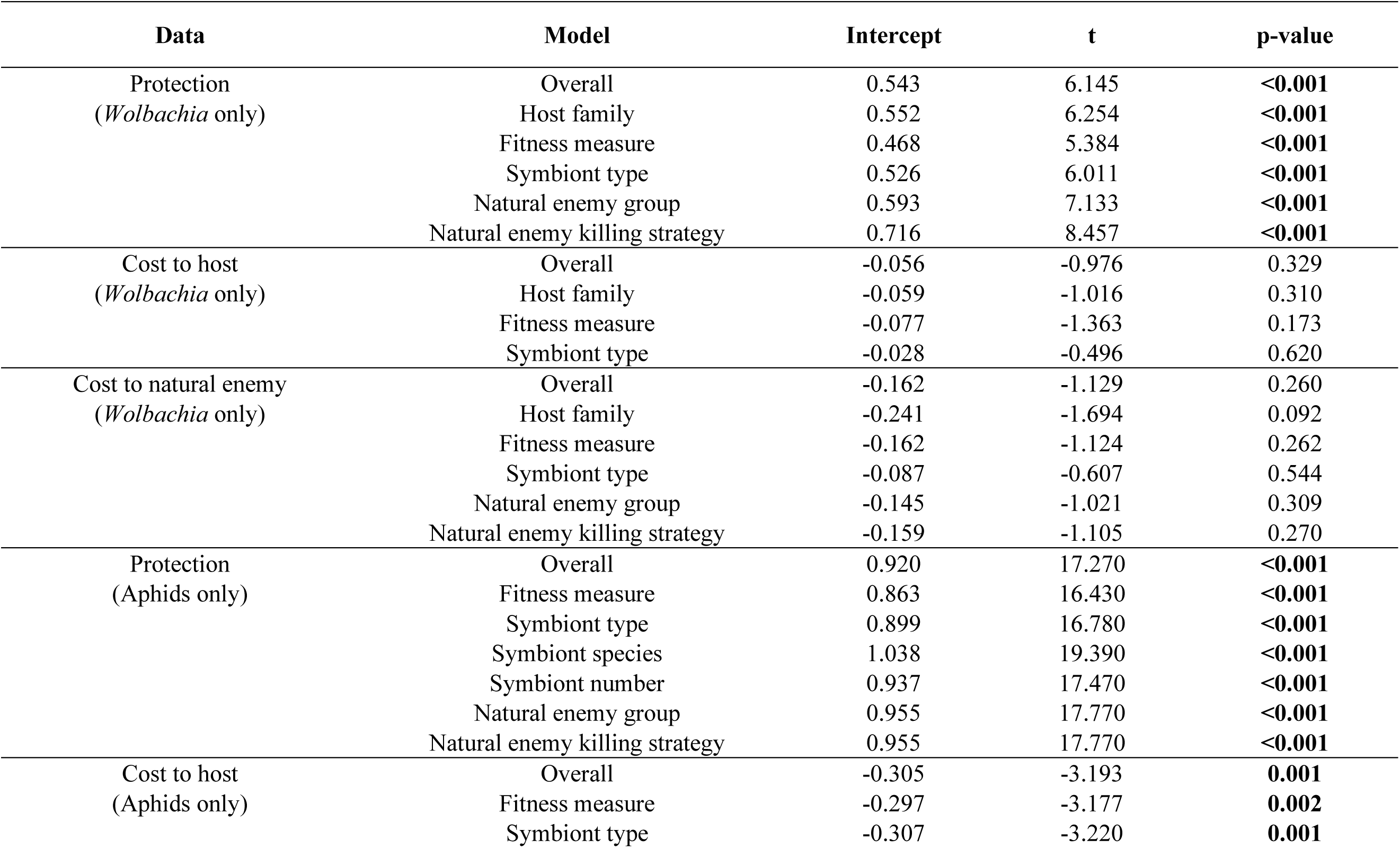
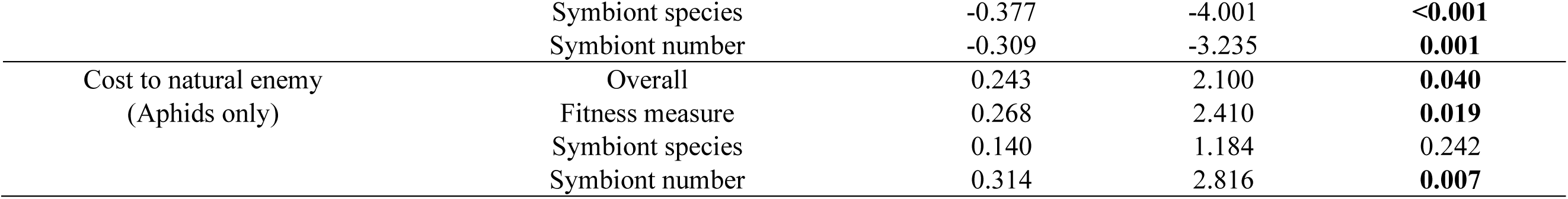
Subset analysis: results of the Egger’s regression test for publication bias for each meta-analytical model. Models with p-values in bold represent the models with potential publication bias.

**Table S16.**
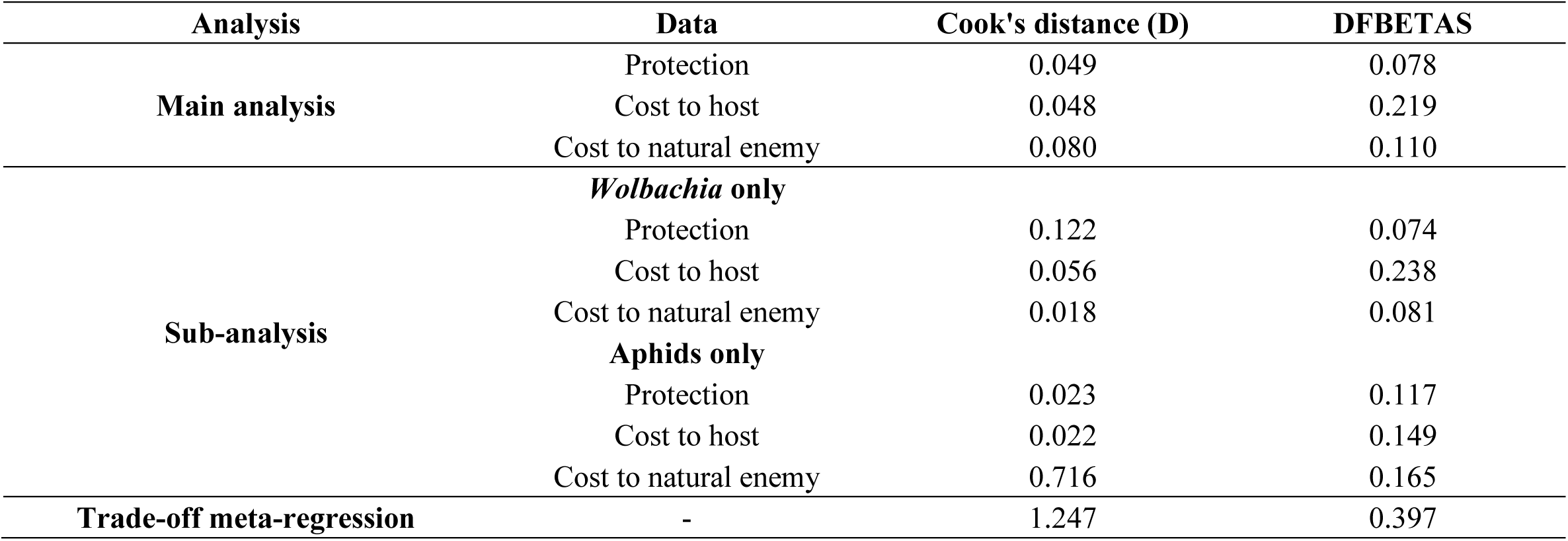
Results of the sensitivity analysis for the overall models, showing the maximum Cook’s distance and DFBETAs values for each model.

