## Supplementary material for "Defensive symbionts provide high protection against natural enemies at low cost to hosts: a meta-analysis": Suplementary material

**List of papers included in the meta-analysis with their respective ID in the raw data tables**

Ahmed MZ, Li SJ, Xue X, Yin XJ, Ren SX, Jiggins FM, Greeff JM, Qiu BL. 2015 The intracellular bacterium *Wolbachia* uses parasitoid wasps as phoretic vectors for efficient horizontal transmission. *PLoS Pathog.* **11**, e1004672. (doi:10.1371/journal.ppat.1004672) (P005)

Alexandrov ID, Alexandrova MV, Goryacheva II, Rochina NV, Shaikevich EV, Zakharov IA. 2007 Removing endosymbiotic *Wolbachia* specifically decreases lifespan of females and competitiveness in a laboratory strain of *Drosophila melanogaster*. *Russ. J. Genet.* **43**, 1147–1152. (doi: 10.1134/S1022795407100080) (P006)

Altinli M, Soms J, Ravallec M, Justy F, Bonneau M, Weill M, Gosselin-Grenet AS, Sicard M. 2019 Sharing cells with *Wolbachia*: the transovarian vertical transmission of *Culex pipiens* densovirus. *Environ. Microbiol.* **21**, 3284–3298. (doi:10.1111/1462-2920.14511) (P163)

Amuzu HE, Tsyganov K, Koh C, Herbert RI, Powell DR, McGraw EA. 2018 *Wolbachia* enhances insect-specific flavivirus infection in *Aedes aegypti* mosquitoes. *Ecol. Evol.* **8**, 5441–5454. (doi:10.1002/ece3.4066) (P069)

Andrews ES, Crain PR, Fu Y, Howe DK, Dobson SL. 2012 Reactive oxygen species production and *Brugia pahangi* survivorship in *Aedes polynesiensis* with artificial *Wolbachia* infection types. *PLoS Pathog.* **8,** e1003075. (doi:10.1371/journal.ppat.1003075) (P053)

Ant TH, Herd CS, Geoghegan V, Hoffmann AA, Sinkins SP. 2018 The *Wolbachia* strain wAu provides highly efficient virus transmission blocking in *Aedes aegypti*. *PLoS Pathog.* **14**, e1006815. (doi:10.1371/journal.ppat.1006815) (P022)

Arai H, Hirano T, Akizuki N, Abe A, Nakai M, Kunimi Y, Inoue MN. 2019 Multiple infection and reproductive manipulations of *Wolbachia* in *Homona magnanima* (Lepidoptera: Tortricidae). *Microb. Ecol.* **77**, 257–266. (doi:10.1007/s00248-018-1210-4) (P209)

Asiimwe P, Kelly SE, Hunter MS. 2014 Symbiont infection affects whitefly dynamics in the field. *Basic Appl. Ecol.* **15**, 507–515. (doi:10.1016/j.baae.2014.08.005) (P165)

Asplen MK, Bano N, Brady CM, Desneux N, Hopper KR, Malouines C, Oliver KM, White JA, Heimpel GE. 2014 Specialisation of bacterial endosymbionts that protect aphids from parasitoids. *Ecol. Entomol.* **39**, 736–739. (doi:10.1111/een.12153) (P055)

Axford JK, Ross PA, Yeap HL, Callahan AG, Hoffmann AA. 2016 Fitness of wAlbB *Wolbachia* infection in *Aedes aegypti*: parameter estimates in an outcrossed background and potential for population invasion. *Am. J. Trop. Med. Hyg.* **94**, 507–516. (doi:10.4269/ajtmh.15-0608) (P023)

Ayoubi A, Talebi AA, Fathipour Y, Mehrabadi M. 2018 Coinfection of the secondary symbionts, *Hamiltonella defensa* and *Arsenophonus sp*. contribute to the performance of the major aphid pest, *Aphis gossypii* (Hemiptera: Aphididae). *Insect Sci.* **27**, 86–98. (doi:10.1111/1744-7917.12603) (P025)

Ballinger MJ, Gawryluk RMR, Perlman SJ. 2019 Toxin and genome evolution in a *Drosophila* defensive symbiosis. *Genome Biol. Evol.* **11**, 253–262. (doi:10.1093/gbe/evy272) (P210)

Ballinger MJ, Perlman SJ. 2017 Generality of toxins in defensive symbiosis: Ribosome-inactivating proteins and defense against parasitic wasps in *Drosophila*. *PLoS Pathog.* **13**, e1006431. (doi:10.1371/journal.ppat.1006431) (P024)

Barribeau SM, Sok D, Gerardo NM. 2010 Aphid reproductive investment in response to mortality risks. *BMC Evol. Biol.* **10**, 251. (doi:10.1186/1471-2148-10-251) (P015)

Baton LA, Pacidônio EC, Gonçalves D da S, Moreira LA. 2013 wFlu: characterization and evaluation of a native *Wolbachia* from the mosquito *Aedes fluviatilis* as a potential vector control agent. *PLoS One* **8,** e0059619. (doi:10.1371/journal.pone.0059619) (P009)

Bian G, Joshi D, Dong Y, Lu P, Zhou G, Pan X, Xu Y, Dimopoulos G, Xi Z. 2013 *Wolbachia* invades *Anopheles stephensi* populations and induces refractoriness to *Plasmodium* infection. *Science.* **340**, 748–751. (doi:10.1126/science.1236192) (P028)

Bian G, Xu Y, Lu P, Xie Y, Xi Z. 2010 The endosymbiotic bacterium *Wolbachia* induces resistance to dengue virus in *Aedes aegypti*. *PLoS Pathog.* **6**, e1000833. (doi:10.1371/journal.ppat.1000833) (P187)

Bian G, Zhou G, Lu P, Xi Z. 2013 Replacing a native *Wolbachia* with a novel strain results in an increase in endosymbiont load and resistance to Dengue virus in a mosquito vector. *PLoS Negl. Trop. Dis.* **7,** e0002250. (doi:10.1371/journal.pntd.0002250) (P196)

Blagrove MSC, Arias-Goeta C, Di Genua C, Failloux AB, Sinkins SP. 2013 A *Wolbachia* wMel transinfection in *Aedes albopictus* is not detrimental to host fitness and inhibits Chikungunya virus. *PLoS Negl. Trop. Dis.* **7**, e0002152. (doi:10.1371/journal.pntd.0002152) (P027)

Blagrove MSC, Arias-Goeta C, Failloux AB, Sinkins SP. 2012 *Wolbachia* strain wMel induces cytoplasmic incompatibility and blocks dengue transmission in *Aedes albopictus*. *Proc. Natl.

*Acad. Sci. U.S.A.* **109**, 255–260. (doi:10.1073/pnas.1112021108) (P026)

Braquart-Varnier C, Altinli M, Pigeault R, Chevalier FD, Grève P, Bouchon D, Sicard M. 2015 The mutualistic side of *Wolbachia*-isopod interactions: *Wolbachia* mediated protection against pathogenic intracellular bacteria. *Front. Microbiol.* **6,** e201501388. (doi:10.3389/fmicb.2015.01388) (P168)

Brelsfoard CL, Dobson SL. 2011 *Wolbachia* effects on host fitness and the influence of male aging on cytoplasmic incompatibility in *Aedes polynesiensis* (Diptera: Culicidae). *J. Med. Entomol.* **48**, 1008–1015. (doi:10.1603/ME10202) (P032)

Brownlie JC, Cass BN, Riegler M, Witsenburg JJ, Iturbe-Ormaetxe I, McGraw EA, O'Neill SL. 2009 Evidence for metabolic provisioning by a common invertebrate endosymbiont, *Wolbachia pipientis*, during periods of nutritional stress. *PLoS Pathog.* **5,** e1000368. (doi:10.1371/journal.ppat.1000368) (P016)

Calvitti M, Moretti R, Lampazzi E, Bellini R, Dobson SL. 2010 Characterization of a new *Aedes albopictus* (Diptera: Culicidae)-*Wolbachia pipientis* (Rickettsiales: Rickettsiaceae) symbiotic association generated by artificial transfer of the wPip strain from *Culex pipiens* (Diptera: Culicidae). *J. Med. Entomol.* **47**, 179–187. (doi:10.1603/ME09140) (P039)

Cao LJ, Jiang W, Hoffmann AA. 2019 Life history effects linked to an advantage for wAu *Wolbachia* in *Drosophila*. *Insects* **10**, 126. (doi:10.3390/insects10050126) (P007)

Caragata EP, Rancès E, O'Neill SL, McGraw EA. 2014 Competition for amino acids between *Wolbachia* and the mosquito host, *Aedes aegypti*. *Microb. Ecol.* **67**, 205–218. (doi:10.1007/s00248-013-0339-4) (P197)

Cass BN, Himler AG, Bondy EC, Bergen JE, Fung SK, Kelly SE, Hunter MS. 2016 Conditional fitness benefits of the *Rickettsia* bacterial symbiont in an insect pest. *Oecologia* **180**, 169–179. (doi:10.1007/s00442-015-3436-x) (P038)

Castañeda LE, Sandrock C, Vorburger C. 2010 Variation and covariation of life history traits in aphids are related to infection with the facultative bacterial endosymbiont *Hamiltonella defensa*. *Biol. J. Linn. Soc.* **100**, 237–247. (doi:10.1111/j.1095-8312.2010.01416.x) (P001)

Cattel J, Nikolouli K, Andrieux T, Martinez J, Jiggins F, Charlat S, Vavre F, Lejon D, Gibert P, Moutone L. 2018 Back and forth *Wolbachia* transfers reveal efficient strains to control spotted wing *Drosophila* populations. *J. Appl. Ecol.* **55**, 2408–2418. (doi:10.1111/1365-2664.13101) (P040)

Cattel J, Martinez J, Jiggins F, Mouton L, Gibert P. 2016 *Wolbachia*-mediated protection against viruses in the invasive pest *Drosophila suzukii*. *Insect Mol. Biol.* **25**, 595–603. (doi:10.1111/imb.12245) (P204)

Cayetano L, Rothacher L, Simon JC, Vorburger C. 2015 Cheaper is not always worse: Strongly protective isolates of a defensive symbiont are less costly to the aphid host. *Proc. R. Soc. B Biol. Sci.* **282,** e20142333. (doi:10.1098/rspb.2014.2333) (P002)

Cayetano L, Vorburger C. 2013 Effects of heat shock on resistance to parasitoids and on life history traits in an aphid/endosymbiont system. *PLoS One* **8**, e0075966. (doi:10.1371/journal.pone.0075966) (P202)

Cayetano L, Vorburger C. 2013 Genotype-by-genotype specificity remains robust to average temperature variation in an aphid/endosymbiont/parasitoid system. *J. Evol. Biol.* **26**, 1603–1610. (doi:10.1111/jeb.12154) (P061)

Cayetano L, Vorburger C. 2015 Symbiont-conferred protection against Hymenopteran parasitoids in aphids: How general is it? *Ecol. Entomol.* **40**, 85–93. (doi:10.1111/een.12161) (P062)

Chen DQ, Montllor CB, Purcell AH. 2000 Fitness effects of two facultative endosymbiotic bacteria on the pea aphid, *Acyrthosiphon pisum*, and the blue alfalfa aphid, *A. kondoi*. *Entomol. Exp. Appl.* **95**, 315–323. (doi:10.1023/a:1004083324807) (P003)

Chiel E, Inbar M, Mozes-Daube N, White JA, Hunter MS, Zchori-Fein E. 2009 Assessments of fitness effects by the facultative symbiont *Rickettsia* in the sweetpotato whitefly (Hemiptera: Aleyrodidae). *Ann. Entomol. Soc. Am.* **102**, 413–418. (doi:10.1603/008.102.0309) (P042)

Chrostek E, Marialva MSP, Esteves SS, Weinert LA, Martinez J, Jiggins FM, Teixeira L. 2013 *Wolbachia* variants induce differential protection to viruses in *Drosophila melanogaster*: a phenotypic and phylogenomic analysis. *PLoS Genet.* **9**, e1003896. (doi:10.1371/journal.pgen.1003896) (P190)

Clarke HV., Foster SP, Oliphant L, Waters EW, Karley AJ. 2018 Co-occurrence of defensive traits in the potato aphid *Macrosiphum euphorbiae*. *Ecol. Entomol.* **43**, 538–542. (doi:10.1111/een.12522) (P063)

Clarke HV, Cullen D, Hubbard SF, Karley AJ. 2017 Susceptibility of *Macrosiphum euphorbiae* to the parasitoid *Aphidius ervi*: larval development depends on host aphid genotype. *Entomol. Exp. Appl.* **162**, 148–158. (doi: 10.1111/eea.12516) (P043)

Costopoulos K, Kovacs JL, Kamins A, Gerardo NM. 2014 Aphid facultative symbionts reduce survival of the predatory lady beetle *Hippodamia convergens*. *BMC Ecol.* **14**, 5. (doi:10.1186/1472-6785-14-5) (P183)

Dion E, Zélé F, Simon JC, Outreman Y. 2011 Rapid evolution of parasitoids when faced with the symbiont-mediated resistance of their hosts. *J. Evol. Biol.* **24**, 741–750. (doi:10.1111/j.1420-9101.2010.02207.x) (P072)

Dionysopoulou NK, Papanastasiou SA, Kyritsis GA, Papadopoulos NT. 2020 Effect of host fruit, temperature and *Wolbachia* infection on survival and development of *Ceratitis capitata* immature stages. *PLoS One.* **15,** e0229727. (doi:10.1371/journal.pone.0229727) (P224)

Dodson BL, Hughes GL, Paul O, Matacchiero AC, Kramer LD, Rasgon JL. 2014 *Wolbachia* Enhances West Nile Virus (WNV) Infection in the *Mosquito Culex tarsalis*. *PLoS Negl. Trop. Dis.* **8**, e0002965. (doi:10.1371/journal.pntd.0002965) (P064)

Donald KJ, Clarke H V., Mitchell C, Cornwell RM, Hubbard SF, Karley AJ. 2016 Protection of pea aphids associated with coinfecting bacterial symbionts persists during superparasitism by a braconid wasp. *Microb. Ecol.* **71**, 1–4. (doi:10.1007/s00248-015-0690-8) (P065)

Doremus MR, Oliver KM. 2017 Aphid heritable symbiont exploits defensive mutualism. *Appl. Environ. Microbiol.* **83**. (doi:10.1128/AEM.03276-16) (P044)

Doremus MR, Smith AH, Kim KL, Holder AJ, Russell JA, Oliver KM. 2018 Breakdown of a defensive symbiosis, but not endogenous defences, at elevated temperatures. *Mol. Ecol.* **27**, 2138–2151. (doi:10.1111/mec.14399) (P066)

Duplouy A, Couchoux C, Hanski I, Van Nouhuys S. 2015 *Wolbachia* infection in a natural parasitoid wasp population. *PLoS One* **10**, 1–16. (doi:10.1371/journal.pone.0134843) (P046)

Dutra HLC, Da Silva VL, Da Rocha Fernandes M, Logullo C, Maciel-De-Freitas R, Moreira LA. 2016 The influence of larval competition on brazilian *Wolbachia*-infected *Aedes aegypti* mosquitoes. *Parasit. Vectors.* **9**, 282. (doi:10.1186/s13071-016-1559-5) (P191)

Dutton TJ, Sinkins SP. 2005 Filarial susceptibility and effects of *Wolbachia* in *Aedes pseudoscutellaris* mosquitoes. *Med. Vet. Entomol.* **19**, 60–65. (doi:10.1111/j.0269-283X.2005.00557.x) (P057)

Dykstra HR, Weldon SR, Martinez AJ, White JA, Hopper KR, Heimpel GE, Asplen MK, Oliver KM. 2014 Factors limiting the spread of the protective symbiont *Hamiltonella defensa* in *Aphis craccivora* aphids. *Appl. Environ. Microbiol.* **80**, 5818–5827. (doi:10.1128/AEM.01775-14)

(P045)

Ebbert MA. 1991 The interaction phenotype in the *Drosophila willistoni*-Spiroplasma symbiosis. *Evolution.* **45**, 971. (doi:10.2307/2409703) (P130)

Farnesi LC, Belinato TA, Gesto JSM, Martins AJ, Bruno RV, Moreira LA. 2019 Embryonic development and egg viability of wMel-infected *Aedes aegypti*. *Parasit. Vectors.* **12**, 211. (doi:10.1186/s13071-019-3474-z) (P076)

Fleury F, Vavre F, Ris N, Fouillet P, Boulétreau M. 2000 Physiological cost induced by the maternally-transmitted endosymbiont *Wolbachia* in the *Drosophila* parasitoid *Leptopilina heterotoma*. *Parasitology* **121**, 493–500. (doi:10.1017/S0031182099006599) (P193)

Fraser JE, De Bruyne JT, Iturbe-Ormaetxe I, Stepnell J, Burns RL, Flores HA, O'Neill SL. 2017 Novel *Wolbachia*-transinfected *Aedes aegypti* mosquitoes possess diverse fitness and vector competence phenotypes. *PLoS Pathog.* **13**, e1006751. (doi:10.1371/journal.ppat.1006751) (P047)

Friberg U, Miller PM, Stewart AD, Rice WR. 2011 Mechanisms promoting the long-term persistence of a *Wolbachia* infection in a laboratory-adapted population of *Drosophila melanogaster*. *PLoS One* **6**, e16448. (doi:10.1371/journal.pone.0016448) (P048)

Fry AJ, Palmer MR, Rand DM. 2004 Variable fitness effects of *Wolbachia* infection in *Drosophila melanogaster*. *Heredity.* **93**, 379–389. (doi:10.1038/sj.hdy.6800514) (P029)

Fytrou A, Schofield PG, Kraaijeveld AR, Hubbard SF. 2006 *Wolbachia* infection suppresses both host defence and parasitoid counter-defence. *Proc. R. Soc. B Biol. Sci.* **273**, 791–796. (doi:10.1098/rspb.2005.3383) (P050)

Garcia G, Sylvestre G, Aguiar R, da Costa GB, Martins AJ, Lima JBP, Petersen MT, de Oliveira RL, Shadbolt MF, *et al.* 2019 Matching the genetics of released and local *Aedes aegypti* populations is critical to assure *Wolbachia* invasion. *PLoS Negl. Trop. Dis.* **13,** e0007023. (doi:10.1371/journal.pntd.0007023) (P186)

Ghosh S, Bouvaine S, Richardson SCW, Ghanim M, Maruthi MN. 2018 Fitness costs associated with infections of secondary endosymbionts in the cassava whitefly species *Bemisia tabaci*. *J. Pest Sci. (2004).* **91**, 17–28. (doi:10.1007/s10340-017-0910-8) (P049)

Gruntenko NE, Ilinsky YY, Adonyeva N V., Burdina E V., Bykov RA, Menshanov PN, Rauschenbach IY. 2017 Various *Wolbachia* genotypes differently influence host *Drosophila* dopamine metabolism and survival under heat stress conditions. *BMC Evol. Biol.* **17**, 252.

(doi:10.1186/s12862-017-1104-y) (P030)

Gruntenko NE, Karpova EK, Adonyeva N V., Andreenkova O V., Burdina E V., Ilinsky YY, Bykov RA, Menshanov PN, Rauschenbach IY. 2019 *Drosophila* female fertility and juvenile hormone metabolism depends on the type of *Wolbachia* infection. *J. Exp. Biol.* **222**, jeb195347. (doi:10.1242/jeb.195347) (P031)

Guidolin AS, Cônsoli FL. 2020 No fitness costs are induced by Spiroplasma infections of *Aphis citricidus* reared on two different host plants. *Brazilian J. Biol.* **80**, 311–318. (doi:10.1590/1519-6984.201210) (P211)

Guo Y, Hoffmann AA, Xu XQ, Zhang X, Huang HJ, Ju JF, Gong JT, Hong XY. 2018 *Wolbachia*-induced apoptosis associated with increased fecundity in *Laodelphax striatellus* (Hemiptera: Delphacidae). *Insect Mol. Biol.* **27**, 796–807. (doi:10.1111/imb.12518) (P194)

Guruprasad NM, Mouton L, Puttaraju HP. 2011 Effect of *Wolbachia* infection and temperature variations on the fecundity of the uzifly *Exorista sorbillans* (Diptera: Tachinidae). *Symbiosis* **54**, 151–158. (doi:10.1007/s13199-011-0138-y) (P195)

Hamm CA, Begun DJ, Vo A, Smith CCR, Saelao P, Shaver AO, Jaenike J, Turelli M. 2014 *Wolbachia* do not live by reproductive manipulation alone: infection polymorphism in *Drosophila suzukii* and *D. subpulchrella*. *Mol. Ecol.* **23**, 4871–4885. (doi:10.1111/mec.12901) (P077)

Hansen AK, Vorburger C, Moran NA. 2012 Genomic basis of endosymbiont-conferred protection against an insect parasitoid. *Genome Res.* **22**, 106–114. (doi:10.1101/gr.125351.111) (P176)

Harcombe W, Hoffmann AA. 2004 *Wolbachia* effects in *Drosophila melanogaster*: in search of fitness benefits. *J. Invertebr. Pathol.* **87**, 45–50. (doi:10.1016/j.jip.2004.07.003) (P021)

Haselkorn TS, Cockburn SN, Hamilton PT, Perlman SJ, Jaenike J. 2013 Infectious adaptation: potential host range of a defensive endosymbiont in *Drosophila*. *Evolution.* **67**, 934–945. (doi:10.1111/evo.12020) (P082)

Hedges LM, Brownlie JC, O'Neill SL, Johnson KN. 2008 *Wolbachia* and virus protection in insects. *Science.* **322**, 702. (doi:10.1126/science.1162418) (P070)

Hendry TA, Hunter MS, Baltrus DA. 2014 The facultative symbiont *Rickettsia* protects an invasive whitefly against entomopathogenic *Pseudomonas syringae* strains. *Appl. Environ. Microbiol.* **80**, 7161–7168. (doi:10.1128/AEM.02447-14) (P078)

Herren JK, Lemaitre B. 2011 Spiroplasma and host immunity: activation of humoral immune responses increases endosymbiont load and susceptibility to certain Gram-negative bacterial pathogens in *Drosophila melanogaster*. *Cell. Microbiol.* **13**, 1385–1396. (doi:10.1111/j.1462-5822.2011.01627.x) (P188)

Heyworth ER, Smee MR, Ferrari J. 2020 Aphid facultative symbionts aid recovery of their obligate symbiont and their host after heat stress. *Front. Ecol. Evol.* **8**, 56. (doi:10.3389/fevo.2020.00056) (P212)

Higashi CHV, Barton BT, Oliver KM. 2020 Warmer nights offer no respite for a defensive mutualism. *J. Anim. Ecol.* **89**, 1895–1905. (doi:10.1111/1365-2656.13238) (P225)

Himler AG, Adachi-Hagimori T, Bergen JE, Kozuch A, Kelly SE, Tabashnik BE, Chiel E, Duckworth VE, Dennehy TJ, Zchori-Fein E, Hunter MS. 2011 Rapid spread of a bacterial symbiont in an invasive whitefly is driven by fitness benefits and female bias. *Science.* **332**, 254–256. (doi:10.1126/science.1199410) (P079)

Hopper KR, Kuhn KL, Lanier K, Rhoades JH, Oliver KM, White JA, Asplen MK, Heimpel GE. 2018 The defensive aphid symbiont *Hamiltonella defensa* affects host quality differently for *Aphelinus glycinis* versus *Aphelinus atriplicis*. *Biol. Control* **116**, 3–9. (doi:10.1016/j.biocontrol.2017.05.008) (P080)

Hrček J, McLean AHC, Godfray HCJ. 2016 Symbionts modify interactions between insects and natural enemies in the field. *J. Anim. Ecol.* **85**, 1605–1612. (doi:10.1111/1365-2656.12586) (P083)

Hughes GL, Koga R, Xue P, Fukatsu T, Rasgon JL. 2011 *Wolbachia* infections are virulent and inhibit the human malaria parasite *Plasmodium falciparum* in *Anopheles gambiae. PLoS Pathog.* **7,** e.1002043. (doi:10.1371/journal.ppat.1002043) (P199)

Hughes GL, Vega-Rodriguez J, Xue P, Rasgon JL. 2012 *Wolbachia* strain wAlbB enhances infection by the rodent malaria parasite *Plasmodium berghei* in *Anopheles gambiae* mosquitoes. *Appl. Environ. Microbiol.* **78**, 1491–1495. (doi:10.1128/AEM.06751-11) (P198)

Hussain M, Lu G, Torres S, Edmonds JH, Kay BH, Khromykh AA, Asgari S. 2013 Effect of *Wolbachia* on replication of West Nile Virus in a mosquito cell line and adult mosquitoes. *J. Virol.* **87**, 851–858. (doi:10.1128/jvi.01837-12) (P084)

Islam MS, Dobson SL. 2006 *Wolbachia* effects on *Aedes albopictus* (Diptera: Culicidae) immature survivorship and development. *J. Med. Entomol.* **43**, 689–695. (doi:10.1603/0022-

2585(2006)43[689:WEOAAD]2.0.CO;2) (P085)

Izraeli Y, Lalzar M, Netanel N, Mozes-Daube N, Steinberg S, Chiel E, Zchori-Fein E. 2021 *Wolbachia* influence on the fitness of *Anagyrus vladimiri* (Hymenoptera: Encyrtidae), a bio-control agent of mealybugs. *Pest Manag. Sci.* **77**, 1023–1034. (doi:10.1002/ps.6117) (P213)

Jaenike J, Unckless R, Cockburn SN, Boelio LM, Perlman SJ. 2010 Adaptation via symbiosis: recent spread of a *Drosophila* defensive symbiont. *Science.* **329**, 212–215. (doi:10.1126/science.1188235) (P184)

Jamin AR, Vorburger C. 2019 Estimating costs of aphid resistance to parasitoids conferred by a protective strain of the bacterial endosymbiont *Regiella insecticola*. *Entomol. Exp. Appl.* **167**, 252–260. (doi:10.1111/eea.12749) (P074)

Jones JE, Hurst GDD. 2019 Symbiont-mediated protection varies with wasp genotype in the *Drosophila melanogaster-Spiroplasma* interaction. *bioRxiv* (doi:10.1101/691683) (P162)

Joubert DA, Walker T, Carrington LB, De Bruyne JT, Kien DH, Hoang Nle T, Chau NV, Iturbe-Ormaetxe I, Simmons CP, O'Neill SL. 2016 Establishment of a *Wolbachia* superinfection in *Aedes aegypti* mosquitoes as a potential approach for future resistance management. *PLoS Pathog.* **12**, e1005434. (doi:10.1371/journal.ppat.1005434) (P075)

Kambris Z, Blagborough AM, Pinto SB, Blagrove MSC, Godfray HCJ, Sinden RE, Sinkins SP. 2010 *Wolbachia* stimulates immune gene expression and inhibits plasmodium development in *Anopheles gambiae*. *PLoS Pathog.* **6,** e1001143. (doi:10.1371/journal.ppat.1001143) (P058)

Kambris Z, Cook PE, Phuc HK, Sinkins SP. 2009 Immune activation by life-shortening *Wolbachia* and reduced filarial competence in mosquitoes. *Science.* **326**, 134–136. (doi:10.1126/science.1177531) (P068)

Karr TL, Yang W, Feder ME. 1998 Overcoming cytoplasmic incompatibility in *Drosophila*. *Proc. R. Soc. B Biol. Sci.* **265**, 391–395. (doi:10.1098/rspb.1998.0307) (P192)

Koga R, Tsuchida T, Sakurai M, Fukatsu T. 2007 Selective elimination of aphid endosymbionts: effects of antibiotic dose and host genotype, and fitness consequences. *FEMS Microbiol. Ecol.* **60**, 229–239. (doi:10.1111/j.1574-6941.2007.00284.x) (P086)

Koop JL, Zeh DW, Bonilla MM, Zeh JA. 2009 Reproductive compensation favours male-killing *Wolbachia* in a live-bearing host. *Proc. R. Soc. B Biol. Sci.* **276**, 4021–4028. (doi:10.1098/rspb.2009.1230) (P208)

Kovacs JL, Wolf C, Voisin D, Wolf S. 2017 Evidence of indirect symbiont conferred protection against the predatory lady beetle *Harmonia axyridis* in the pea aphid. *BMC Ecol.* **17**, 1–11. (doi:10.1186/s12898-017-0136-x) (P182)

Laughton AM, Fan MH, Gerardo NM. 2014 The combined effects of bacterial symbionts and aging on life history traits in the pea aphid, *Acyrthosiphon pisum*. *Appl. Environ. Microbiol.* **80**, 470–477. (doi:10.1128/AEM.02657-13) (P081)

Le Clec'h W, Dittmer J, Raimond M, Bouchon D, Sicard M. 2017 Phenotypic shift in *Wolbachia* virulence towards its native host across serial horizontal passages. *Proc. R. Soc. B Biol. Sci.* **284**, 20171076. (doi:10.1098/rspb.2017.1076) (P200)

Leclair M, Polin S, Jousseaume T, Simon JC, Sugio A, Morlière S, Fukatsu T, Tsuchida T, Outreman Y. 2017 Consequences of coinfection with protective symbionts on the host phenotype and symbiont titres in the pea aphid system. *Insect Sci.* **24**, 798–808. (doi:10.1111/1744-7917.12380) (P087)

Lenhart PA, White JA. 2017 A defensive endosymbiont fails to protect aphids against the parasitoid community present in the field. *Ecol. Entomol.* **42**, 680–684. (doi:10.1111/een.12419) (P088)

Leonardo TE, Mondor EB. 2006 Symbiont modifies host life-history traits that affect gene flow. *Proc. R. Soc. B Biol. Sci.* **273**, 1079–1084. (doi:10.1098/rspb.2005.3408) (P181)

Leonardo TE. 2004 Removal of a specialization-associated symbiont does not affect aphid fitness. *Ecol. Lett.* **7**, 461–468. (doi:10.1111/j.1461-0248.2004.00602.x) (P180)

Li S, Liu D, Zhang R, Zhai Y, Huang X, Wang D, Shi X. 2018 Effects of a presumably protective endosymbiont on life-history characters and their plasticity for its host aphid on three plants. *Ecol. Evol.* **8**, 13004–13013. (doi:10.1002/ece3.4754) (P089)

Longdon B, Fabian DK, Hurst GDD, Jiggins FM. 2012 Male-killing *Wolbachia* do not protect *Drosophila bifasciata* against viral infection. *BMC Microbiol.* **12**, S8. (doi:10.1186/1471-2180-12-S1-S8) (P206)

Łukasik P, Dawid MA, Ferrari J, Godfray HCJ. 2013 The diversity and fitness effects of infection with facultative endosymbionts in the grain aphid, *Sitobion avenae*. *Oecologia* **173**, 985–996. (doi:10.1007/s00442-013-2660-5) (P041)

Lukasik P, Guo H, Van Asch M, Ferrari J, Godfray HCJ. 2013 Protection against a fungal pathogen conferred by the aphid facultative endosymbionts *Rickettsia* and *Spiroplasma* is expressed in multiple host genotypes and species and is not influenced by co-infection with another symbiont. *J. Evol. Biol.* **26**, 2654–2661. (doi:10.1111/jeb.12260) (P035)

Łukasik P, van Asch M, Guo H, Ferrari J, Godfray HC. 2013 Unrelated facultative endosymbionts protect aphids against a fungal pathogen. *Ecol. Lett.* **16**, 214-218. (doi: 10.1111/ele.12031) (P060)

Łukasik P, Guo H, van Asch M, Henry LM, Godfray HCJ, Ferrari J. 2015 Horizontal transfer of facultative endosymbionts is limited by host relatedness. *Evolution*. **69**, 2757–2766. (doi:10.1111/evo.12767) (P036)

Lukasik P, Hancock EL, Ferrari J, Godfray HCJ. 2011 Grain aphid clones vary in frost resistance, but this trait is not influenced by facultative endosymbionts. *Ecol. Entomol.* **36**, 790–793. (doi:10.1111/j.1365-2311.2011.01321.x) (P037)

Luo C, Gatti JL, Monticelli LS, Poirié M, Desneux N, Zhao H, Hu Z. 2020 An increased risk of parasitism mediated by the facultative symbiont *Regiella insecticola*. *J. Pest Sci. (2004).* **93**, 737–745. (doi:10.1007/s10340-019-01189-3) (P226)

Luo C, Luo K, Meng L, Wan B, Zhao H, Hu Z. 2017 Ecological impact of a secondary bacterial symbiont on the clones of *Sitobion avenae* (Fabricius) (Hemiptera: Aphididae). *Sci. Rep.* **7**, 40754. (doi:10.1038/srep40754) (P033)

Madhav M, Brown G, Morgan JAT, Asgari S, McGraw EA, James P. 2020 Transinfection of buffalo flies (*Haematobia irritans exigua*) with *Wolbachia* and effect on host biology. *Parasit Vectors* **13**, 296. (doi:10.1186/s13071-020-04161-8) (P215)

Mains JW, Brelsfoard CL, Crain PR, Huang Y, Dobson SL. 2013 Population impacts of *Wolbachia* on *Aedes albopictus*. *Ecol. Appl.* **23**, 493–501. (doi:10.1890/12-1097.1) (P034)

Martinez AJ, Doremus MR, Kraft LJ, Kim KL, Oliver KM. 2018 Multi-modal defences in aphids offer redundant protection and increased costs likely impeding a protective mutualism. *J. Anim. Ecol.* **87**, 464–477. (doi:10.1111/1365-2656.12675) (P013)

Martinez AJ, Kim KL, Harmon JP, Oliver KM. 2016 Specificity of multi-modal aphid defenses against two rival parasitoids. *PLoS One* **11,** e0154670. (doi:10.1371/journal.pone.0154670) (P014)

Martinez AJ, Weldon SR, Oliver KM. 2014 Effects of parasitism on aphid nutritional and protective symbioses. *Mol. Ecol.* **23**, 1594–1607. (doi:10.1111/mec.12550) (P056)

Martinez J, Cogni R, Cao C, Smith S, Illingworth CJR, Jiggins FM. 2016 Addicted? Reduced host resistance in populations with defensive symbionts. *Proc. R. Soc. B Biol. Sci.* **283**, 20160778. (doi:10.1098/rspb.2016.0778) (P059)

Martinez J, Duplouy A, Woolfit M, Vavre F, O'Neill SL, Varaldi J. 2012 Influence of the virus LbFV and of *Wolbachia* in a host-parasitoid interaction. *PLoS One* **7**, e0035081. (doi:10.1371/journal.pone.0035081) (P205)

Martinez J, Longdon B, Bauer S, Chan YS, Miller WJ, Bourtzis K, Teixeira L, Jiggins FM. 2014 Symbionts commonly provide broad spectrum resistance to viruses in insects: a comparative analysis of *Wolbachia* strains. *PLoS Pathog.* **10,** e1004369. (doi:10.1371/journal.ppat.1004369) (P054)

Martinez J, Ok S, Smith S, Snoeck K, Day JP, Jiggins FM. 2015 Should symbionts be nice or selfish? antiviral effects of *Wolbachia* are costly but reproductive parasitism is not. *PLoS Pathog.* **11,** e1005021. (doi:10.1371/journal.ppat.1005021) (P010)

Martinez J, Tolosana I, Ok S, Smith S, Snoeck K, Day JP, Jiggins FM. 2017 Symbiont strain is the main determinant of variation in *Wolbachia*-mediated protection against viruses across *Drosophila* species. *Mol. Ecol.* **26**, 4072–4084. (doi:10.1111/mec.14164) (P052)

Mateos M, Winter L, Winter C, Higareda-Alvear VM, Martinez-Romero E, Xie J. 2016 Independent origins of resistance or susceptibility of parasitic wasps to a defensive symbiont. *Ecol. Evol.* **6**, 2679–2687. (doi:10.1002/ece3.2085) (P020)

Mathé-Hubert H, Kaech H, Ganesanandamoorthy P, Vorburger C. 2019 Evolutionary costs and benefits of infection with diverse strains of *Spiroplasma* in pea aphids. *Evolution.* **73**, 1466–1481. (doi:10.1111/evo.13740) (P090)

McGraw EA, Merritt DJ, Droller JN, O'Neill SL. 2002 *Wolbachia* density and virulence attenuation after transfer into a novel host. *Proc. Natl. Acad. Sci. U. S. A.* **99**, 2918–2923. (doi:10.1073/pnas.052466499) (P094)

McLean AHC, Ferrari J, Godfray HCJ. 2018 Do facultative symbionts affect fitness of pea aphids in the sexual generation? *Entomol. Exp. Appl.* **166**, 32–40. (doi:10.1111/eea.12641) (P171)

McLean AHC, Godfray HCJ. 2015 Evidence for specificity in symbiont conferred protection against parasitoids. *Proc. R. Soc. B Biol. Sci.* **282,** e20150977. (doi:10.1098/rspb.2015.0977) (P091)

McLean AHC, Godfray HCJ. 2017 The outcome of competition between two parasitoid species is influenced by a facultative symbiont of their aphid host. *Funct. Ecol.* **31**, 927–933. (doi:10.1111/1365-2435.12781) (P092)

McLean AHC, Hrček J, Parker BJ, Mathé-Hubert H, Kaech H, Paine C, Godfray HCJ. 2020 Multiple phenotypes conferred by a single insect symbiont are independent: Independent phenotypes of aphid symbiont. *Proc. R. Soc. B Biol. Sci.* **287,** e20200562. (doi:10.1098/rspb.2020.0562) (P135)

McLean AHC, Van Asch M, Ferrari J, Godfray HCJ. 2011 Effects of bacterial secondary symbionts on host plant use in pea aphids. *Proc. R. Soc. B Biol. Sci.* **278**, 760–766. (doi:10.1098/rspb.2010.1654) (P170)

McMeniman CJ, O'Neill SL. 2010 A virulent *Wolbachia* infection decreases the viability of the dengue vector *Aedes aegypti* during periods of embryonic quiescence. *PLoS Negl. Trop. Dis.* **4,** e0000748. (doi:10.1371/journal.pntd.0000748) (P172)

Meany MK, Conner WR, Richter S V., Bailey JA, Turelli M, Cooper BS. 2019 Loss of cytoplasmic incompatibility and minimal fecundity effects explain relatively low *Wolbachia* frequencies in *Drosophila mauritiana*. *Evolution.* **73**, 1278–1295. (doi:10.1111/evo.13745) (P161)

Miller WJ, Ehrman L, Schneider D. 2010 Infectious speciation revisited: impact of symbiont-depletion on female fitness and mating behavior of *Drosophila paulistorum*. *PLoS Pathog.* **6**, e1001214. (doi:10.1371/journal.ppat.1001214) (P017)

Montenegro H, Petherwick AS, Hurst GDD, Klaczko LB. 2006 Fitness effects of *Wolbachia* and *Spiroplasma* in *Drosophila melanogaster*. *Genetica.* **127**, 207–215. (doi:10.1007/s10709-005-3766-4) (P018)

Moreira LA, Iturbe-Ormaetxe I, Jeffery JA, Lu G, Pyke AT, Hedges LM, Rocha BC, Hall-Mendelin S, Day A, Riegler M, Hugo LE, Johnson KN, Kay BH, McGraw EA, van den Hurk AF, Ryan PA, O'Neill SL. 2009 A *Wolbachia* symbiont in *Aedes aegypti* limits infection with Dengue, Chikungunya, and Plasmodium. *Cell.* **139**, 1268–1278. (doi:10.1016/j.cell.2009.11.042) (P093)

Moretti R, Yen PS, Houé V, Lampazzi E, Desiderio A, Failloux AB, Calvitti M. 2018 Combining *Wolbachia*-induced sterility and virus protection to fight *Aedes albopictus*-borne viruses. *PLoS Negl. Trop. Dis.* **12**, e0006626. (doi:10.1371/journal.pntd.0006626) (P095)

Mousson L, Martin E, Zouache K, Madec Y, Mavingui P, Failloux AB. 2010 *Wolbachia* modulates Chikungunya replication in *Aedes albopictus*. *Mol. Ecol.* **19**, 1953–1964. (doi:10.1111/j.1365-294X.2010.04606.x) (P096)

Mousson L, Zouache K, Arias-Goeta C, Raquin V, Mavingui P, Failloux AB. 2012 The native *Wolbachia* symbionts limit transmission of Dengue virus in *Aedes albopictus*. *PLoS Negl. Trop. Dis.* **6,** e0001989. (doi:10.1371/journal.pntd.0001989) (P164)

Nakayama S, Parratt SR, Hutchence KJ, Lewis Z, Price TAR, Hurst GDD. 2015 Can maternally inherited endosymbionts adapt to a novel host? Direct costs of *Spiroplasma* infection, but not vertical transmission efficiency, evolve rapidly after horizontal transfer into *D. melanogaster*. *Heredity.* **114**, 539–543. (doi:10.1038/hdy.2014.112) (P097)

Niepoth N, Ellers J, Henry LM. 2018 Symbiont interactions with non-native hosts limit the formation of new symbioses. *BMC Evol. Biol.* **18**, 27. (doi:10.1186/s12862-018-1143-z) (P098)

Noriyuki S, Suzuki-Ohno Y, Takakura KI. 2016 Variation of clutch size and trophic egg proportion in a ladybird with and without male-killing bacterial infection. *Evol. Ecol.* **30**, 1081–1095. (doi:10.1007/s10682-016-9861-4) (P099)

Nyabuga FN, Outreman Y, Simon JC, Heckel DG, Weisser WW. 2010 Effects of pea aphid secondary endosymbionts on aphid resistance and development of the aphid parasitoid *Aphidius ervi*: A correlative study. *Entomol. Exp. Appl.* **136**, 243–253. (doi:10.1111/j.1570-7458.2010.01021.x) (P166)

O'Shea KL, Singh ND. 2015 Tetracycline-exposed *Drosophila melanogaster* males produce fewer offspring but a relative excess of sons. *Ecol. Evol.* **5**, 3130–3139. (doi:10.1002/ece3.1535) (P105)

Okayama K, Katsuki M, Sumida Y, Okada K. 2016 Costs and benefits of symbiosis between a bean beetle and *Wolbachia*. *Anim. Behav.* **119**, 19–26. (doi:10.1016/j.anbehav.2016.07.004) (P167)

Oliver KM, Campos J, Moran NA, Hunter MS. 2008 Population dynamics of defensive symbionts in aphids. *Proc. R. Soc. B Biol. Sci.* **275**, 293–299. (doi:10.1098/rspb.2007.1192) (P102)

Oliver KM, Moran NA, Hunter MS. 2005 Variation in resistance to parasitism in aphids is due to symbionts not host genotype. *Proc. Natl. Acad. Sci. U. S. A.* **102**, 12795–12800. (doi:10.1073/pnas.0506131102) (P101)

Oliver KM, Moran NA, Hunter MS. 2006 Costs and benefits of a superinfection of facultative symbionts in aphids. *Proc. R. Soc. B Biol. Sci.* **273**, 1273–1280. (doi:10.1098/rspb.2005.3436) (P103)

Oliver KM, Noge K, Huang EM, Campos JM, Becerra JX, Hunter MS. 2012 Parasitic wasp responses to symbiont-based defense in aphids. *BMC Biol.* **10**, 11. (doi:10.1186/1741-7007-10-11) (P104)

Oliver KM, Russell JA, Morant NA, Hunter MS. 2003 Facultative bacterial symbionts in aphids confer resistance to parasitic wasps. *Proc. Natl. Acad. Sci. U.S.A.* **100**, 1803–1807. (doi:10.1073/pnas.0335320100) (P100)

Olsen K, Reynolds KT, Hoffmann AA. 2001 A field cage test of the effects of the endosymbiont *Wolbachia* on *Drosophila melanogaster*. *Heredity.* **86**, 731–737. (doi:10.1046/j.1365-2540.2001.00892.x) (P019)

Osborne SE, Leong YS, O'Neill SL, Johnson KN. 2009 Variation in antiviral protection mediated by different *Wolbachia* strains in *Drosophila simulans*. *PLoS Pathog.* **5**, e1000656. (doi:10.1371/journal.ppat.1000656) (P173)

Panteleev DY, Goryacheva II, Andrianov B V., Reznik NL, Lazebny OE, Kulikov AM. 2007 The endosymbiotic bacterium *Wolbachia* enhances the nonspecific resistance to insect pathogens and alters behavior of *Drosophila melanogaster*. *Russ. J. Genet.* **43**, 1066–1069. (P107)

Parker BJ, Spragg CJ, Altincicek B, Gerardo NM. 2013 Symbiont-mediated protection against fungal pathogens in pea aphids: A role for pathogen specificity. *Appl. Environ. Microbiol.* **79**, 2455–2458. (doi:10.1128/AEM.03193-12) (P106)

Poinsot D, Mercot H. 1997 *Wolbachia* infection in *Drosophila simulans*: does the female host bear a physiological cost? *Evolution.* **51**, 180. (doi:10.2307/2410971) (P108)

Polin S, Le Gallic JF, Simon JC, Tsuchida T, Outreman Y. 2015 Conditional reduction of predation risk associated with a facultative symbiont in an insect. *PLoS One* **10,** e0143728. (doi:10.1371/journal.pone.0143728) (P174)

Pons I, Renoz F, Hance T. 2019 Fitness costs of the cultivable symbiont *Serratia symbiotica* and its phenotypic consequences to aphids in presence of environmental stressors. *Evol. Ecol.* **33**, 825–838. (doi:10.1007/s10682-019-10012-5) (P216)

Pons I, Renoz F, Noël C, Hance T. 2019 New insights into the nature of symbiotic associations in aphids: infection process, biological effects, and transmission mode of cultivable *Serratia symbiotica* bacteria. *Appl. Environ. Microbiol.* **85,** e02445-18. (doi:10.1128/AEM.02445-18) (P116)

Puttaraju HP, Prakash BM. 2005 Effects of *Wolbachia*-targeted tetracycline on a host-parasitoid-symbiont interaction. *Eur. J. Entomol.* **102**, 669–674. (doi:10.14411/eje.2005.095) (P115)

Qian L, Jia F, Jingxuan S, Manqun W, Julian C. 2018 Effect of the secondary symbiont *Hamiltonella defensa* on fitness and relative abundance of *Buchnera aphidicola* of wheat aphid, *Sitobion* *miscanthi*. *Front. Microbiol.* **9**, 582. (doi:10.3389/fmicb.2018.00582) (P114)

Rasgon JL. 2012 *Wolbachia* induces male-specific mortality in the mosquito *Culex pipiens* (lin strain). *PLoS One* **7**, e30381. (doi:10.1371/journal.pone.0030381) (P119)

Reyes ML, Laughton AM, Parker BJ, Wichmann H, Fan M, Sok D, Hrček J, Acevedo T, Gerardo NM. 2019 The influence of symbiotic bacteria on reproductive strategies and wing polyphenism in pea aphids responding to stress. *J. Anim. Ecol.* **88**, 601–611. (doi:10.1111/1365-2656.12942) (P118)

Reynolds KT, Thomson LJ, Hoffmann AA. 2003 The effects of host age, host nuclear background and temperature on phenotypic effects of the virulent *Wolbachia* strain popcorn in *Drosophila melanogaster*. *Genetics.* **164**, 1027–1034. (doi:10.1093/genetics/164.3.1027) (P175)

Richardson KM, Griffin PC, Lee SF, Ross PA, Endersby-Harshman NM, Schiffer M, Hoffmann AA. 2019 A *Wolbachia* infection from *Drosophila* that causes cytoplasmic incompatibility despite low prevalence and densities in males. *Heredity.* **122**, 428–440. (doi:10.1038/s41437-018-0133-7) (P117)

Riegler M, Charlat S, Stauffer C, Merçot H. 2004 *Wolbachia* transfer from rhagoletis cerasi to *Drosophila simulans*: investigating the outcomes of host-symbiont coevolution. *Appl. Environ. Microbiol.* **70**, 273–279. (doi:10.1128/AEM.70.1.273-279.2004) (P120)

Ross PA, Endersby NM, Yeap HL, Hoffmann AA. 2014 Larval competition extends developmental time and decreases adult size of wMelPop *Wolbachia*-infected *Aedes aegypti*. *Am. J. Trop. Med. Hyg.* **91**, 198–205. (doi:10.4269/ajtmh.13-0576) (P179)

Rottschaefer SM, Lazzaro BP. 2012 No effect of *Wolbachia* on resistance to intracellular infection by pathogenic bacteria in *Drosophila melanogaster*. *PLoS One*. **7**, e40500. (doi:10.1371/journal.pone.0040500) (P121)

McLean AHC, Hrček J, Parker BJ, Godfray HCJ. 2017 Cascading effects of herbivore protective symbionts on hyperparasitoids. *Ecol. Entomol.* **42**:601-609. (doi:10.1111/een.12424) (P122)

Ruang-Areerate T, Kittayapong P. 2006 *Wolbachia* transinfection in *Aedes aegypti*: A potential gene driver of dengue vectors. *Proc. Natl. Acad. Sci. U.S.A.* **103**, 12534–12539. (doi:10.1073/pnas.0508879103) (P123)

Russell JA, Moran NA. 2005 Horizontal transfer of bacterial symbionts: Heritability and fitness effects in a novel aphid host. *Appl. Environ. Microbiol.* **71**, 7987–7994. (doi:10.1128/AEM.71.12.7987-7994.2005) (P177)

Russell JA, Moran NA. 2006 Costs and benefits of symbiont infection in aphids: Variation among symbionts and across temperatures. *Proc. R. Soc. B Biol. Sci.* **273**, 603–610. (doi:10.1098/rspb.2005.3348) (P127)

Sakurai M, Koga R, Tsuchida T, Meng XY, Fukatsu T. 2005 *Rickettsia* symbiont in the pea aphid *Acyrthosiphon pisum*: Novel cellular tropism, effect on host fitness, and interaction with the essential symbiont *Buchnera*. *Appl. Environ. Microbiol.* **71**, 4069–4075. (doi:10.1128/AEM.71.7.4069-4075.2005) (P126)

Sarakatsanou A, Diamantidis AD, Papanastasiou SA, Bourtzis K, Papadopoulos NT. 2011 Effects of *Wolbachia* on fitness of the Mediterranean fruit fly (Diptera: Tephritidae). *J. Appl. Entomol.* **135**, 554–563. (doi:10.1111/j.1439-0418.2011.01610.x) (P129)

Scarborough CL, Ferrari J, Godfray HCJ. 2005 Ecology: Aphid protected from pathogen by endosymbiont. *Science.* **310**, 1781. (doi:10.1126/science.1120180) (P124)

Schmid M, Sieber R, Zimmermann YS, Vorburger C. 2012 Development, specificity and sublethal effects of symbiont-conferred resistance to parasitoids in aphids. *Funct. Ecol.* **26**, 207–215. (doi:10.1111/j.1365-2435.2011.01904.x) (P125)

Schneider DI *et al.* 2019 Spatio-temporal distribution of *Spiroplasma* infections in the tsetse fly (*Glossina fuscipes fuscipes*) in northern Uganda. *PLoS Negl. Trop. Dis.* **13**, e0007340. (doi:10.1371/journal.pntd.0007340) (P159)

Schneider DI, Riegler M, Arthofer W, Merçot H, Stauffer C, Miller WJ. 2013 Uncovering *Wolbachia* diversity upon artificial host transfer. *PLoS One* **8,** e0082402. (doi:10.1371/journal.pone.0082402) (P178)

Semiatizki A, Weiss B, Bagim S, Rohkin-Shalom S, Kaltenpoth M, Chiel E. 2020 Effects, interactions, and localization of *Rickettsia* and *Wolbachia* in the house fly parasitoid, *Spalangia endius*. *Microb. Ecol.* **80**, 718–728. (doi:10.1007/s00248-020-01520-x) (P217)

Serga SV., Maistrenko OM, Matiytsiv NP, Vaiserman AM, Kozeretska IA. 2021 Effects of *Wolbachia* infection on fitness-related traits in *Drosophila melanogaster*. *Symbiosis* **83**, 163–172. (doi:10.1007/s13199-020-00743-3) (P218)

Shan HW, Zhang CR, Yan TT, Tang HQ, Wang XW, Liu SS, Liu YQ. 2016 Temporal changes of symbiont density and host fitness after rifampicin treatment in a whitefly of the *Bemisia tabaci* species complex. *Insect Sci.* **23**, 200–214. (doi:10.1111/1744-7917.12276) (P128)

Simon JC, Boutin S, Tsuchida T, Koga R, Gallic JF, Frantz A, Outreman Y, Fukatsu T. 2011 Facultative symbiont infections affect aphid reproduction. *PLoS One*. **6**, e21831. (doi:10.1371/journal.pone.0021831) (P132)

Simon JC, Sakurai M, Bonhomme J, Tsuchida T, Koga R, Fukatsu T. 2007 Elimination of a specialised facultative symbiont does not affect the reproductive mode of its aphid host. *Ecol. Entomol.* **32**, 296–301. (doi:10.1111/j.1365-2311.2007.00868.x) (P131)

Singh R, Linksvayer TA. 2020 *Wolbachia*-infected ant colonies have increased reproductive investment and an accelerated life cycle. *J. Exp. Biol.* **223**, jeb220079. (doi:10.1242/jeb.220079) (P158)

Skelton E, Rancès E, Frentiu FD, Kusmintarsih ES, Iturbe-Ormaetxe I, Caragata EP, Woolfit M, O'Neill SL. 2016 A native *Wolbachia* endosymbiont does not limit dengue virus infection in the mosquito *Aedes notoscriptus* (Diptera: Culicidae). *J. Med. Entomol.* **53**, 401–408. (doi:10.1093/jme/tjv235) (P133)

Sontowski R, Gerth M, Richter S, Gruppe A, Schlegel M, van Dam NM, Bleidorn C. 2020 Infection patterns and fitness effects of *Rickettsia* and *Sodalis* symbionts in the green lacewing *Chrysoperla carnea*. *Insects* **11**, 1–17. (doi:10.3390/insects11120867) (P157)

Su Q Oliver KM, Pan H, Jiao X, Liu B, Xie W, Wang S, Wu Q, Xu B, White JA, Zhou X, Zhang Y. 2013 Facultative symbiont *Hamiltonella* confers benefits to *Bemisia tabaci* (Hemiptera: Aleyrodidae), an invasive agricultural pest worldwide. *Environ. Entomol.* **42**, 1265–1271. (doi:10.1603/EN13182) (P134)

Sumida Y, Katsuki M, Okada K, Okayama K, Lewis Z. 2017 *Wolbachia* induces costs to life-history and reproductive traits in the moth *Ephestia kuehniella*. *J. Stored Prod. Res.* **71**, 93–98. (doi:10.1016/j.jspr.2017.02.003) (P169)

Suzuki J, Uda A, Watanabe K, Shimizu T, Watarai M. 2016 Symbiosis with *Francisella tularensis* provides resistance to pathogens in the silkworm. *Sci. Rep.* **6**, 31476. (doi:10.1038/srep31476) (P185)

Teixeira L, Ferreira Á, Ashburner M. 2008 The bacterial symbiont *Wolbachia* induces resistance to RNA viral infections in *Drosophila melanogaster*. *PLoS Biol.* **6**, 2753–2763. (doi:10.1371/journal.pbio.1000002) (P189)

Tsuchida T, Koga R, Fujiwara A, Fukatsu T. 2014 Phenotypic effect of Candidatus *Rickettsiella viridis*, a facultative symbiont of the pea aphid (*Acyrthosiphon pisum*), and its interaction with a coexisting symbiont. *Appl. Environ. Microbiol.* **80**, 525–533. (doi:10.1128/AEM.03049-13) (P138)

Turley AP, Zalucki MP, O'Neill SL, McGraw EA. 2013 Transinfected *Wolbachia* have minimal effects on male reproductive success in *Aedes aegypti*. *Parasit. Vectors* **6**, 36. (doi:10.1186/1756-3305-6-36) (P136)

Unckless RL, Jaenike J. 2012 Maintenance of a male-killing *Wolbachia* in *Drosophila innubila* by male-killing dependent and male-killing independent mechanisms. *Evolution.* **66**, 678–689. (doi:10.1111/j.1558-5646.2011.01485.x) (P142)

Vala F, Breeuwer JAJ, Sabelis MW. 2003 Sorting out the effects of *Wolbachia*, genotype and inbreeding on life-history traits of a spider mite. *Exp. Appl. Acarol.* **29**, 253–264. (doi:10.1023/A:1025810414956) (P139)

van den Hurk AF, Hall-Mendelin S, Pyke AT, Frentiu FD, McElroy K, Day A, Higgs S, O'Neill SL. 2012 Impact of *Wolbachia* on infection with Chikungunya and Yellow Fever viruses in the mosquito vector *Aedes aegypti. PLoS Negl. Trop. Dis.* **6,** e0001892. (doi:10.1371/journal.pntd.0001892) (P145)

van Nouhuys S, Kohonen M, Duplouy A. 2016 *Wolbachia* increases the susceptibility of a parasitoid wasp to hyperparasitism. *J. Exp. Biol.* **219**, 2984–2990. (doi:10.1242/jeb.140699) (P067)

Vasquez CJ, Stouthamer R, Jeong G, Morse JG. 2011 Discovery of a CI-inducing *Wolbachia* and its associated fitness costs in the biological control agent *Aphytis melinus* DeBach (Hymenoptera: Aphelinidae). *Biol. Control* **58**, 192–198. (doi:10.1016/j.biocontrol.2011.06.006) (P137)

Von Burg S, Ferrari J, Müller CB, Vorburger C. 2008 Genetic variation and covariation of susceptibility to parasitoids in the aphid *Myzus persicae*: No evidence for trade-offs. *Proc. R. Soc. B Biol. Sci.* **275**, 1089–1094. (doi:10.1098/rspb.2008.0018) (P143)

Vorburger C, Ganesanandamoorthy P, Kwiatkowski M. 2013 Comparing constitutive and induced costs of symbiont-conferred resistance to parasitoids in aphids. *Ecol. Evol.* **3**, 706–713. (doi:10.1002/ece3.491) (P207)

Vorburger C, Gehrer L, Rodriguez P. 2010 A strain of the bacterial symbiont *Regiella insecticola* protects aphids against parasitoids. *Biol. Lett.* **6**, 109–111. (doi:10.1098/rsbl.2009.0642) (P141)

Vorburger C, Gouskov A. 2011 Only helpful when required: A longevity cost of harbouring defensive symbionts. *J. Evol. Biol.* **24**, 1611–1617. (doi:10.1111/j.1420-9101.2011.02292.x) (P140)

Vorburger C, Rouchet R. 2016 Are aphid parasitoids locally adapted to the prevalence of defensive symbionts in their hosts? *BMC Evol. Biol.* **16**, 1–11. (doi:10.1186/s12862-016-0811-0) (P154)

Vorburger C, Sandrock C, Gouskov A, Castañeda LE, Ferrari J. 2009 Genotypic variation and the role of defensive endosymbionts in an all-parthenogenetic host-parasitoid interaction. *Evolution.* **63**, 1439–1450. (doi:10.1111/j.1558-5646.2009.00660.x) (P151)

Walker T, Johnson PH, Moreira LA, Iturbe-Ormaetxe I, Frentiu FD, McMeniman CJ, Leong YS, Dong Y, Axford J, Kriesner P, Lloyd AL, Ritchie SA, O'Neill SL, Hoffmann AA. 2011 The wMel *Wolbachia* strain blocks dengue and invades caged *Aedes aegypti* populations. *Nature* **476**, 450–455. (doi:10.1038/nature10355) (P153)

Wang D, Shi X, Dai P, Liu D, Dai X, Shang Z, Ge Z, Meng X. 2016 Comparison of fitness traits and their plasticity on multiple plants for *Sitobion avenae* infected and cured of a secondary endosymbiont. *Sci. Rep.* **6**, 23177. (doi:10.1038/srep23177) (P150)

Weldon SR, Russell JA, Oliver KM. 2020 More is not always better: Coinfections with defensive symbionts generate highly variable outcomes. *Appl. Environ. Microbiol.* **86**, 1–14. (doi:10.1128/AEM.02537-19) (P214)

White JA, Kelly SE, Cockburn SN, Perlman SJ, Hunter MS. 2011 Endosymbiont costs and benefits in a parasitoid infected with both *Wolbachia* and *Cardinium*. *Heredity.* **106**, 585–591. (doi:10.1038/hdy.2010.89) (P149)

Wong ZS, Hedges LM, Brownlie JC, Johnson KN. 2011 *Wolbachia*-mediated antibacterial protection and immune gene regulation in *Drosophila*. *PLoS One* **6,** e0025430. (doi:10.1371/journal.pone.0025430) (P071)

Wu LH, Hoffmann AA, Thomson LJ. 2016 *Trichogramma* parasitoids for control of Lepidopteran borers in Taiwan: Species, life-history traits and *Wolbachia* infections. *J. Appl. Entomol.* **140**, 353–363. (doi:10.1111/jen.12263) (P160)

Xie J, Butler S, Sanchez G, Mateos M. 2014 Male killing *Spiroplasma* protects *Drosophila melanogaster* against two parasitoid wasps. *Heredity.* **112**, 399–408. (doi:10.1038/hdy.2013.118) (P012)

Xie J, Tiner B, Vilchez I, Mateos M. 2011 Effect of the *Drosophila* endosymbiont *Spiroplasma* on parasitoid wasp development and on the reproductive fitness of wasp-attacked fly survivors. *Evol. Ecol.* **25**, 1065–1079. (doi:10.1007/s10682-010-9453-7) (P004)

Xie J, Vilchez I, Mateos M. 2010 Spiroplasma bacteria enhance survival of *Drosophila hydei* attacked by the parasitic wasp *Leptopilina heterotoma*. *PLoS One* **5**, e12149. (doi:10.1371/journal.pone.0012149) (P011)

Xie J, Winter C, Winter L, Mateos M. 2015 Rapid spread of the defensive endosymbiont *Spiroplasma* in *Drosophila hydei* under high parasitoid wasp pressure. *FEMS Microbiol. Ecol.* **91**, 1-11. (doi:10.1093/femsec/iu017) (P203)

Xu P, Liu Y, Graham RI, Wilson K, Wu K. 2014 Densovirus is a mutualistic symbiont of a global crop pest (*Helicoverpa armigera*) and protects against a Baculovirus and Bt biopesticide. *PLoS Pathog.* **10**, e1004490. (doi:10.1371/journal.ppat.1004490) (P146)

Xue X, Li SJ, Ahmed MZ, De Barro PJ, Ren SX, Qiu BL. 2012 Inactivation of *Wolbachia* reveals its biological roles in whitefly host. *PLoS One.* **7**, e48148. (doi:10.1371/journal.pone.0048148) (P144)

Yadav S, Frazer J, Banga A, Pruitt K, Harsh S, Jaenike J, Eleftherianos I. 2018 Endosymbiont-based immunity in *Drosophila melanogaster* against parasitic nematode infection. *PLoS One*. **13**, e0192183. (doi:10.1371/journal.pone.0192183) (P201)

Yang K, Xie K, Zhu YX, Huo SM, Hoffmann A, Hong XY. 2020 *Wolbachia* dominate *Spiroplasma* in the co-infected spider mite *Tetranychus truncatus*. *Insect Mol. Biol.* **29**, 19–37. (doi:10.1111/imb.12607) (P223)

Ye YH, Carrasco AM, Dong Y, Sgrò CM, McGraw EA. 2016 The effect of temperature on *Wolbachia*-mediated dengue virus blocking in *Aedes aegypti*. *Am. J. Trop. Med. Hyg.* **94**, 812–819. (doi:10.4269/ajtmh.15-0801) (P147)

Ye YH, Woolfit M, Rancès E, O'Neill SL, McGraw EA. 2013 *Wolbachia*-associated bacterial protection in the mosquito *Aedes aegypti*. *PLoS Negl. Trop. Dis.* **7**, e2362. (doi:10.1371/journal.pntd.0002362) (P148)

Yeap HL, Mee P, Walker T, Weeks AR, O'Neill SL, Johnson P, Ritchie SA, Richardson KM, Doig C, Endersby NM, Hoffmann AA. 2011 Dynamics of the "Popcorn" *Wolbachia* infection in outbred *Aedes aegypti* informs prospects for mosquito vector control. *Genetics.* **187**, 583–595. (doi:10.1534/genetics.110.122390) (P152)

Yoshida K, Sanada-Morimura S, Huang SH, Tokuda M. 2019 Influences of two coexisting endosymbionts, CI-inducing *Wolbachia* and male-killing *Spiroplasma*, on the performance of their host *Laodelphax striatellus* (Hemiptera: Delphacidae). *Ecol. Evol.* **9**, 8214–8224.

(doi:10.1002/ece3.5392) (P222)

Zélé F, Altıntas M, Santos I, Cakmak I, Magalhães S. 2020 Population-specific effect of *Wolbachia* on the cost of fungal infection in spider mites. *Ecol. Evol.* **10**, 3868–3880. (doi:10.1002/ece3.6015) (P155)

Zélé F, Denoyelle J, Duron O, Rivero A. 2018 Can *Wolbachia* modulate the fecundity costs of *Plasmodium* in mosquitoes? *Parasitology* **145,** 775–782. (doi:10.1017/S0031182017001330) (P008)

Zélé F, Nicot A, Berthomieu A, Weill M, Duron O, Rivero A. 2014 *Wolbachia* increases susceptibility to *Plasmodium* infection in a natural system. *Proc. R. Soc. B Biol. Sci.* **281,** 20132837. (doi:10.1098/rspb.2013.2837) (P073)

Zélé F, Nicot A, Duron O, Rivero A. 2012 Infection with *Wolbachia* protects mosquitoes against *Plasmodium*-induced mortality in a natural system. *J. Evol. Biol.* **25**, 1243–1252. (doi:10.1111/j.1420-9101.2012.02519.x) (P051)

Zélé F, Santos JL, Godinho DP, Magalhães S. 2018 *Wolbachia* both aids and hampers the performance of spider mites on different host plants. *FEMS Microbiol. Ecol.* **94**. (doi:10.1093/femsec/fiy187) (P156)

Zhang CR, Shan HW, Xiao N, Zhang F Di, Wang XW, Liu YQ, Liu SS. 2015 Differential temporal changes of primary and secondary bacterial symbionts and whitefly host fitness following antibiotic treatments. *Sci. Rep.* **5**, 15898. (doi:10.1038/srep15898) (P112)

Zhang XF, Zhao DX, Hong XY. 2012 *Cardinium*-the leading factor of cytoplasmic incompatibility in the planthopper *Sogatella furcifera* doubly infected with *Wolbachia* and *Cardinium. Environ. Entomol.* **41**, 833–840. (doi:10.1603/EN12078) (P111)

Zhang YK, Yang K, Zhu YX, Hong XY. 2018 Symbiont-conferred reproduction and fitness benefits can favour their host occurrence. *Ecol. Evol.* **8**, 1626–1633. (doi:10.1002/ece3.3784) (P113)

Zhao DX, Zhang XF, Chen DS, Zhang YK, Hong XY. 2013 *Wolbachia*-host interactions: host mating patterns affect *Wolbachia* density dynamics. *PLoS One.* **8,** e0066373. (doi:10.1371/journal.pone.0066373) (P109)

Zhao DX, Zhang XF, Hong XY. 2013 Host-symbiont interactions in spider mite *Tetranychus truncates* doubly infected with *Wolbachia* and *Cardinium*. *Environ. Entomol.* **42**, 445–452. (doi:10.1603/EN12354) (P110)

Zhu YX, Song YL, Hoffmann AA, Jin PY, Huo SM, Hong XY. 2019 A change in the bacterial community of spider mites decreases fecundity on multiple host plants. *MicrobiologyOpen* **8**, e00743. (doi:10.1002/mbo3.743) (P221)

Zhu YX, Song ZR, Huo SM, Yang K, Hong XY. 2020 Variation in the microbiome of the spider mite *Tetranychus truncatus* with sex, instar and endosymbiont infection. *FEMS Microbiol. Ecol.* **96**, fiaa004. (doi:10.1093/femsec/fiaa004) (P220)

Zhu YX, Song ZR, Song YL, Hong XY. 2020 Double infection of *Wolbachia* and *Spiroplasma* alters induced plant defense and spider mite fecundity. *Pest Manag. Sci.* **76**, 3273–3281. (doi:10.1002/ps.5886) (P219)

Ali M, Sajjad A, Shakeel Q, Farooqi MA, Aqueel MA, Tariq K, Ullah MI, Iqbal A, Jamal A, Saeed MF, et al. 2022 Influence of bacterial secondary symbionts in *Sitobion avenae* on its survival fitness against entomopathogenic fungi, *Beauveria bassiana* and *Metarhizium brunneum*. *Insects* **13**, 1037. (doi:10.3390/insects13111037) (P227)

Ayoubi A, Talebi AA, Fathipour Y, Hoffmann AA, Mehrabadi M. 2025 Symbiont-mediated insect host defense against parasitism: insights from the endosymbiont, *Hamiltonella defensa* and the insect host, *Myzus persicae*. *Pest Manag. Sci*. **81**, 4886–4893. (doi:10.1002/ps.8844) (P228)

Bilgo E, Mancini MV, Gnambani JE, Dokpomiwa HAT, Murdochy S, Lovett B, St Leger R, Sinkins SP, Diabate A. 2024 *Wolbachia* confers protection against the entomopathogenic fungus *Metarhizium pingshaense* in African *Aedes aegypti*. *Environ. Microbiol. Rep.* **16**, e13316. (doi:10.1111/1758-2229.13316) (P229)

Bruner-Montero G, Jiggins FM. 2023 *Wolbachia* protects *Drosophila melanogaster* against two naturally occurring and virulent viral pathogens. *Sci. Rep.* **13**, 8518. (doi:10.1038/s41598-023-35726-z) (P230)

Clavé C, Sugio A, Morlière S, Pincebourde S, Simon JC, Foray V. 2022 Physiological costs of facultative endosymbionts in aphids assessed from energy metabolism. *Funct. Ecol.* **36**, 2580–2592. (doi:10.1111/1365-2435.14157) (P231)

Deconninck G, Larges J, Henri H, et al. 2024 *Wolbachia* improves the performance of an invasive fly after a diet shift. *J. Pest Sci.* **97**, 2087–2099. (doi:10.1007/s10340-023-01739-w) (P232)

Dera K-sM, Barro DT, Kaboré BA, Gstöttenmayer F, Dieng MM, Pagabeleguem S, Weiss BL, Fiorenza G, Piccinno R, Malacrida AR, Aksoy S, de Beer CJ, Mach RL, Vreysen MJB, Abd-Alla AMM.

2025 *Spiroplasma* infection in colonized *Glossina fuscipes fuscipes*: impact on mass rearing and the sterile insect technique. *Insect Sci.* **32**, 1761–1776. (doi:10.1111/1744-7917.70078) (P233)

Gu X, Berran M, Prithiv Sivaji Dorai A, Yang Q, Stelmach M, Ross PA, Gill A, Ansermin E, Yeatman E, Umina PA, Hoffmann AA. 2025 Transinfections of the endosymbiont *Rickettsiella viridis* in different *Myzus persicae* (Hemiptera: Aphididae) clones show consistent deleterious effects and stable transmission. *J. Econ. Entomol.* **118**, 1544–1552. (doi:10.1093/jee/toaf114) (P234)

Higashi CHV, Nichols WL, Chevignon G, Patel V, Allison SE, Kim KL, Strand MR, Oliver KM. 2023 An aphid symbiont confers protection against a specialized RNA virus, another increases vulnerability to the same pathogen. *Mol. Ecol.* **32**, 936–950. (doi:10.1111/mec.16801) (P235)

Higashi CHV, Patel V, Kamalaker B, Inaganti R, Bressan A, Russell JA, et al. 2024 Another tool in the toolbox: Aphid-specific *Wolbachia* protect against fungal pathogens. *Environ. Microbiol.* **26**, e70005. (doi:10.1111/1462-2920.70005) (P236)

Kaech H, Jud S, Vorburger C. 2022 Similar cost of *Hamiltonella defensa* in experimental and natural aphid-endosymbiont associations. *Ecol. Evol.* **12**, e8551. (doi:10.1002/ece3.8551) (P237)

Kalyanaraman D, Riemsloh KMZ, Kürschner LK, Gadau J, Lammers M. 2023 Substantial fitness costs in terms of parasitization rates, offspring production and sex ratio of *Wolbachia* infection in *Nasonia vitripennis* without modified host preference. *Ecol. Entomol.* **48**, 765–774. (doi:10.1111/een.13271) (P238)

Kiefer JST, Schmidt G, Krüsemer R, Kaltenpoth M, Engl T. 2022 *Wolbachia* causes cytoplasmic incompatibility but not male-killing in a grain pest beetle. *Mol. Ecol.* **31**, 6570–6587. (doi:10.1111/mec.16717) (P239)

Heidari Latibari M, Moravvej G, Rakhshani E, Karimi J, Arias-Penna DC, Butcher BA. 2023 *Arsenophonus*: a double-edged sword of aphid defense against parasitoids. *Insects* **14**, 763. (doi:10.3390/insects14090763) (P240)

de Paula Sena LC, Moreira MM, de Paula Bouzada Dias L, Guedes JJM, Yotoko KSC. 2026 Reduced fecundity associated with *Wolbachia* infection in a Neotropical drosophilid. *Ecol. Entomol.* 51, 475–488. (doi:10.1111/een.70051) (P241)

Strunov A, Lerch S, Blanckenhorn WU, Miller WJ, Kapun M. 2022 Complex effects of environment and *Wolbachia* infections on the life history of *Drosophila melanogaster* hosts. *J. Evol. Biol.* **35**, 788–802. (doi:10.1111/jeb.14016) (P242)

Ueda M, Arai H, Masaike K, Nakai M, Inoue MN. 2023 Distinct effects of three *Wolbachia* strains on fitness and immune traits in *Homona magnanima*. *Heredity* **130**, 22–29. (doi:10.1038/s41437-022-00574-6) (P243)

Wang ZW, Zhao J, Li GY, Hu D, Wang ZG, Ye C, Wang JJ. 2024 The endosymbiont *Serratia symbiotica* improves aphid fitness by disrupting the predation strategy of ladybeetle larvae. *Insect Sci.* 31, 1555–1568. (doi:10.1111/1744-7917.13315) (P244)

Zhu DH, Liu LT. 2024 Cytoplasmic incompatibility and female fecundity associated with *Wolbachia* infection in a cricket species. *Ecol. Entomol.* **49**, 67–76. (doi:10.1111/een.13282) (P245)

Katlav A, Cook JM, Riegler M. 2022 Common endosymbionts affect host fitness and sex allocation via egg size provisioning. *Proc. R. Soc. B.* **289**, 20212582. (doi:10.1098/rspb.2021.2582) (P246)

### Supplementary figures

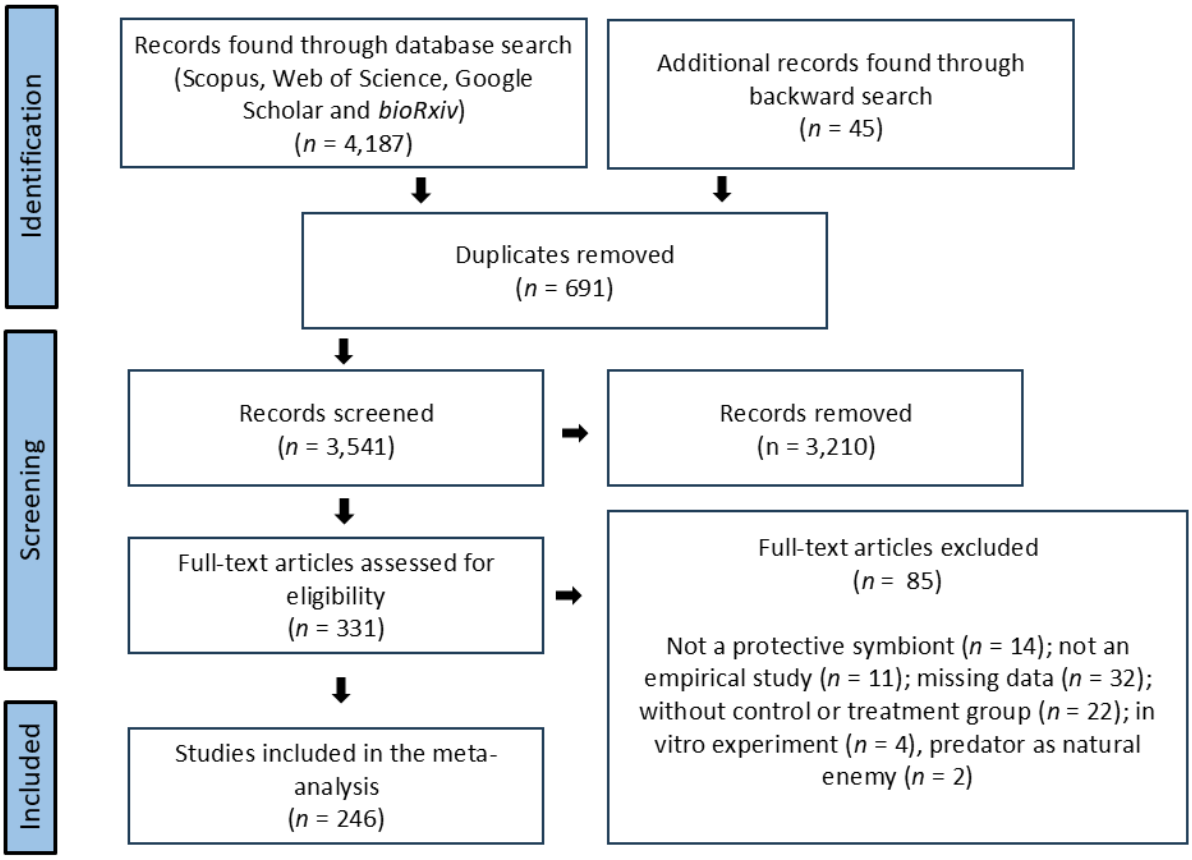

**Figure S1.** Full screening process of inclusion and exclusion of studies following the PRISMA protocol.

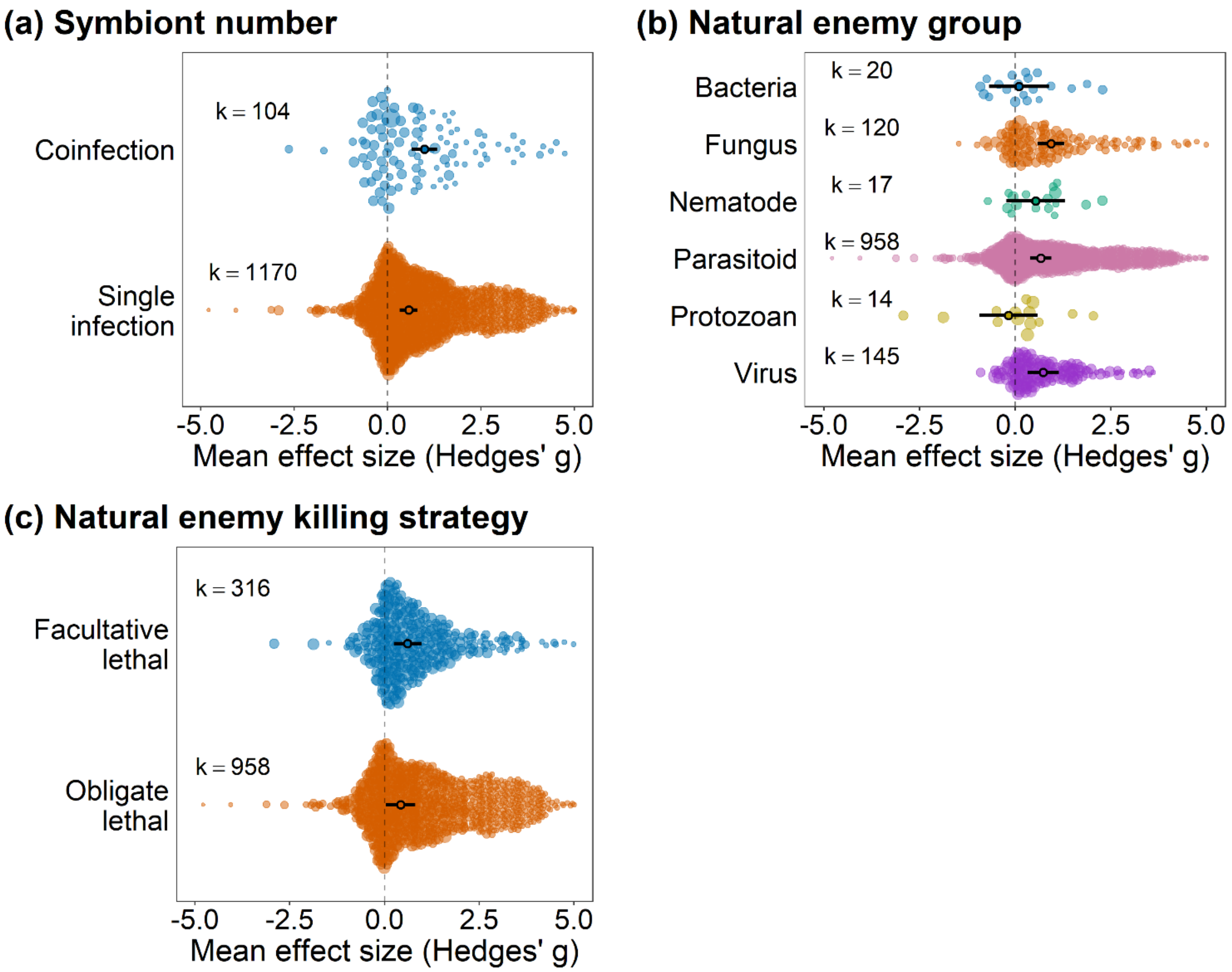

**Figure S2.** Effect of defensive symbionts on the fitness of hosts that are under attack of natural enemies. (a) model including symbiont number as moderator, (b) model including natural enemy group as moderator, and (c) model including natural enemy killing strategy as moderator. Points are the weighted mean effect sizes ± 95% confidence intervals. Each colored circle represents an effect size, scaled according to precision (inverse of the sampling error variance). 'k' is the number of effect sizes.

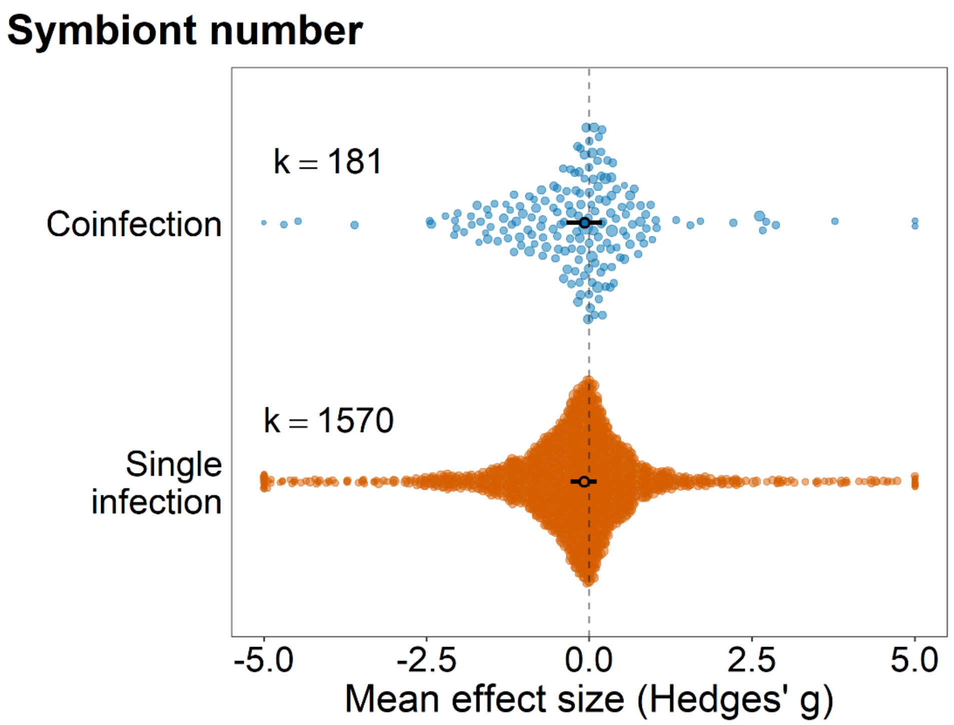

**Figure S3.** Effect of defensive symbionts on the fitness of hosts that are in the absence of natural enemies, including symbiont number as moderator. Points are the weighted mean effect sizes ± 95% confidence intervals. Each colored circle represents an effect size, scaled according to precision (inverse of the sampling error variance). 'k' is the number of effect sizes. Sixteen points fell below −5 and 13 points fell above +5, which expanded the x-axis range and compressed the visual spacing of the remaining points. For visualization purposes only, these points are shown at ±5. All data points were included in the statistical analyses.

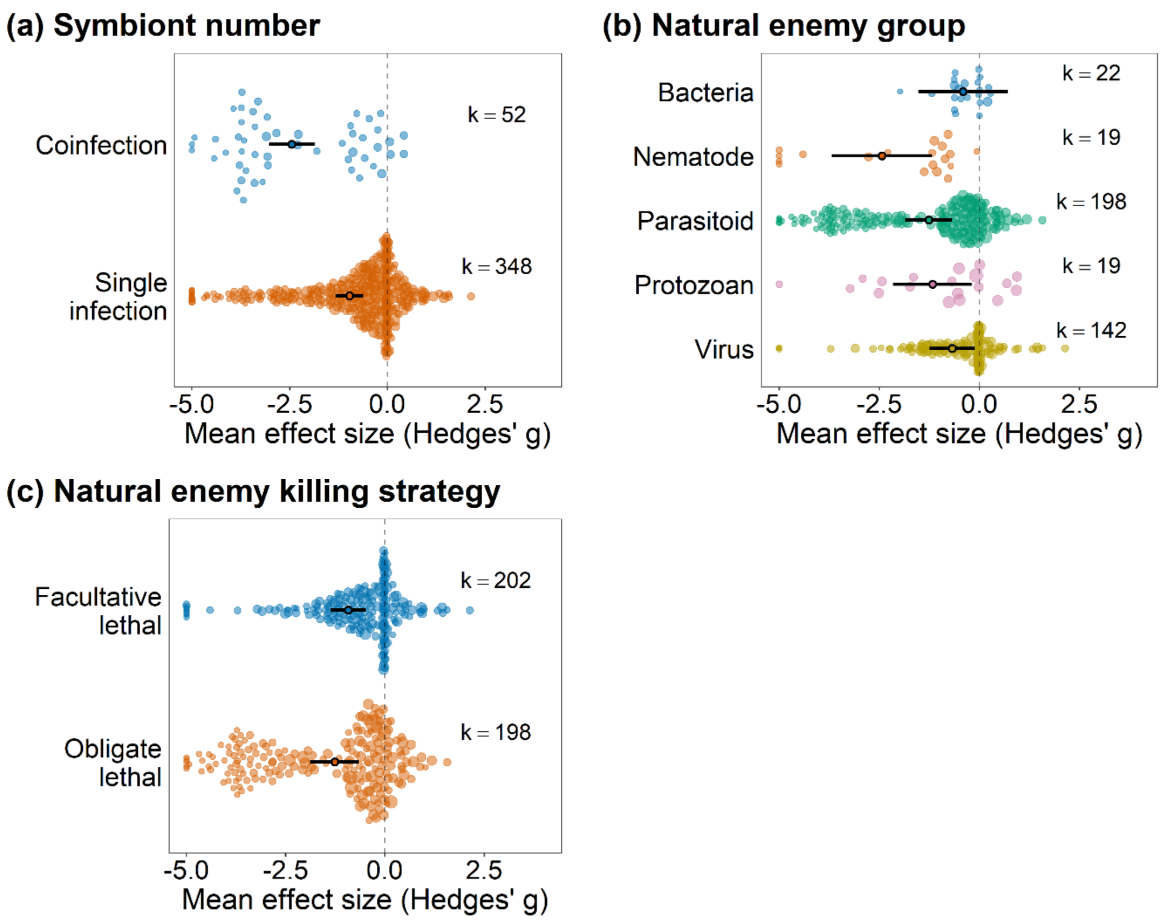

**Figure S4.** Effect of defensive symbionts on the fitness of natural enemies attacking symbiont-infected hosts. (a) model including symbiont number as moderator, (b) model including natural enemy group as moderator, and (c) model including natural enemy killing strategy as moderator. Points are the weighted mean effect sizes ± 95% confidence intervals. Each colored circle represents an effect size, scaled according to precision (inverse of the sampling error variance). 'k' is the number of effect sizes. Thirteen points had values below −5, which expanded the x-axis range and compressed the visual spacing of the remaining points. For visualization purposes, these points are shown at −5. All data points were included in the statistical analyses.

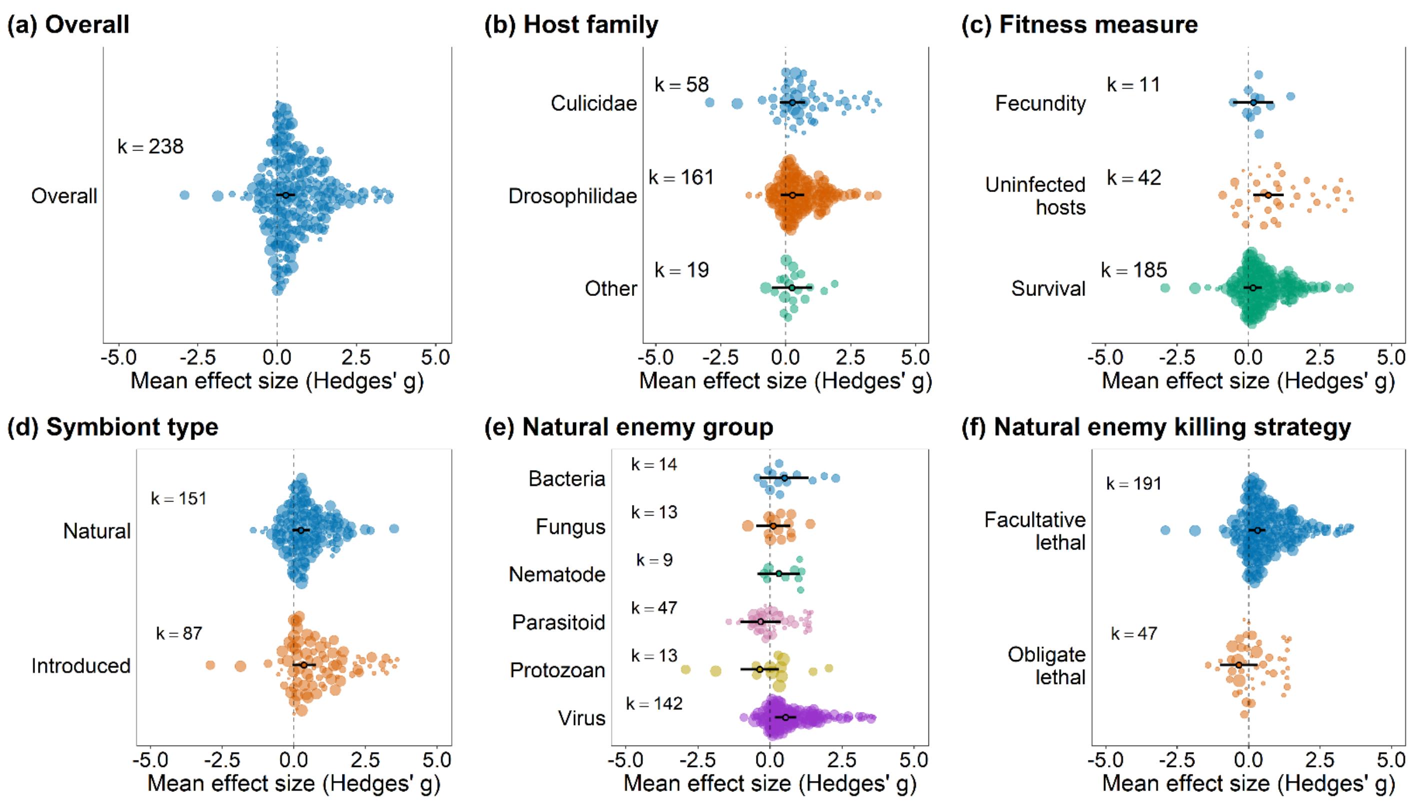

**Figure S5.** Sub-analysis: *Wolbachia* only. Effect of defensive symbionts on the fitness of hosts that are under attack of natural enemies. (a) overall model, (b) model including host family as moderator, (c) model including fitness measure as moderator, (d) model including symbiont type (i.e., if the symbiont naturally infects the host species or if it was artificially introduced) as moderator, (e) model including natural enemy group as moderator, and (f) model including natural enemy killing strategy as moderator. Positive values indicate positive effects on host fitness. Hence, positive values indicate higher fecundity, survival, and proportion of hosts that were uninfected (i.e., uninfected hosts - number of individuals that were unsuccessfully parasitized or infected by natural enemies). Negative values indicate that symbionts have a negative effect on host fitness. Points are the weighted mean effect sizes ± 95% confidence intervals. Each colored circle represents an effect size, scaled according to precision (inverse of the sampling error variance). ‘k’ is the number of effect sizes.

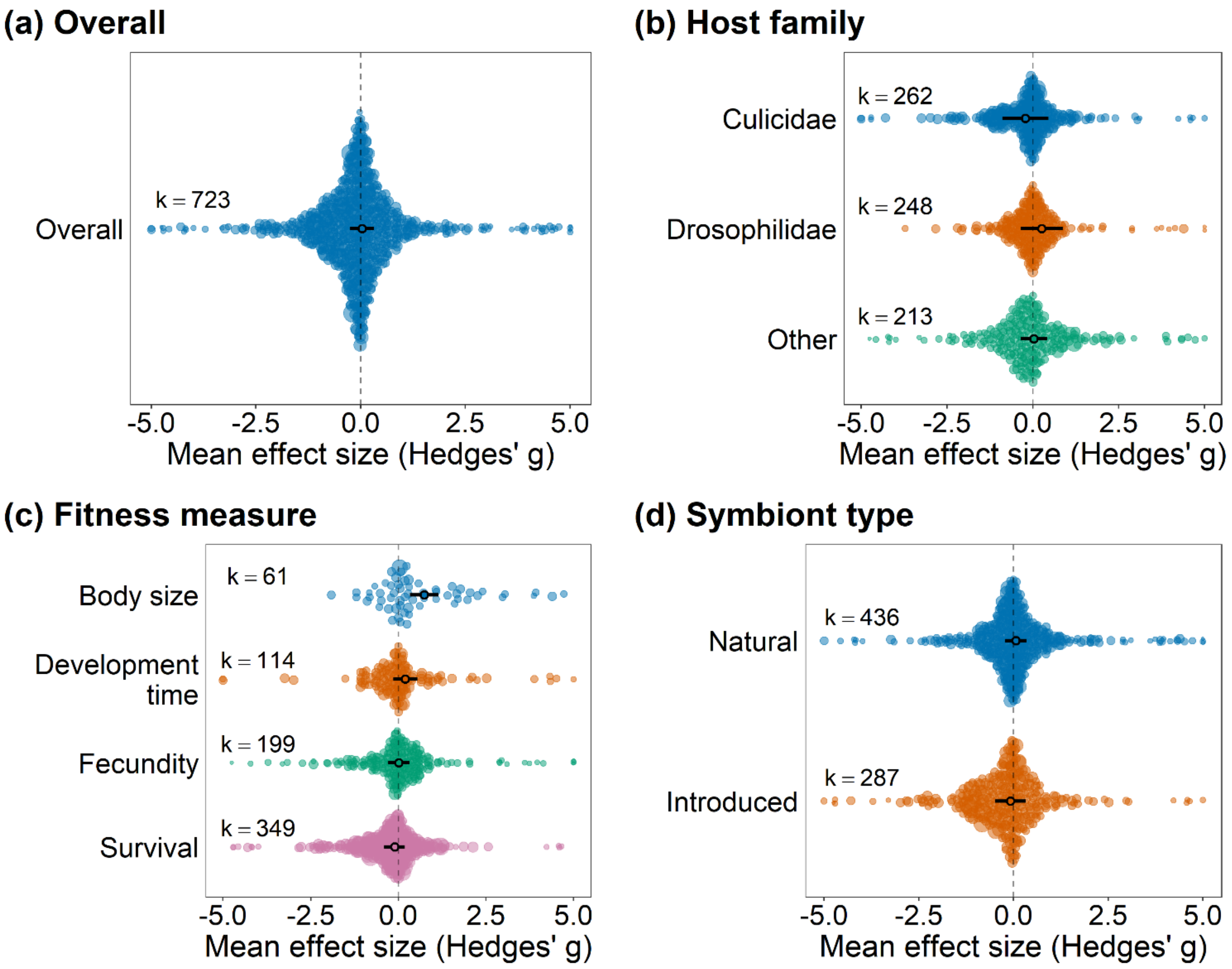

**Figure S6.** Sub-analysis: *Wolbachia* only. Effect of defensive symbionts on the fitness of hosts that are in the absence of natural enemies. (a) overall model, (b) model including host family as moderator, (c) model including fitness measure as moderator, and (d) model including symbiont type (i.e., if the symbiont naturally infects the host species or if it was artificially introduced) as moderator. Positive values indicate positive effects on host fitness. Hence, positive values indicate higher fecundity, survival, body size and shorter development time. Negative values indicate that symbionts have a negative effect on fitness. Points are the weighted mean effect sizes ± 95% confidence intervals. Each colored circle represents an effect size, scaled according to precision (inverse of the sampling error variance). 'k' is the number of effect sizes. Two points fell below −5 and four points fell above +5; for visualization purposes only, these points are shown at ±5. All data points were included in the statistical analyses.

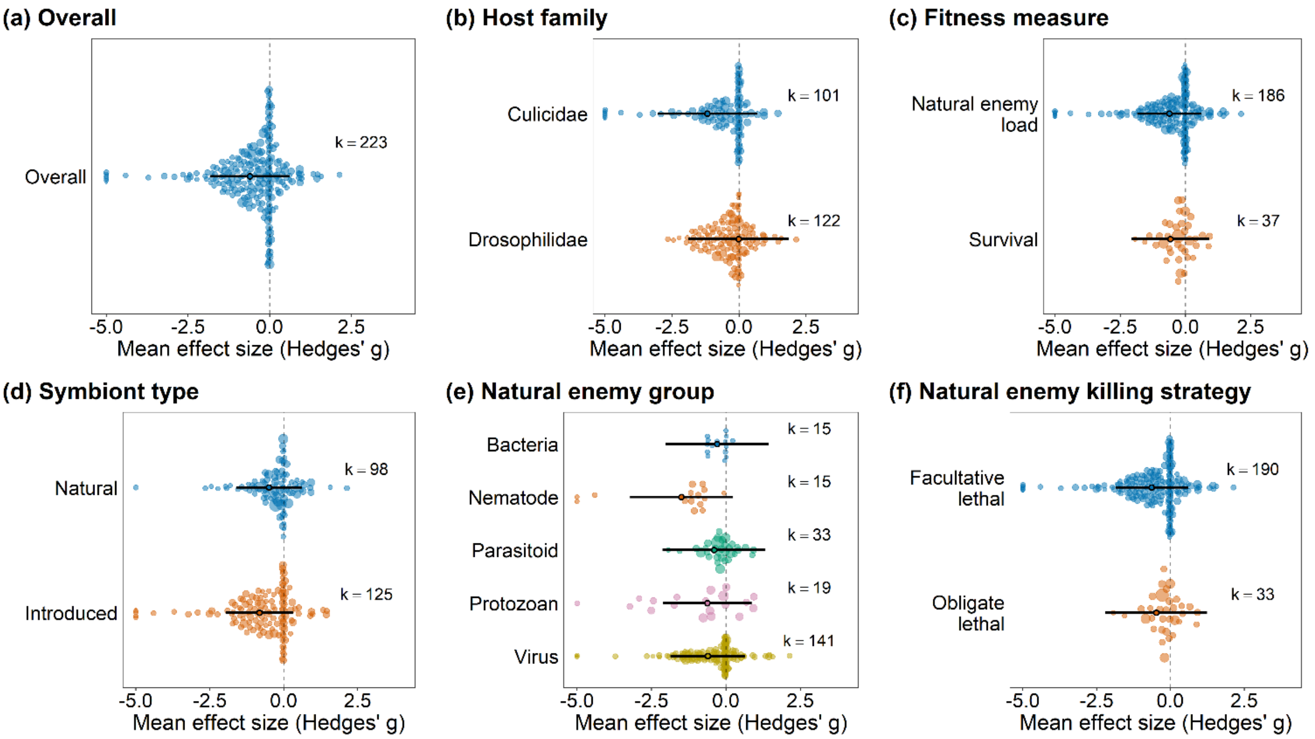

**Figure S7.** Sub-analysis: *Wolbachia* only. Effect of defensive symbionts on the fitness of natural enemies infecting symbiont-infected hosts. (a) overall model, (b) model including host family as moderator, (c) model including fitness measure as moderator, (d) model including symbiont type (i.e., if the symbiont naturally infects the host species or if it was artificially introduced) as moderator, (e) natural enemy group as moderator, and (f) model including natural enemy killing strategy as moderator. Positive values indicate positive effects on host fitness. Hence, positive values indicate higher survival, and higher natural enemy load (quantity of natural enemies replicating inside the host such as viruses and bacteria). Points are the weighted mean effect sizes ± 95% confidence intervals. Each colored circle represents an effect size, scaled according to precision (inverse of the sampling error variance). ‘k’ is the number of effect sizes. Six points fell below −5; for visualization purposes only, these points are shown at -5. All data points were included in the statistical analyses.

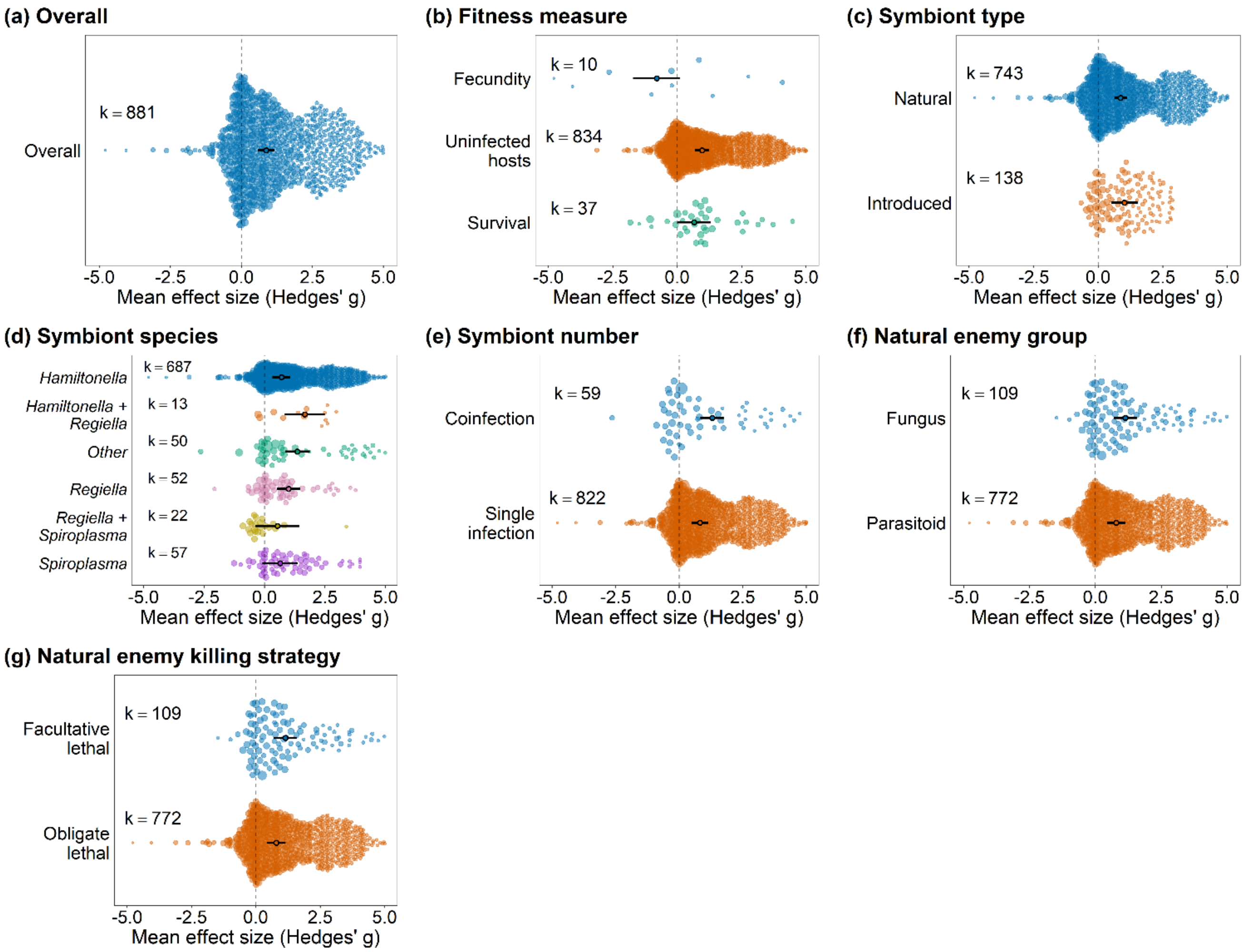

**Figure S8.** Sub-analysis: Aphids only. Effect of defensive symbionts on the fitness of aphids that are under attack of natural enemies. (a) overall model, (b) model including fitness measure as moderator, (c) model including symbiont type (i.e., if the symbiont naturally infects the host species or if it was artificially introduced) as moderator, (d) model including symbiont species as moderator, (e) model including symbiont number as moderator, (f) model including natural enemy group as moderator, and (g) model including natural enemy killing strategy as moderator. Positive values indicate positive effects on host fitness. Hence, positive values indicate higher fecundity, survival, and proportion of hosts that were uninfected (i.e., uninfected hosts - number of individuals that were unsuccessfully parasitized or infected). Negative values indicate that symbionts have a negative effect on host fitness. Points are the weighted mean effect sizes ± 95% confidence intervals. Each colored circle represents an effect size, scaled according to precision (inverse of the sampling error variance). ‘k’ is the number of effect sizes.

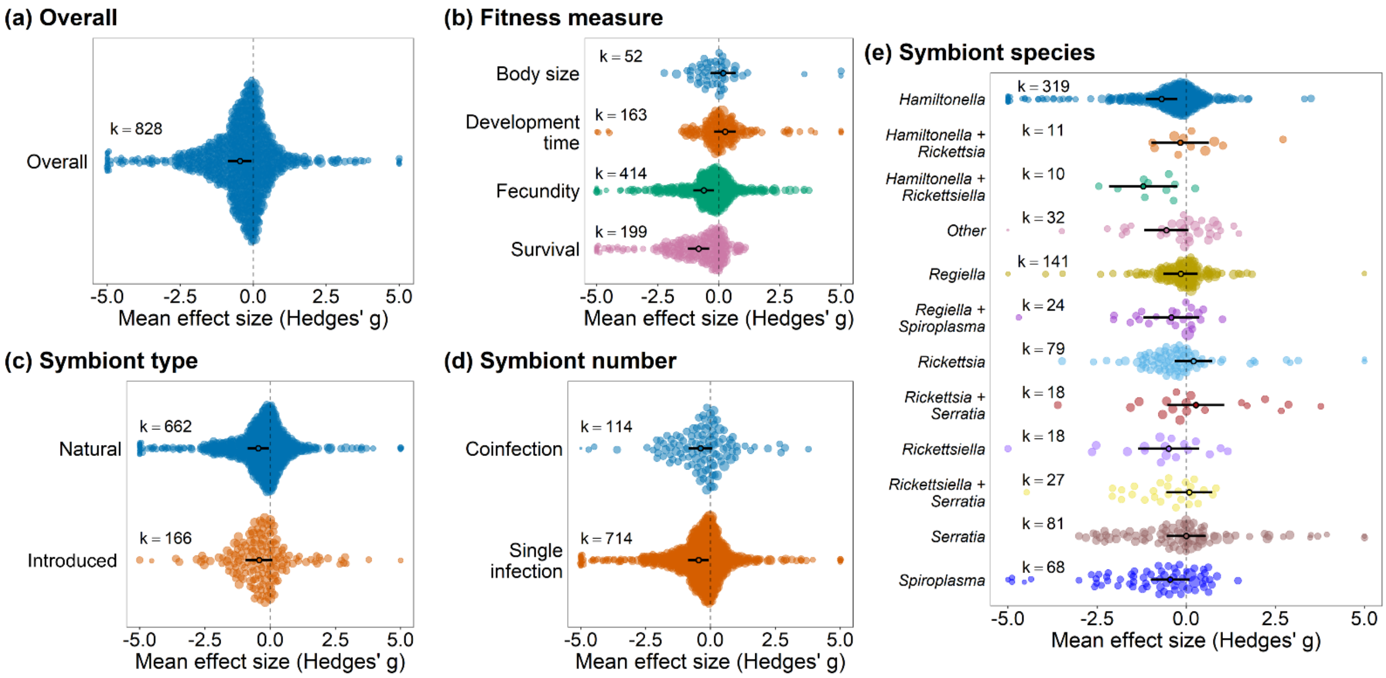

**Figure S9.** Sub-analysis: Aphids only. Effect of defensive symbionts on the fitness of aphids that are in the absence of natural enemies. (a) overall model, (b) model including fitness measure as moderator, (c) model including symbiont type (i.e., if the symbiont naturally infects the host species or if it was artificially introduced) as moderator, (d) model including symbiont number as moderator, and (e) model including symbiont species as moderator. Positive values indicate positive effects on host fitness. Hence, positive values indicate higher fecundity, survival, body size and shorter development time. Negative values indicate that symbionts have a negative effect on fitness. Points are the weighted mean effect sizes ± 95% confidence intervals. Each colored circle represents an effect size, scaled according to precision (inverse of the sampling error variance). 'k' is the number of effect sizes. Fourteen points fell below −5 and four points fell above +5; for visualization purposes only, these points are shown at ±5. All data points were included in the statistical analyses.

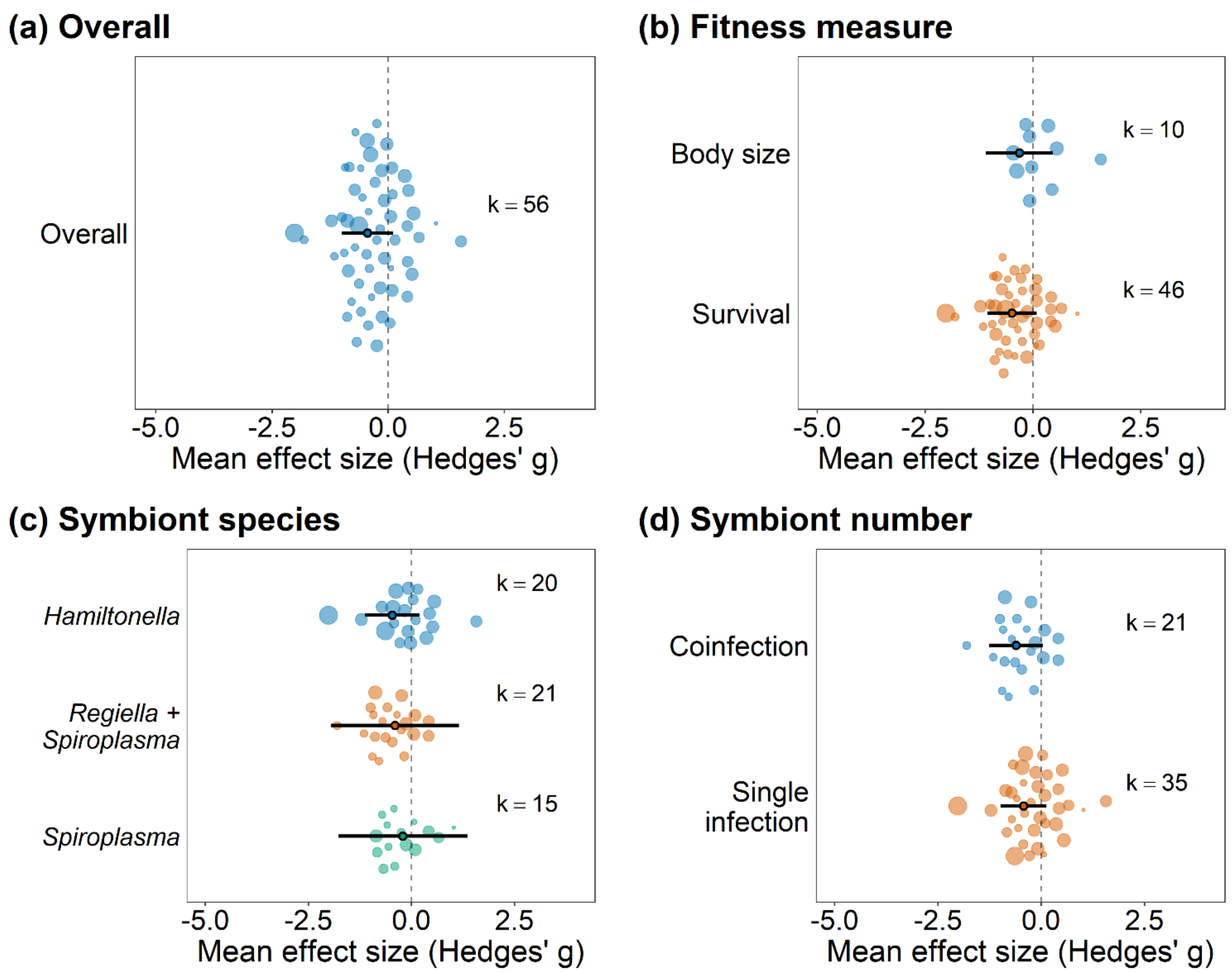

**Figure S10.** Sub-analysis: Aphids only. Effect of defensive symbionts on the fitness of natural enemies infecting symbiont-infected hosts. (a) overall model, (b) model including fitness measure as moderator, (c) model including symbiont species as moderator, and (d) model including symbiont number as moderator. Positive values indicate positive effects on host fitness. Hence, positive values indicate bigger body size and higher survival. Points are the weighted mean effect sizes ± 95% confidence intervals. Each colored circle represents an effect size, scaled according to precision (inverse of the sampling error variance). 'k' is the number of effect sizes.

### Supplementary tables

**Table S1.** Study systems from “protection” dataset that were included in the meta-analysis.

| Host species | Host family | Symbiont species | Natural enemy group |
|---|---|---|---|
| *Acyrthosiphon pisum* | Aphididae | *Hamiltonella* | Bacteria |
| *Acyrthosiphon pisum* | Aphididae | *Regiella* | Bacteria |
| *Acyrthosiphon pisum* | Aphididae | *Serratia* | Bacteria |
| *Aedes aegypti* | Culicidae | *Wolbachia* | Bacteria |
| *Aphis fabae* | Aphididae | *Serratia* | Bacteria |
| *Armadillidium vulgare* | Armadillidiidae | *Wolbachia* | Bacteria |
| *Drosophila melanogaster* | Drosophilidae | *Spiroplasma* | Bacteria |
| *Porcellio dilatatus* | Porcellionidae | *Wolbachia* | Bacteria |
| *Acyrthosiphon pisum* | Aphididae | *Hamiltonella* | Fungus |
| *Acyrthosiphon pisum* | Aphididae | *Hamiltonella + Regiella* | Fungus |
| *Acyrthosiphon pisum* | Aphididae | *Hamiltonella + Rickettsia* | Fungus |
| *Acyrthosiphon pisum* | Aphididae | *Hamiltonella + Spiroplasma* | Fungus |
| *Acyrthosiphon pisum* | Aphididae | *Regiella* | Fungus |
| *Acyrthosiphon pisum* | Aphididae | *Rickettsia* | Fungus |
| *Acyrthosiphon pisum* | Aphididae | *Rickettsiella* | Fungus |
| *Acyrthosiphon pisum* | Aphididae | *Spiroplasma* | Fungus |
| *Aedes aegypti* | Culicidae | *Wolbachia* | Fungus |
| *Aphis fabae* | Aphididae | *Serratia* | Fungus |
| *Drosophila melanogaster* | Drosophilidae | *Wolbachia* | Fungus |
| *Drosophila simulans* | Drosophilidae | *Wolbachia* | Fungus |
| *Pentalonia nigronervosa* | Aphididae | *Wolbachia* | Fungus |
| *Sitobion avenae* | Aphididae | *Hamiltonella* | Fungus |
| *Sitobion avenae* | Aphididae | *Hamiltonella + Rickettsia* | Fungus |
| *Sitobion avenae* | Aphididae | *Hamiltonella + Spiroplasma* | Fungus |
| *Sitobion avenae* | Aphididae | *Regiella* | Fungus |

| | | | |
|---|---|---|---|
| *Sitobion avenae* | Aphididae | *Rickettsia* | Fungus |
| *Sitobion avenae* | Aphididae | *Spiroplasma* | Fungus |
| *Tetranychus urticae* | Tetranychidae | *Wolbachia* | Fungus |
| *Aedes aegypti* | Culicidae | *Wolbachia* | Nematode |
| *Aedes pseudoscutellaris* | Culicidae | *Wolbachia* | Nematode |
| *Anopheles gambiae* | Culicidae | *Wolbachia* | Nematode |
| *Drosophila falleni* | Drosophilidae | *Spiroplasma* | Nematode |
| *Drosophila melanogaster* | Drosophilidae | *Wolbachia* | Nematode |
| *Drosophila neotestacea* | Drosophilidae | *Spiroplasma* | Nematode |
| *Drosophila neotestacea* | Drosophilidae | *Wolbachia* | Nematode |
| *Drosophila orientacea* | Drosophilidae | *Spiroplasma* | Nematode |
| *Drosophila putrida* | Drosophilidae | *Spiroplasma* | Nematode |
| *Drosophila testacea* | Drosophilidae | *Spiroplasma* | Nematode |
| *Acyrthosiphon pisum* | Aphididae | *Fukatsuia* | Parasitoid |
| *Acyrthosiphon pisum* | Aphididae | *Fukatsuia + Hamiltonella* | Parasitoid |
| *Acyrthosiphon pisum* | Aphididae | *Hamiltonella* | Parasitoid |
| *Acyrthosiphon pisum* | Aphididae | *Hamiltonella + Regiella* | Parasitoid |
| *Acyrthosiphon pisum* | Aphididae | *Hamiltonella + Rickettsiella* | Parasitoid |
| *Acyrthosiphon pisum* | Aphididae | *Hamiltonella + Serratia* | Parasitoid |
| *Acyrthosiphon pisum* | Aphididae | *Hamiltonella + Spiroplasma* | Parasitoid |
| *Acyrthosiphon pisum* | Aphididae | *Regiella* | Parasitoid |
| *Acyrthosiphon pisum* | Aphididae | *Regiella + Spiroplasma* | Parasitoid |
| *Acyrthosiphon pisum* | Aphididae | *Rickettsia* | Parasitoid |
| *Acyrthosiphon pisum* | Aphididae | *Rickettsiella* | Parasitoid |
| *Acyrthosiphon pisum* | Aphididae | *Serratia* | Parasitoid |
| *Acyrthosiphon pisum* | Aphididae | *Serratia + Spiroplasma* | Parasitoid |
| *Acyrthosiphon pisum* | Aphididae | *Spiroplasma* | Parasitoid |
| *Aphis craccivora* | Aphididae | *Arsenophonus* | Parasitoid |
| *Aphis craccivora* | Aphididae | *Hamiltonella* | Parasitoid |
| *Aphis fabae* | Aphididae | *Hamiltonella* | Parasitoid |

| | | | |
|---|---|---|---|
| *Aphis fabae* | Aphididae | *Regiella* | Parasitoid |
| *Aphis fabae* | Aphididae | *Serratia* | Parasitoid |
| *Aphis gossypii* | Aphididae | *Arsenophonus + Hamiltonella* | Parasitoid |
| *Drosophila hydei* | Drosophilidae | *Spiroplasma* | Parasitoid |
| *Drosophila melanogaster* | Drosophilidae | *Spiroplasma* | Parasitoid |
| *Drosophila melanogaster* | Drosophilidae | *Spiroplasma + Wolbachia* | Parasitoid |
| *Drosophila melanogaster* | Drosophilidae | *Wolbachia* | Parasitoid |
| *Drosophila neotestacea* | Drosophilidae | *Spiroplasma* | Parasitoid |
| *Drosophila simulans* | Drosophilidae | *Wolbachia* | Parasitoid |
| *Macrosiphum euphorbiae* | Aphididae | *Hamiltonella* | Parasitoid |
| *Myzus persicae* | Aphididae | *Hamiltonella* | Parasitoid |
| *Myzus persicae* | Aphididae | *Regiella* | Parasitoid |
| *Pentalonia nigronervosa* | Aphididae | *Wolbachia* | Parasitoid |
| *Sitobion avenae* | Aphididae | *Hamiltonella* | Parasitoid |
| *Sitobion avenae* | Aphididae | *Regiella* | Parasitoid |
| *Anopheles gambiae* | Culicidae | *Wolbachia* | Protozoan |
| *Culex pipiens* | Culicidae | *Wolbachia* | Protozoan |
| *Drosophila melanogaster* | Drosophilidae | *Wolbachia* | Protozoan |
| *Glossina fuscipes* | Glossinidae | *Spiroplasma* | Protozoan |
| *Aedes aegypti* | Culicidae | *Wolbachia* | Virus |
| *Aedes albopictus* | Culicidae | *Wolbachia* | Virus |
| *Aedes notoscriptus* | Culicidae | *Wolbachia* | Virus |
| *Bemisia tabaci* | Aleyrodidae | *Arsenophonus + Rickettsia* | Virus |
| *Culex tarsalis* | Culicidae | *Wolbachia* | Virus |
| *Drosophila ananassae* | Drosophilidae | *Wolbachia* | Virus |
| *Drosophila baimaii* | Drosophilidae | *Wolbachia* | Virus |
| *Drosophila bifasciata* | Drosophilidae | *Wolbachia* | Virus |
| *Drosophila melanogaster* | Drosophilidae | *Wolbachia* | Virus |
| *Drosophila nikananu* | Drosophilidae | *Wolbachia* | Virus |
| *Drosophila prosaltans* | Drosophilidae | *Wolbachia* | Virus |

| | | | |
|---|---|---|---|
| *Drosophila santomea* | Drosophilidae | *Wolbachia* | Virus |
| *Drosophila sechellia* | Drosophilidae | *Wolbachia* | Virus |
| *Drosophila simulans* | Drosophilidae | *Wolbachia* | Virus |
| *Drosophila stonei* | Drosophilidae | *Wolbachia* | Virus |
| *Drosophila sturtevanti* | Drosophilidae | *Wolbachia* | Virus |
| *Drosophila suzukii* | Drosophilidae | *Wolbachia* | Virus |
| *Drosophila teissieri* | Drosophilidae | *Wolbachia* | Virus |
| *Drosophila triauraria* | Drosophilidae | *Wolbachia* | Virus |
| *Drosophila tropicalis* | Drosophilidae | *Wolbachia* | Virus |

**Table S2.** Study systems from "cost to hosts" dataset that were included in the meta-analysis.

| Host species | Host family | Symbiont species |
|---|---|---|
| *Sitobion avenae* | Aphididae | *Hamiltonella* |
| *Acyrthosiphon pisum* | Aphididae | *Hamiltonella* |
| *Acyrthosiphon pisum* | Aphididae | *Hamiltonella + Serratia* |
| *Sitobion avenae* | Aphididae | *Regiella* |
| *Acyrthosiphon pisum* | Aphididae | *Regiella* |
| *Aphis fabae* | Aphididae | *Serratia* |
| *Aphis citricidus* | Aphididae | *Spiroplasma* |
| *Aphis gossypii* | Aphididae | *Arsenophonus + Hamiltonella* |
| *Acyrthosiphon pisum* | Aphididae | *Serratia* |
| *Aphis fabae* | Aphididae | *Hamiltonella* |
| *Acyrthosiphon pisum* | Aphididae | *Rickettsia* |
| *Acyrthosiphon kondoi* | Aphididae | *Rickettsia* |
| *Acyrthosiphon pisum* | Aphididae | *Rickettsia +Serratia* |
| *Acyrthosiphon kondoi* | Aphididae | *Serratia* |
| *Macrosiphum euphorbiae* | Aphididae | *Hamiltonella* |
| *Acyrthosiphon pisum* | Aphididae | *Fukatsuia + Hamiltonella* |
| *Acyrthosiphon pisum* | Aphididae | *Fukatsuia* |
| *Aphis craccivora* | Aphididae | *Hamiltonella* |
| *Myzus persicae* | Aphididae | *Rickettsiella* |
| *Pentalonia nigronervosa* | Aphididae | *Wolbachia* |
| *Aphis fabae* | Aphididae | *Regiella* |
| *Myzus persicae* | Aphididae | *Regiella* |
| *Acyrthosiphon pisum* | Aphididae | *Hamiltonella + Rickettsiella* |
| *Acyrthosiphon pisum* | Aphididae | *Rickettsiella* |
| *Acyrthosiphon pisum* | Aphididae | *Hamiltonella + Spiroplasma* |
| *Sitobion avenae* | Aphididae | *Hamiltonella + Spiroplasma* |
| *Acyrthosiphon pisum* | Aphididae | *Spiroplasma* |
| *Sitobion avenae* | Aphididae | *Spiroplasma* |

| | | |
|---|---|---|
| *Acyrthosiphon pisum* | Aphididae | *Hamiltonella + Rickettsia* |
| *Sitobion avenae* | Aphididae | *Hamiltonella + Rickettsia* |
| *Sitobion avenae* | Aphididae | *Rickettsia* |
| *Acyrthosiphon pisum* | Aphididae | *Regiella + Spiroplasma* |
| *Acyrthosiphon pisum* | Aphididae | *Hamiltonella + Regiella* |
| *Sitobion miscanthi* | Aphididae | *Hamiltonella* |
| *Sitobion miscanthi* | Aphididae | *Regiella + Spiroplasma* |
| *Acyrthosiphon pisum* | Aphididae | *Rickettsiella + Serratia* |
| *Aedes aegypti* | Culicidae | *Wolbachia* |
| *Anopheles stephensi* | Culicidae | *Wolbachia* |
| *Aedes albopictus* | Culicidae | *Wolbachia* |
| *Aedes polynesiensis* | Culicidae | *Wolbachia* |
| *Culex pipiens* | Culicidae | *Wolbachia* |
| *Ochlerotatus fluviatilis* | Culicidae | *Wolbachia* |
| *Anopheles gambiae* | Culicidae | *Wolbachia* |
| *Drosophila simulans* | Drosophilidae | *Wolbachia* |
| *Drosophila melanogaster* | Drosophilidae | *Wolbachia* |
| *Drosophila melanogaster* | Drosophilidae | *Spiroplasma* |
| *Drosophila neotestacea* | Drosophilidae | *Spiroplasma* |
| *Drosophila hydei* | Drosophilidae | *Spiroplasma* |
| *Drosophila melanogaster* | Drosophilidae | *Spiroplasma + Wolbachia* |
| *Drosophila suzukii* | Drosophilidae | *Wolbachia* |
| *Drosophila willistoni* | Drosophilidae | *Spiroplasma* |
| *Drosophila subpulchrella* | Drosophilidae | *Wolbachia* |
| *Drosophila falleni* | Drosophilidae | *Spiroplasma* |
| *Drosophila orientacea* | Drosophilidae | *Spiroplasma* |
| *Drosophila putrida* | Drosophilidae | *Spiroplasma* |
| *Drosophila neotestacea* | Drosophilidae | *Spiroplasma + Wolbachia* |
| *Drosophila neotestacea* | Drosophilidae | *Wolbachia* |
| *Drosophila mauritiana* | Drosophilidae | *Wolbachia* |

| | | |
|---|---|---|
| *Drosophila paulistorum* | Drosophilidae | *Wolbachia* |
| *Drosophila willistoni* | Drosophilidae | *Wolbachia* |
| *Drosophila pseudotakahashii* | Drosophilidae | *Wolbachia* |
| *Drosophila sturtevanti* | Drosophilidae | *Wolbachia* |
| *Drosophila innubila* | Drosophilidae | *Wolbachia* |
| *Bemisia tabaci* | Aleyrodidae | *Wolbachia* |
| *Sogatella furcifera* | Delphacidae | *Cardinium* + *Wolbachia* |
| *Tetranychus truncatus* | Tetranychidae | *Cardinium* + *Wolbachia* |
| *Bemisia tabaci* | Aleyrodidae | *Hamiltonella* + *Rickettsia* |
| *Bemisia tabaci* | Aleyrodidae | *Hamiltonella* |
| *Bemisia tabaci* | Aleyrodidae | *Arsenophonus* + *Rickettsia* |
| *Bemisia tabaci* | Aleyrodidae | *Rickettsia* |
| *Chrysoperla carnea* | Chrysopidae | *Rickettsia* |
| *Tetranychus truncatus* | Tetranychidae | *Spiroplasma* |
| *Tetranychus truncatus* | Tetranychidae | *Spiroplasma* + *Wolbachia* |
| *Homona magnanima* | Tortricidae | *Wolbachia* |
| *Ceratitis capitata* | Tephritidae | *Wolbachia* |
| *Exorista sorbillans* | Tachinidae | *Wolbachia* |
| *Callosobruchus chinensis* | Chrysomelidae | *Wolbachia* |
| *Ephestia kuehniella* | Pyralidae | *Wolbachia* |
| *Tetranychus urticae* | Tetranychidae | *Wolbachia* |
| *Sogatella furcifera* | Delphacidae | *Wolbachia* |
| *Tetranychus truncatus* | Tetranychidae | *Wolbachia* |
| *Glossina fuscipes* | Glossinidae | *Spiroplasma* |
| *Oryzaephilus surinamensis* | Silvanidae | *Wolbachia* |
| *Hyposoter horticola* | Ichneumonidae | *Wolbachia* |
| *Leptopilina heterotoma* | Figitidae | *Wolbachia* |
| *Laodelphax striatellus* | Delphacidae | *Wolbachia* |
| *Anagyrus vladimiri* | Encyrtidae | *Wolbachia* |
| *Nasonia vitripennis* | Pteromalidae | *Wolbachia* |

| | | |
|---|---|---|
| *Cordylochernes scorpioides* | Chernetidae | *Wolbachia* |
| *Armadillidium vulgare* | Armadillidiidae | *Wolbachia* |
| *Haematobia irritans exigua* | Muscidae | *Wolbachia* |
| *Harmonia yedoensis* | Coccinellidae | *Spiroplasma* |
| *Spalangia endius* | Spalangiidae | *Wolbachia* |
| *Spalangia endius* | Spalangiidae | *Rickettsia* |
| *Spalangia endius* | Spalangiidae | *Rickettsia + Wolbachia* |
| *Monomorium pharaonis* | Formicidae | *Wolbachia* |
| *Aphytis melinus* | Aphelinidae | *Wolbachia* |
| *Encarsia inaron* | Aphelinidae | *Cardinium + Wolbachia* |
| *Encarsia inaron* | Aphelinidae | *Wolbachia* |
| *Trichogramma ostriniae* | Trichogrammatidae | *Wolbachia* |
| *Laodelphax striatellus* | Delphacidae | *Spiroplasma + Wolbachia* |
| *Laodelphax striatellus* | Delphacidae | *Spiroplasma* |
| *Velarifictorus aspersus* | Gryllidae | *Wolbachia* |
| *Pezothrips kellyanus* | Thripidae | *Cardinium + Wolbachia* |

**Table S3.** Study systems from "cost to natural enemy" dataset that were included in the meta-analysis.

| Host species | Host family | Symbiont species | Natural enemy group |
|---|---|---|---|
| *Aedes aegypti* | Culicidae | *Wolbachia* | Bacteria |
| *Armadillidium vulgare* | Armadillidiidae | *Wolbachia* | Bacteria |
| *Bemisia tabaci* | Aleyrodidae | *Rickettsia* | Bacteria |
| *Drosophila melanogaster* | Drosophilidae | *Wolbachia* | Bacteria |
| *Porcellio dilatatus* | Porcellionidae | *Wolbachia* | Bacteria |
| *Aedes aegypti* | Culicidae | *Wolbachia* | Nematode |
| *Aedes polynesiensis* | Culicidae | *Wolbachia* | Nematode |
| *Anopheles gambiae* | Culicidae | *Wolbachia* | Nematode |
| *Drosophila falleni* | Drosophilidae | *Spiroplasma* | Nematode |
| *Drosophila orientacea* | Drosophilidae | *Spiroplasma* | Nematode |
| *Drosophila putrida* | Drosophilidae | *Spiroplasma* | Nematode |
| *Drosophila testacea* | Drosophilidae | *Spiroplasma* | Nematode |
| *Acyrthosiphon pisum* | Aphididae | *Fukatsuia + Hamiltonella* | Parasitoid |
| *Acyrthosiphon pisum* | Aphididae | *Hamiltonella* | Parasitoid |
| *Acyrthosiphon pisum* | Aphididae | *Regiella + Spiroplasma* | Parasitoid |
| *Acyrthosiphon pisum* | Aphididae | *Spiroplasma* | Parasitoid |
| *Aphis craccivora* | Aphididae | *Hamiltonella* | Parasitoid |
| *Aphis fabae* | Aphididae | *Hamiltonella* | Parasitoid |
| *Aphis fabae* | Aphididae | *Serratia* | Parasitoid |
| *Bemisia tabaci* | Aleyrodidae | *Wolbachia* | Parasitoid |
| *Drosophila hydei* | Drosophilidae | *Spiroplasma* | Parasitoid |
| *Drosophila melanogaster* | Drosophilidae | *Spiroplasma* | Parasitoid |
| *Drosophila melanogaster* | Drosophilidae | *Spiroplasma + Wolbachia* | Parasitoid |
| *Drosophila melanogaster* | Drosophilidae | *Wolbachia* | Parasitoid |
| *Drosophila neotestacea* | Drosophilidae | *Spiroplasma* | Parasitoid |
| *Drosophila simulans* | Drosophilidae | *Wolbachia* | Parasitoid |
| *Hyposoter horticola* | Ichneumonidae | *Wolbachia* | Parasitoid |
| *Sitobion avenae* | Aphididae | *Regiella* | Parasitoid |

| | | | |
|---|---|---|---|
| *Aedes aegypti* | Culicidae | *Wolbachia* | Protozoan |
| *Aedes fluviatilis* | Culicidae | *Wolbachia* | Protozoan |
| *Anopheles gambiae* | Culicidae | *Wolbachia* | Protozoan |
| *Anopheles stephensi* | Culicidae | *Wolbachia* | Protozoan |
| *Culex pipiens* | Culicidae | *Wolbachia* | Protozoan |
| *Aedes aegypti* | Culicidae | *Wolbachia* | Virus |
| *Aedes albopictus* | Culicidae | *Wolbachia* | Virus |
| *Aedes notoscripyus* | Culicidae | *Wolbachia* | Virus |
| *Aedes polynesiensis* | Culicidae | *Wolbachia* | Virus |
| *Bemisia tabaci* | Aleyrodidae | *Arsenophonus* + *Rickettsia* | Virus |
| *Culex pipiens* | Culicidae | *Wolbachia* | Virus |
| *Drosophila ananassae* | Drosophilidae | *Wolbachia* | Virus |
| *Drosophila baimaii* | Drosophilidae | *Wolbachia* | Virus |
| *Drosophila melanogaster* | Drosophilidae | *Wolbachia* | Virus |
| *Drosophila prosaltans* | Drosophilidae | *Wolbachia* | Virus |
| *Drosophila sechellia* | Drosophilidae | *Wolbachia* | Virus |
| *Drosophila simulans* | Drosophilidae | *Wolbachia* | Virus |
| *Drosophila sturtevanti* | Drosophilidae | *Wolbachia* | Virus |
| *Drosophila suzukii* | Drosophilidae | *Wolbachia* | Virus |
| *Drosophila teissieri* | Drosophilidae | *Wolbachia* | Virus |
| *Drosophila triauraria* | Drosophilidae | *Wolbachia* | Virus |
| *Drosophila tropicalis* | Drosophilidae | *Wolbachia* | Virus |

**Table S4.** List of all fitness measures extracted from the studies to calculate the mean effect sizes. Measures are grouped in six broad categories.

| Category | Fitness measure |
|---|---|
| Survival | Longevity (mean number of days alive)<br>Adult survival (% or number alive)<br>Larval survival (% or number alive)<br>Pupa survival (% or number alive)<br>Egg hatch (% or number of eggs that hatched) |
| Uninfected hosts | Non-parasitized (% or number of individuals unsuccessfully parasitized by parasitoids)<br>Non-infected (% or number of individuals unsuccessfully infected by other natural enemies) |
| Fecundity | Fecundity (number of eggs or offspring)<br>Fertility (number of ovarioles or oocytes) |
| Body size | Wing length<br>Wing width<br>Cephalothorax length<br>Thorax length<br>Mass (adult weight) |
| Development time | Development time (Time taken to develop from one life stage into another. Examples: larva to pupa, pupa to adult, larva to adult, nymph to adult) |
| Natural enemy load | Viral load<br>Bacterial load<br>Ookinet load<br>Oocyst load<br>Sporozoite load<br>Worm load |

**Table S5.** Variables collected from the original studies to be used as moderators in the models.

| Moderator | Moderator level |
|---|---|
| Host family | Aphididae |
| | Culicidae |
| | Drosophilidae |
| | Other families |
| Symbiont type | Natural |
| | Introduced |
| Symbiont species | *Cardinium + Wolbachia* |
| | *Hamiltonella* |
| | *Hamiltonella +Rickettsia* |
| | *Hamiltonella + Rickttsiella* |
| | Other species |
| | *Regiella* |
| | *Regiella + Spiroplasma* |
| | *Rickettsia* |
| | *Rickettsiella* |
| | *Serratia* |
| | *Spiroplasma* |
| | *Spiroplasma + Wolbachia* |
| | *Wolbachia* |
| Natural enemy group | Bacteria |
| | Fungus |
| | Nematode |
| | Parasitoid |
| | Protozoan |
| | Virus |
| Fitness measure | Survival |
| | Uninfected hosts (% or number of uninfected individuals) |
| | Fecundity |
| | Body size |
| | Development time |
| | Natural enemy load |

**Table S6.** Main analysis: multilevel meta-analytical models performed in this study, specifying their random and fixed factors (moderators) and moderator levels.

| Data | Model | Random factors | Fixed factors (moderators) | Moderator levels |
|---|---|---|---|---|
| **Protection** | **Overall** | Study ID + effect size ID + host species + host phylogeny | - | - |
| | **Host family** | Study ID + effect size ID + host species + host phylogeny | Host family | Aphididae<br>Culicidae<br>Drosophilidae<br>Other families |
| | **Symbiont type** | Study ID + effect size ID + host species + host phylogeny | Symbiont type | Natural<br>Introduced |
| | **Symbiont species** | Study ID + effect size ID + host species + host phylogeny | Symbiont species | *Hamiltonella*<br>*Hamiltonella + Regiella*<br>Other species<br>*Regiella*<br>*Regiella + Spiroplasma*<br>*Spiroplasma*<br>*Spiroplasma + Wolbachia*<br>*Wolbachia* |
| | **Fitness measure** | Study ID + effect size ID + host species + host phylogeny | Fitness measure | Fecundity<br>Survival<br>Uninfected hosts |
| | **Natural enemy group** | Study ID + effect size ID + host species + host phylogeny | Natural enemy group | Bacteria<br>Fungus<br>Nematode |

| | | | | |
|---|---|---|---|---|
| | | | | Parasitoid<br>Protozoan<br>Virus |
| | **Symbiont number** | Study ID + effect size ID + host species + host phylogeny | Symbiont number | Single<br>Coinfection |
| | **Natural enemy killing strategy** | Study ID + effect size ID + host species + host phylogeny | Natural enemy killing strategy | Facultative lethal<br>Obligate lethal |
| **Cost to hosts** | **Overall** | Study ID + effect size ID + host species + host phylogeny | - | - |
| | **Host family** | Study ID + effect size ID + host species + host phylogeny | Host family | Aphididae<br>Culicidae<br>Drosophilidae<br>Other families |
| | **Symbiont type** | Study ID + effect size ID + host species + host phylogeny | Symbiont type | Natural<br>Introduced |
| | **Symbiont species** | Study ID + effect size ID + host species + host phylogeny | Symbiont species | *Cardinium* + *Wolbachia*<br>*Hamiltonella*<br>*Hamiltonella* +*Rickettsia*<br>*Hamiltonella* + *Rickttsiella*<br>Other species<br>*Regiella*<br>*Regiella* + *Spiroplasma*<br>*Rickettsia*<br>*Rickettsiella*<br>*Serratia*<br>*Spiroplasma*<br>*Spiroplasma* + *Wolbachia*<br>*Wolbachia* |
| | **Fitness measure** | Study ID + effect size ID + host species + host phylogeny | Fitness measure | Body size |

| | | | | |
|---|---|---|---|---|
| | | | | Development time<br>Fecundity<br>Survival |
| | **Symbiont number** | Study ID + effect size ID + host species + host phylogeny | Symbiont number | Single<br>Coinfection |
| **Cost to natural enemy** | **Overall** | Study ID + effect size ID + host species + host phylogeny | - | - |
| | **Host family** | Study ID + effect size ID + host species + host phylogeny | Host family | Aphididae<br>Culicidae<br>Drosophilidae<br>Other families |
| | **Symbiont type** | Study ID + effect size ID + host species + host phylogeny | Symbiont type | Natural<br>Introduced |
| | **Symbiont species** | Study ID + effect size ID + host species + host phylogeny | Symbiont species | *Hamiltonella*<br>Other species<br>*Spiroplasma* |
| | **Fitness measure** | Study ID + effect size ID + host species + host phylogeny | Fitness measure | Body size<br>Natural enemy load<br>Survival |
| | **Natural enemy group** | Study ID + effect size ID + host species + host phylogeny | Natural enemy group | Bacteria<br>Nematode<br>Parasitoid<br>Protozoan<br>Virus |
| | **Symbiont number** | Study ID + effect size ID + host species + host phylogeny | Symbiont number | Single<br>Coinfection |
| | **Natural enemy killing strategy** | Study ID + effect size ID + host species + host phylogeny | Natural enemy killing strategy | Facultative lethal<br>Obligate lethal |

**Table S7.** Subgroup analysis: analyses performed only with *Wolbachia* data. Multilevel meta-analytical models performed in this study, specifying their random and fixed factors (moderators) and moderator levels.

| Data (*Wolbachia*) | Model | Random factors | Fixed factors (moderators) | Moderator levels |
|---|---|---|---|---|
| **Protection** | **Overall** | Study ID + effect size ID + host species + host phylogeny | - | - |
| | **Host family** | Study ID + effect size ID + host species + host phylogeny | Host family | Culicidae<br>Drosophilidae<br>Other families |
| | **Symbiont type** | Study ID + effect size ID + host species + host phylogeny | Symbiont type | Natural<br>Introduced |
| | **Natural enemy group** | Study ID + effect size ID + host species + host phylogeny | Natural enemy group | Bacteria<br>Fungus<br>Nematode<br>Parasitoid<br>Protozoan<br>Virus |
| | **Fitness measure** | Study ID + effect size ID + host species + host phylogeny | Fitness measure | Fecundity<br>Survival<br>Uninfected hosts |
| | **Natural enemy killing strategy** | Study ID + effect size ID + host species + host phylogeny | Natural enemy killing strategy | Facultative lethal<br>Obligate lethal |
| **Cost to hosts** | **Overall** | Study ID + effect size ID + host species + host phylogeny | - | - |

| | | | | |
|---|---|---|---|---|
| | **Host family** | Study ID + effect size ID + host species + host phylogeny | Host family | Culicidae<br>Drosophilidae<br>Other families |
| | **Symbiont type** | Study ID + effect size ID + host species + host phylogeny | Symbiont type | Natural<br>Introduced |
| | **Fitness measure** | Study ID + effect size ID + host species + host phylogeny | Fitness measure | Body size<br>Development time<br>Fecundity<br>Survival |
| **Cost to the natural enemy** | **Overall** | Study ID + effect size ID + host species + host phylogeny | - | - |
| | **Host family** | Study ID + effect size ID + host species + host phylogeny | Host family | Culicidae<br>Drosophilidae |
| | **Symbiont type** | Study ID + effect size ID + host species + host phylogeny | Symbiont type | Natural<br>Introduced |
| | **Natural enemy group** | Study ID + effect size ID + host species + host phylogeny | Natural enemy group | Bacteria<br>Nematode<br>Parasitoid<br>Protozoan<br>Virus |
| | **Fitness measure** | Study ID + effect size ID + host species + host phylogeny | Fitness measure | Natural enemy load<br>Survival |
| | **Natural enemy killing strategy** | Study ID + effect size ID + host species + host phylogeny | Natural enemy killing strategy | Facultative lethal<br>Obligate lethal |

**Table S8.** Subgroup analysis: analyses performed only with Aphididae data. Multilevel meta-analytical models performed in this study, specifying their random and fixed factors (moderators) and moderator levels. Moderators "symbiont type" and "natural enemy group" were not included in the cost to the natural enemy data analysis because they presented only one moderator level ("natural" and "parasitoid", respectively).

| Data (Aphididae) | Model | Random factors | Fixed factors (moderators) | Moderator levels |
|---|---|---|---|---|
| **Protection** | **Overall** | Study ID + effect size ID + host species + host phylogeny | - | - |
| | **Symbiont type** | Study ID + effect size ID + host species + host phylogeny | Symbiont type | Natural<br>Introduced |
| | **Symbiont species** | Study ID + effect size ID + host species + host phylogeny | Symbiont species | *Hamiltonella*<br>*Hamiltonella* + *Regiella*<br>Other species<br>*Regiella*<br>*Regiella* + *Spiroplasma*<br>*Spiroplasma* |
| | **Natural enemy group** | Study ID + effect size ID + host species + host phylogeny | Natural enemy group | Fungus<br>Parasitoid |
| | **Fitness measure** | Study ID + effect size ID + host species + host phylogeny | Fitness measure | Fecundity<br>Survival<br>Uninfected hosts |
| | **Symbiont number** | Study ID + effect size ID + host species + host phylogeny | Symbiont number | Single<br>Coinfection |
| | **Natural enemy killing strategy** | Study ID + effect size ID + host species + host phylogeny | Natural enemy killing strategy | Obligate lethal<br>Facultative lethal |
| **Cost to hosts** | **Overall** | Study ID + effect size ID + host species + host phylogeny | - | - |

| | | | | |
|---|---|---|---|---|
| | **Symbiont type** | Study ID + effect size ID + host species + host phylogeny | Symbiont type | Natural<br>Introduced |
| | **Symbiont species** | Study ID + effect size ID + host species + host phylogeny | Symbiont species | *Hamiltonella*<br>*Hamiltonella* + *Rickettsia*<br>*Hamiltonella* + *Rickettsiella*<br>Other species<br>*Regiella*<br>*Regiella* + *Spiroplasma*<br>*Rickettsia*<br>*Rickettsia* + *Serratia*<br>*Rickettsiella*<br>*Ricketsiella* + *Serratia*<br>*Serratia*<br>*Spiroplasma* |
| | **Fitness measure** | Study ID + effect size ID + host species + host phylogeny | Fitness measure | Body size<br>Development time<br>Fecundity<br>Survival |
| | **Symbiont number** | Study ID + effect size ID + host species + host phylogeny | Symbiont number | Single<br>Coinfection |
| **Cost to the natural enemy** | **Overall** | Study ID + effect size ID + host species + host phylogeny | - | - |
| | **Symbiont species** | Study ID + effect size ID + host species + host phylogeny | Symbiont species | *Hamiltonella*<br>*Regiella* + *Spiroplasma*<br>*Spiroplasma* |
| | **Fitness measure** | Study ID + effect size ID + host species + host phylogeny | Fitness measure | Body size<br>Survival |
| | **Symbiont number** | Study ID + effect size ID + host species + host phylogeny | Symbiont number | Single<br>Coinfection |

**Table S9.** Main analysis: results of the meta-analyses on the costs and benefits of hosting defensive symbionts including symbiont number, natural enemy group and natural enemy killing strategy as moderators.

| Data | Model and moderator levels | k | m | Mean effect size | Lower 95% C.I. | Upper 95% C.I. | p-value | QM | p-value |
|---|---|---|---|---|---|---|---|---|---|
| **Protection** | **Symbiont number** | | | | | | | 33.08 | **<0.001** |
| | Single | 1170 | 97 | 0.568 | 0.328 | 0.809 | **<0.001** | | |
| | Coinfection | 104 | 12 | 0.986 | 0.642 | 1.331 | **<0.001** | | |
| | **Natural enemy group** | | | | | | | 41.88 | **<0.001** |
| | Bacteria | 20 | 5 | 0.105 | -0.678 | 0.889 | 0.792 | | |
| | Fungus | 120 | 17 | 0.935 | 0.581 | 1.288 | **<0.001** | | |
| | Nematode | 17 | 5 | 0.537 | -0.229 | 1.304 | 0.169 | | |
| | Parasitoid | 958 | 54 | 0.667 | 0.386 | 0.949 | **<0.001** | | |
| | Protozoan | 14 | 7 | -0.170 | -0.927 | 0.587 | 0.660 | | |
| | Virus | 145 | 19 | 0.731 | 0.320 | 1.142 | **<0.001** | | |
| | **Natural enemy killing strategy** | | | | | | | 11.14 | **0.004** |
| | Facultative lethal | 316 | 52 | 0.609 | 0.248 | 0.970 | **<0.001** | | |
| | Obligate lethal | 958 | 54 | 0.424 | 0.039 | 0.809 | **0.030** | | |
| **Cost to host** | **Symbiont number** | | | | | | | 0.65 | 0.721 |
| | Single | 1570 | 166 | -0.083 | -0.286 | 0.119 | 0.418 | | |
| | Coinfection | 181 | 28 | -0.077 | -0.348 | 0.193 | 0.575 | | |
| **Cost to natural enemy** | **Symbiont number** | | | | | | | 67.70 | **<0.001** |
| | Single | 348 | 55 | -0.968 | -1.319 | -0.616 | **<0.001** | | |
| | Coinfection | 52 | 4 | -2.443 | -3.031 | -1.855 | **<0.001** | | |

| | | | | | | | | |
|---|---|---|---|---|---|---|---|---|
| **Natural enemy group** | | | | | | | 43.85 | **<0.001** |
| Bacteria | 22 | 6 | -0.407 | -1.519 | 0.706 | 0.473 | | |
| Nematode | 19 | 4 | -2.433 | -3.682 | -1.184 | **<0.001** | | |
| Parasitoid | 198 | 19 | -1.263 | -1.846 | -0.679 | **<0.001** | | |
| Protozoan | 19 | 8 | -1.171 | -2.151 | -0.191 | **0.019** | | |
| Virus | 142 | 21 | -0.679 | -1.240 | -0.119 | **0.018** | | |
| **Natural enemy killing strategy** | | | | | | | 33.04 | **<0.001** |
| Facultative lethal | 202 | 38 | -0.928 | -1.373 | -0.483 | **<0.001** | | |
| Obligate lethal | 198 | 19 | -1.267 | -1.881 | -0.653 | **<0.001** | | |

**Table S10.** Subgroup analysis: results of the analyses including only effect sizes from hosts harboring bacterial symbiont *Wolbachia*.

| **Data (*Wolbachia* only)** | **Model and moderator levels** | **k** | **m** | **Mean effect size** | **Lower 95% C.I.** | **Upper 95% C.I.** | **p-value** | **QM** | **p-value** |
|---|---|---|---|---|---|---|---|---|---|
| **Protection** | **Overall** | 238 | 38 | 0.264 | -0.032 | 0.560 | 0.081 | | |
| | **Host family** | | | | | | | 2.62 | 0.453 |
| | Culicidae | 58 | 19 | 0.257 | -0.239 | 0.754 | 0.309 | | |
| | Drosophilidae | 161 | 16 | 0.257 | -0.196 | 0.710 | 0.267 | | |
| | Other families | 19 | 3 | 0.235 | -0.536 | 1.006 | 0.549 | | |
| | **Fitness measure** | | | | | | | 6.57 | 0.087 |
| | Fecundity | 11 | 3 | 0.169 | -0.534 | 0.873 | 0.637 | | |
| | Uninfected hosts | 42 | 11 | 0.689 | 0.152 | 1.228 | **0.012** | | |
| | Survival | 185 | 25 | 0.153 | -0.170 | 0.476 | 0.353 | | |
| | **Symbiont type** | | | | | | | 3.48 | 0.175 |
| | Natural | 151 | 25 | 0.242 | -0.067 | 0.552 | 0.125 | | |
| | Introduced | 87 | 15 | 0.348 | -0.061 | 0.756 | 0.095 | | |
| | **Natural enemy group** | | | | | | | 13.71 | **0.033** |
| | Bacteria | 14 | 2 | 0.498 | -0.350 | 1.346 | 0.250 | | |
| | Fungus | 13 | 6 | 0.108 | -0.482 | 0.699 | 0.718 | | |
| | Nematode | 9 | 4 | 0.303 | -0.428 | 1.034 | 0.417 | | |
| | Parasitoid | 47 | 3 | -0.323 | -1.023 | 0.376 | 0.364 | | |
| | Protozoan | 13 | 6 | -0.355 | -1.015 | 0.304 | 0.291 | | |
| | Virus | 142 | 18 | 0.538 | 0.177 | 0.900 | **0.004** | | |
| | **Natural enemy killing strategy** | | | | | | | 7.09 | **0.029** |
| | Facultative lethal | 191 | 36 | 0.308 | 0.016 | 0.599 | **0.038** | | |
| | Obligate lethal | 47 | 3 | -0.347 | -1.015 | 0.321 | 0.308 | | |
| **Cost to hosts** | **Overall** | 723 | 43 | 0.036 | -0.248 | 0.320 | 0.804 | | |
| | **Host family** | | | | | | | 1.14 | 0.767 |
| | Culicidae | 262 | 29 | -0.217 | -0.885 | 0.451 | 0.524 | | |
| | Drosophilidae | 248 | 32 | 0.261 | -0.346 | 0.868 | 0.399 | | |

| | | | | | | | | | |
|---|---|---|---|---|---|---|---|---|---|
| | Other families | 213 | 35 | 0.032 | -0.356 | 0.419 | 0.873 | | |
| | **Fitness measure** | | | | | | | 25.53 | **<0.001** |
| | Body size | 61 | 17 | 0.742 | 0.335 | 1.150 | **<0.001** | | |
| | Development time | 114 | 31 | 0.200 | -0.147 | 0.547 | 0.259 | | |
| | Fecundity | 199 | 74 | 0.013 | -0.291 | 0.317 | 0.934 | | |
| | Survival | 349 | 69 | -0.107 | -0.401 | 0.187 | 0.474 | | |
| | **Symbiont type** | | | | | | | 0.65 | 0.724 |
| | Natural | 436 | 72 | 0.061 | -0.224 | 0.347 | 0.673 | | |
| | Introduced | 287 | 30 | -0.080 | -0.484 | 0.324 | 0.697 | | |
| **Cost to natural enemy** | **Overall** | 223 | 36 | -0.612 | -1.834 | 0.609 | 0.326 | | |
| | **Host family** | | | | | | | 1.55 | 0.459 |
| | Culicidae | 101 | 25 | -1.181 | -3.039 | 0.676 | 0.213 | | |
| | Drosophilidae | 122 | 11 | -0.017 | -1.891 | 1.858 | 0.986 | | |
| | **Fitness measure** | | | | | | | 0.97 | 0.615 |
| | Natural enemy load | 186 | 33 | -0.618 | -1.849 | 0.612 | 0.325 | | |
| | Survival | 37 | 4 | -0.578 | -2.074 | 0.919 | 0.449 | | |
| | **Symbiont type** | | | | | | | 2.40 | 0.301 |
| | Natural | 98 | 19 | -0.503 | -1.611 | 0.605 | 0.374 | | |
| | Introduced | 125 | 21 | -0.822 | -1.967 | 0.323 | 0.160 | | |
| | **Natural enemy group** | | | | | | | 3.09 | 0.685 |
| | Bacteria | 15 | 3 | -0.295 | -2.026 | 1.436 | 0.738 | | |
| | Nematode | 15 | 3 | -1.494 | -3.216 | 0.229 | 0.089 | | |
| | Parasitoid | 33 | 3 | -0.401 | -2.128 | 1.326 | 0.649 | | |
| | Protozoan | 19 | 8 | -0.626 | -2.114 | 0.862 | 0.410 | | |
| | Virus | 141 | 20 | -0.612 | -1.865 | 0.641 | 0.338 | | |
| | **Natural enemy killing strategy** | | | | | | | 1.01 | 0.603 |
| | Facultative lethal | 190 | 33 | -0.633 | -1.867 | 0.602 | 0.315 | | |
| | Obligate lethal | 33 | 3 | -0.481 | -2.200 | 1.239 | 0.584 | | |

**Table S11.** Subgroup analysis: results of the analyses including only effect sizes from hosts from the Aphididae family. Moderators "symbiont type" and "natural enemy group" were not included in the cost to the natural enemy data analysis because they presented only one moderator level ("natural" and "parasitoid", respectively).

| **Data** (Aphids only) | **Model and moderator levels** | **k** | **m** | **Mean effect size** | **Lower 95% C.I.** | **Upper 95% C.I.** | **p-value** | **QM** | **p-value** |
|---|---|---|---|---|---|---|---|---|---|
| **Protection** | **Overall** | 881 | 54 | 0.872 | 0.586 | 1.159 | **<0.001** | | |
| | **Fitness measure** | | | | | | | 59.03 | **<0.001** |
| | Fecundity | 10 | 3 | -0.795 | -1.714 | 0.124 | 0.090 | | |
| | Uninfected hosts | 834 | 46 | 0.966 | 0.685 | 1.247 | **<0.001** | | |
| | Survival | 37 | 9 | 0.659 | 0.003 | 1.314 | **0.049** | | |
| | **Symbiont type** | | | | | | | 49.90 | **<0.001** |
| | Natural | 743 | 52 | 0.868 | 0.621 | 1.115 | **<0.001** | | |
| | Introduced | 138 | 5 | 1.013 | 0.502 | 1.524 | **<0.001** | | |
| | **Symbiont species** | | | | | | | 38.24 | **<0.001** |
| | *Hamiltonella* | 687 | 40 | 0.699 | 0.327 | 1.071 | **<0.001** | | |
| | *Hamiltonella* + *Regiella* | 13 | 2 | 1.664 | 0.835 | 2.494 | **<0.001** | | |
| | Other | 50 | 14 | 1.363 | 0.856 | 1.869 | **<0.001** | | |
| | *Regiella* | 52 | 14 | 0.996 | 0.508 | 1.483 | **<0.001** | | |
| | *Regiella* + *Spiroplasma* | 22 | 2 | 0.533 | -0.374 | 1.441 | 0.250 | | |
| | *Spiroplasma* | 57 | 5 | 0.645 | -0.090 | 1.380 | 0.085 | | |
| | **Symbiont number** | | | | | | | 33.68 | **<0.001** |
| | Single | 822 | 52 | 0.827 | 0.502 | 1.152 | **<0.001** | | |
| | Coinfection | 59 | 10 | 1.305 | 0.833 | 1.776 | **<0.001** | | |
| | **Natural enemy group** | | | | | | | 26.80 | **<0.001** |
| | Fungus | 109 | 12 | 1.143 | 0.696 | 1.589 | **<0.001** | | |
| | Parasitoid | 772 | 47 | 0.795 | 0.438 | 1.151 | **<0.001** | | |

| | | | | | | | | | |
|---|---|---|---|---|---|---|---|---|---|
| | **Natural enemy killing strategy** | | | | | | | 26.80 | **<0.001** |
| | Facultative lethal | 109 | 12 | 1.143 | 0.696 | 1.589 | **<0.001** | | |
| | Obligate lethal | 772 | 47 | 0.795 | 0.438 | 1.151 | **<0.001** | | |
| **Cost to host** | **Overall** | 828 | 11 | -0.456 | -0.858 | -0.053 | **0.027** | | |
| | **Fitness measure** | | | | | | | 105.96 | **<0.001** |
| | Body size | 52 | 13 | 0.182 | -0.340 | 0.704 | 0.494 | | |
| | Development time | 163 | 23 | 0.253 | -0.191 | 0.698 | 0.264 | | |
| | Fecundity | 414 | 49 | -0.621 | -1.046 | -0.196 | **0.004** | | |
| | Survival | 199 | 28 | -0.830 | -1.267 | -0.393 | **<0.001** | | |
| | **Symbiont type** | | | | | | | 4.90 | **0.086** |
| | Natural | 662 | 48 | -0.461 | -0.870 | -0.052 | **0.027** | | |
| | Introduced | 166 | 12 | -0.424 | -0.942 | 0.095 | 0.109 | | |
| | **Symbiont species** | | | | | | | 38.02 | **<0.001** |
| | *Hamiltonella* | 319 | 33 | -0.690 | -1.131 | -0.248 | **0.002** | | |
| | *Hamiltonella + Rickettsia* | 11 | 2 | -0.166 | -0.972 | 0.641 | 0.687 | | |
| | *Hamiltonella + Rickettsiella* | 10 | 2 | -1.206 | -2.163 | -0.249 | **0.014** | | |
| | Other species | 32 | 7 | -0.560 | -1.179 | 0.059 | 0.076 | | |
| | *Regiella* | 141 | 20 | -0.156 | -0.638 | 0.326 | 0.527 | | |
| | *Regiella + Spiroplasma* | 24 | 3 | -0.420 | -1.200 | 0.360 | 0.291 | | |
| | *Rickettsia* | 79 | 7 | 0.204 | -0.321 | 0.729 | 0.447 | | |
| | *Rickettsia + Serratia* | 18 | 1 | 0.270 | -0.528 | 1.068 | 0.507 | | |
| | *Rickettsiella* | 18 | 3 | -0.493 | -1.350 | 0.365 | 0.260 | | |
| | *Rickettsiella + Serratia* | 27 | 1 | 0.087 | -0.558 | 0.732 | 0.792 | | |
| | *Serratia* | 81 | 12 | 0.001 | -0.551 | 0.552 | 0.998 | | |
| | *Spiroplasma* | 68 | 5 | -0.449 | -0.994 | 0.096 | 0.106 | | |
| | **Symbiont number** | | | | | | | 5.36 | 0.069 |
| | Single | 714 | 51 | -0.462 | -0.861 | -0.062 | **0.023** | | |
| | Coinfection | 114 | 12 | -0.389 | -0.851 | 0.074 | 0.100 | | |
| **Cost to natural enemy** | **Overall** | 56 | 7 | -0.446 | -0.998 | 0.105 | 0.113 | | |

| | | | | | | | | |
|---|---|---|---|---|---|---|---|---|
| **Fitness measure** | | | | | | | 2.81 | 0.246 |
| Body size | 10 | 2 | -0.316 | -1.091 | 0.459 | 0.425 | | |
| Survival | 46 | 6 | -0.486 | -1.056 | 0.084 | 0.094 | | |
| **Symbiont species** | | | | | | | 3.04 | 0.386 |
| *Hamiltonella* | 20 | 6 | -0.473 | -1.140 | 0.194 | 0.165 | | |
| *Regiella + Spiroplasma* | 21 | 1 | -0.405 | -1.958 | 1.148 | 0.609 | | |
| *Spiroplasma* | 15 | 1 | -0.213 | -1.777 | 1.351 | 0.789 | | |
| **Symbiont number** | | | | | | | 3.34 | 0.188 |
| Single | 35 | 7 | -0.428 | -0.984 | 0.128 | 0.131 | | |
| Coinfection | 21 | 1 | -0.610 | -1.265 | 0.044 | 0.068 | | |

**Table S12**. Main analysis: results of the heterogeneity test ($I^2$) for each meta-analytical model. For each model, we quantified the total amount of heterogeneity ($I^2$) and the percentage of contribution of each random effect for the variance in the models: study, effect size, host phylogeny and host species.

| Data | Model | $I^2$ total (%) | $I^2$ Study (%) | $I^2$ Effect size (%) | $I^2$ Host phylogeny (%) | $I^2$ Host species (%) |
|---|---|---|---|---|---|---|
| Protection | Overall | 97.08 | 28.60 | 61.98 | 0.87 | 5.61 |
| | Host family | 97.09 | 29.03 | 61.69 | 4.92E-06 | 6.36 |
| | Fitness measure | 97.11 | 32.48 | 60.73 | 1.54E-06 | 3.89 |
| | Symbiont type | 97.09 | 28.66 | 61.77 | 1.12 | 5.54 |
| | Symbiont species | 97.10 | 30.05 | 60.58 | 1.77E-06 | 6.46 |
| | Symbiont number | 97.07 | 28.89 | 61.66 | 0.22 | 6.30 |
| | Natural enemy group | 96.99 | 28.77 | 63.55 | 1.02E-05 | 4.67 |
| | Natural enemy killing strategy | 97.16 | 27.04 | 60.07 | 3.65 | 6.40 |
| Cost to hosts | Overall | 98.21 | 19.63 | 61.53 | 3.70E-06 | 17.05 |
| | Host family | 98.19 | 20.42 | 62.49 | 4.33E-06 | 15.28 |
| | Fitness measure | 98.16 | 20.46 | 58.77 | 0.55 | 18.39 |
| | Symbiont type | 98.20 | 19.91 | 61.98 | 9.30E-06 | 16.31 |
| | Symbiont species | 98.20 | 20.05 | 61.88 | 1.39E-06 | 16.28 |
| | Symbiont number | 98.21 | 19.65 | 61.55 | 5.69E-06 | 17.01 |
| Cost to natural enemy | Overall | 97.58 | 41.01 | 56.56 | 4.28E-08 | 1.30E-06 |
| | Host family | 98.17 | 30.04 | 42.26 | 25.86 | 5.86E-07 |
| | Fitness measure | 97.52 | 39.44 | 58.08 | 3.44E-07 | 8.95E-07 |
| | Symbiont type | 97.58 | 40.32 | 56.42 | 2.34E-06 | 8.35E-01 |
| | Symbiont species | 97.36 | 34.78 | 36.75 | 22.30 | 3.53 |
| | Symbiont number | 97.40 | 43.11 | 54.29 | 3.95E-10 | 5.15E-06 |
| | Natural enemy group | 97.45 | 37.88 | 59.57 | 9.93E-08 | 8.79E-07 |
| | Natural enemy killing strategy | 97.59 | 41.33 | 56.26 | 2.06E-07 | 9.98E-07 |

**Table S13.** Subgroup analysis: results of the heterogeneity test ($I^2$) for each meta-analytical model. For each model, we quantified the total amount of heterogeneity ($I^2$) and the percentage of contribution of each random effect for the variance in the models: study, effect size, host phylogeny and host species.

| Data | Model | $I^2$ total (%) | $I^2$ Study (%) | $I^2$ Effect size (%) | $I^2$ Host phylogeny (%) | $I^2$ Host species (%) |
|---|---|---|---|---|---|---|
| Protection (*Wolbachia* only) | Overall | 96.91 | 24.29 | 55.83 | 2.77E-07 | 16.78 |
| | Host family | 97.03 | 25.57 | 53.56 | 1.16E-06 | 17.89 |
| | Fitness measure | 96.87 | 24.45 | 56.23 | 9.88E-07 | 16.19 |
| | Symbiont type | 96.90 | 24.50 | 56.29 | 4.29E-07 | 16.11 |
| | Natural enemy group | 96.74 | 22.52 | 57.88 | 5.28E-07 | 16.34 |
| | Natural enemy killing strategy | 96.75 | 18.40 | 59.10 | 4.94E-07 | 19.25 |
| Cost to host (*Wolbachia* only) | Overall | 98.95 | 11.47 | 60.94 | 1.32E-06 | 26.54 |
| | Host family | 98.98 | 10.92 | 59.25 | 2.55E-06 | 28.81 |
| | Fitness measure | 98.89 | 12.57 | 61.44 | 1.47E-06 | 24.89 |
| | Symbiont type | 98.93 | 12.57 | 62.08 | 9.91E-07 | 24.28 |
| Cost to natural enemy (*Wolbachia* only) | Overall | 97.68 | 25.16 | 41.65 | 30.87 | 1.05E-06 |
| | Host family | 97.81 | 24.00 | 39.14 | 34.67 | 6.48E-07 |
| | Fitness measure | 97.70 | 25.71 | 41.44 | 30.55 | 1.46E-06 |
| | Symbiont type | 97.49 | 26.71 | 45.41 | 25.37 | 1.08E-06 |
| | Natural enemy group | 97.69 | 26.87 | 41.78 | 29.05 | 7.30E-07 |
| | Natural enemy killing strategy | 97.71 | 26.15 | 41.22 | 30.34 | 1.27E-06 |
| Protection (Aphids only) | Overall | 97.32 | 33.15 | 62.84 | 2.76E-04 | 1.33 |
| | Fitness measure | 97.46 | 38.91 | 58.55 | 2.51E-07 | 2.51E-06 |
| | Symbiont type | 97.33 | 34.58 | 62.73 | 2.00E-03 | 0.02 |
| | Symbiont species | 97.4 | 33.16 | 60.08 | 4.47E-06 | 4.16 |
| | Symbiont number | 97.33 | 32.16 | 62.27 | 3.75E-06 | 2.9 |
| | Natural enemy group | 97.31 | 29.7 | 62.95 | 3.02E-06 | 4.66 |

| | | | | | | |
|---|---|---|---|---|---|---|
| | Natural enemy killing strategy | 97.31 | 29.7 | 62.95 | 3.02E-06 | 4.66 |
| Cost to host (Aphids only) | Overall | 94.38 | 23.43 | 60.42 | 2.62E-06 | 10.53 |
| | Fitness measure | 93.94 | 23.88 | 56.76 | 7.72E-06 | 13.30 |
| | Symbiont type | 94.41 | 23.65 | 60.14 | 2.67E-06 | 10.62 |
| | Symbiont species | 94.46 | 24.10 | 58.60 | 2.93E-06 | 11.76 |
| | Symbiont number | 94.38 | 23.73 | 60.52 | 2.45E-06 | 10.13 |
| Cost to natural enemy (Aphids only) | Overall | 81.66 | 68.18 | 13.48 | 3.36E-07 | 6.69E-08 |
| | Fitness measure | 81.13 | 66.11 | 15.02 | 3.84E-07 | 4.42E-08 |
| | Symbiont species | 84.49 | 72.91 | 11.58 | 2.09E-07 | 3.61E-08 |
| | Symbiont number | 81.91 | 68.07 | 13.84 | 1.72E-07 | 4.04E-08 |

**Table S14.** Main analysis: results of the Egger's regression test for publication bias for each meta-analytical model. Models with p-values in bold represent the models with potential publication bias.

| Data | Model | Intercept | t | p-value |
|---|---|---|---|---|
| Protection | Overall | 0.905 | 20.650 | **<0.001** |
| | Host family | 0.778 | 18.230 | **<0.001** |
| | Fitness measure | 0.733 | 17.460 | **<0.001** |
| | Symbiont type | 0.909 | 20.710 | **<0.001** |
| | Symbiont species | 0.850 | 19.610 | **<0.001** |
| | Symbiont number | 0.890 | 20.120 | **<0.001** |
| | Natural enemy group | 0.812 | 18.580 | **<0.001** |
| | Natural enemy killing strategy | 1.029 | 23.300 | **<0.001** |
| Cost to host | Overall | -0.167 | -3.659 | **<0.001** |
| | Host family | -0.034 | -0.761 | 0.447 |
| | Fitness measure | -0.194 | -4.328 | **<0.001** |
| | Symbiont type | -0.158 | -3.463 | **<0.001** |
| | Symbiont species | -0.147 | -3.255 | **0.001** |
| | Symbiont number | -0.167 | -3.662 | **<0.001** |
| Cost to natural enemy | Overall | -0.337 | -3.068 | **0.002** |
| | Host family | -0.179 | -1.639 | 0.102 |
| | Fitness measure | -0.315 | -2.917 | **0.004** |
| | Symbiont type | -0.346 | -3.122 | **0.002** |
| | Symbiont species | -0.088 | -0.823 | 0.411 |
| | Symbiont number | -0.197 | -1.828 | 0.068 |
| | Natural enemy group | -0.336 | -3.162 | **0.002** |
| | Natural enemy killing strategy | -0.294 | -2.703 | **0.007** |

**Table S15.** Subset analysis: results of the Egger's regression test for publication bias for each meta-analytical model. Models with p-values in bold represent the models with potential publication bias.

| Data | Model | Intercept | t | p-value |
|---|---|---|---|---|
| Protection (*Wolbachia* only) | Overall | 0.543 | 6.145 | **<0.001** |
| | Host family | 0.552 | 6.254 | **<0.001** |
| | Fitness measure | 0.468 | 5.384 | **<0.001** |
| | Symbiont type | 0.526 | 6.011 | **<0.001** |
| | Natural enemy group | 0.593 | 7.133 | **<0.001** |
| | Natural enemy killing strategy | 0.716 | 8.457 | **<0.001** |
| Cost to host (*Wolbachia* only) | Overall | -0.056 | -0.976 | 0.329 |
| | Host family | -0.059 | -1.016 | 0.310 |
| | Fitness measure | -0.077 | -1.363 | 0.173 |
| | Symbiont type | -0.028 | -0.496 | 0.620 |
| Cost to natural enemy (*Wolbachia* only) | Overall | -0.162 | -1.129 | 0.260 |
| | Host family | -0.241 | -1.694 | 0.092 |
| | Fitness measure | -0.162 | -1.124 | 0.262 |
| | Symbiont type | -0.087 | -0.607 | 0.544 |
| | Natural enemy group | -0.145 | -1.021 | 0.309 |
| | Natural enemy killing strategy | -0.159 | -1.105 | 0.270 |
| Protection (Aphids only) | Overall | 0.920 | 17.270 | **<0.001** |
| | Fitness measure | 0.863 | 16.430 | **<0.001** |
| | Symbiont type | 0.899 | 16.780 | **<0.001** |
| | Symbiont species | 1.038 | 19.390 | **<0.001** |
| | Symbiont number | 0.937 | 17.470 | **<0.001** |
| | Natural enemy group | 0.955 | 17.770 | **<0.001** |
| | Natural enemy killing strategy | 0.955 | 17.770 | **<0.001** |
| Cost to host (Aphids only) | Overall | -0.305 | -3.193 | **0.001** |
| | Fitness measure | -0.297 | -3.177 | **0.002** |
| | Symbiont type | -0.307 | -3.220 | **0.001** |

| | | | | |
|---|---|---|---|---|
| | Symbiont species | -0.377 | -4.001 | **<0.001** |
| | Symbiont number | -0.309 | -3.235 | **0.001** |
| Cost to natural enemy (Aphids only) | Overall | 0.243 | 2.100 | **0.040** |
| | Fitness measure | 0.268 | 2.410 | **0.019** |
| | Symbiont species | 0.140 | 1.184 | 0.242 |
| | Symbiont number | 0.314 | 2.816 | **0.007** |

**Table S16.** Results of the sensitivity analysis for the overall models, showing the maximum Cook's distance and DFBETAs values for each model.

| Analysis | Data | Cook's distance (D) | DFBETAS |
|---|---|---|---|
| **Main analysis** | Protection | 0.049 | 0.078 |
| | Cost to host | 0.048 | 0.219 |
| | Cost to natural enemy | 0.080 | 0.110 |
| **Sub-analysis** | ***Wolbachia* only** | | |
| | Protection | 0.122 | 0.074 |
| | Cost to host | 0.056 | 0.238 |
| | Cost to natural enemy | 0.018 | 0.081 |
| | **Aphids only** | | |
| | Protection | 0.023 | 0.117 |
| | Cost to host | 0.022 | 0.149 |
| | Cost to natural enemy | 0.716 | 0.165 |
| **Trade-off meta-regression** | - | 1.247 | 0.397 |
